# Autoimmunity and clinical pathology amelioration in SLE by Dexamethasone primed Mesenchymal Stem Cell derived conditioned media

**DOI:** 10.1101/2024.06.29.601320

**Authors:** Khushbu Priya, Sonali Rawat, Doli Das, Manaswi Chaubey, Hiral Thacker, Kiran Giri, Shambhavi Singh, Madhukar Rai, Sujata Mohanty, Geeta Rai

**Affiliations:** Department of Molecular and Human Genetics, Institute of Science, Banaras Hindu University, Varanasi-221005, India; Stem Cell Facility, DBT-Centre of Excellence for Stem Cell Research, AIIMS, New Delhi-110029, India; Department of Medicine, Institute of Medical Sciences, Banaras Hindu University, Varanasi-221005, India; Department of Pharmacology, Institute of Medical Sciences, Banaras Hindu University Varanasi-221005, India; DY Patil University, Navi Mumbai-400706, India

## Abstract

This study proposes a novel approach, utilizing cell-free dexamethasone (Dex) primed Whartons jelly mesenchymal stem cells derived conditioned media (DW), offering improved efficacy, simplicity, and alternative medicine for addressing complications associated with systemic lupus erythematosus (SLE). This study explores the immunomodulatory effects of DW treatment on immune cell populations in SLE patients and a pristane-induced lupus (PIL) mouse model. DW induces significant expansion of Tregs, Bregs, suppressing Th17, double-negative T cells, and inflammatory neutrophils through modulating IL-10 and IL-17A production. Comparisons with the standard drug hydroxychloroquine reveal similar effects, suggesting TGF beta; pathway mediation in DW’s actions. Compared with the immunosuppressive drug Dex, DW better attenuated autoantibody production, increased anti-inflammatory cytokines and maintained a balanced Th17/Treg ratio. In the preclinical in vivo studies, DW exhibits therapeutic efficacy, reducing mortality, preventing proteinuria, and reversing limb inflammation, seizure and alopecia. Organ specific evaluations using advanced live imaging or histopathological analysis highlighted DW’s protective effects on kidneys, liver, lungs, heart, and spleen. This provided insight into the immunomodulatory benefits of DW at various levels and suggested that it could be a potential therapeutic avenue for managing complications related to SLE.

## INTRODUCTION

Systemic lupus erythematosus (SLE) is an autoimmune disorder characterized by the production of antinuclear antibodies that target the host nuclear antigens. This condition affects various organs, causing chronic inflammation, tissue damage, and diverse clinical manifestations. Despite genetic and environmental factors contributing to its onset, SLE predominantly affects women, impacting fertility and increasing the risk of complications like miscarriage and premature birth. Advancements in treatment notwithstanding, SLE remains incurable, with existing therapies, such as corticosteroids and immunosuppressants, often bearing severe side effects. In this context, stem cell therapy, particularly utilizing mesenchymal stem cells (MSCs), has emerged as a promising approach due to its immunoregulatory properties and potential for tissue repair. Over the last decade, MSCs have been reported to possess marked immune-regulatory effects against autoimmune disorders and have been shown to suppress T/B cell proliferation (Blanco et al., 2016, Fontaine et al., 2016).

However, challenges in standardizing protocols and addressing MSC heterogeneity hinder clinical trial success, resulting in low cell survival, migration, and differentiation rates of transplanted stem cells. The harsh microenvironment of damaged tissues further contributes significantly to poor outcomes in stem cell therapy, prompting the exploration of preconditioning methods. Effective preconditioning of MSCs with cytokines (Duijvestein et al., 2011), hypoxia (Sun et al., 2015), oxidative stress (Numasawa et al., 2011), or chemicals (Li et al., 2016) is emerging as a promising approach to enhance MSC function post-transplantation. The primary objective of MSC priming is to prepare cells for challenging post-transplantation environments and facilitate desired therapeutic effects. Nonetheless, concerns persist regarding potential low cell survival, migration, and differentiation rates.

Recognizing these challenges evaluating the immunomodulatory potential a cell-free preconditioned/primed MSC-derived conditioned media (MSC-CM) in SLE seems imperative. The secreted factors or the secretome from the MSCs growing in the culture medium is harvested as CM. Being cell-free eliminates inconsistencies associated with MSCs, and simplified storage and transportation compared to MSCs positions CM as a novel therapeutic option with improved efficacy in mitigating disease severity. Various inducers, including pharmacological, biological, and physical stimuli, can be employed for preconditioning, and known to positively impacting cell survival (Zhang et al., 2001), proliferation and differentiation (Lu et al., 2017), immunomodulation (Song et al., 2017), paracrine signaling (Chang et al., 2013), and angiogenesis (Lan et al., 2015). This preconditioned or primed MSC-CM offers an alternative to cell-based therapeutic strategies, leveraging the concept of paracrine signaling.

A new approach of preconditioning MSC-CM with Dexamethasone (Dex) was used and reported earlier by members of our research group (Rawat et al., 2021). This study extends the use of Dex preconditioned MSC-CM for assessing its immunomodulatory role in SLE patients. Dex, a synthetic glucocorticoid with anti-inflammatory and immunosuppressant properties, mimics the body’s anti-inflammatory hormones. While its exact mechanism of action is not fully understood, animal studies suggest that Dex acts by binding to the glucocorticoid receptor (Whelan and Apfel, 2013). It is well-tolerated and effective in treating inflammation in various parts of the body, autoimmune diseases (Paolino et al., 2017; Bertsias et al., 2012), skin conditions (Deckers et al., 2018), asthma (Sellers et al., 2022), and other lung conditions (Ray et al., 1996).

The treatment of SLE has been a challenging subject in medical research. However, a therapeutic approach that combines immunomodulation with pathological suppression can be a promising intervention. In particular, to achieve this approach Dex-primed Wharton’s jelly (WJ) derived MSCs CM (DW) garnered significant attention due to its potential to lower the use of glucocorticoids, which are known to lead to morbidity owing to their adverse side effects. Therefore, such an approach represents an attractive strategy for the treatment of SLE. By reducing the use of glucocorticoids, patients can experience better quality of life, and the overall morbidity rates are expected to decrease. Thus, we used DW as a strategy to explore it *in vitro* and in preclinical settings.

With the growing incidence of autoimmune diseases, particularly SLE, this study focuses on the exploration for novel therapeutic approach through DW. A comprehensive assessment of the immunomodulatory effects of DW treatment was conducted employing a pristane-induced lupus (PIL) mouse model and peripheral blood mononuclear cells (PBMCs) from patients with SLE. Examining diverse patient demographics, including age and autoantibody profiles, the study revealed significant expansions of different regulatory T cell subtypes assessed phenotypically defined as (i) CD4^+^ CD25^+^ CD127^−^, (ii) CD4^+^ CD25^+^ CD127^−^ IL-10^+^, (iii) CD4^+^ IL-10^+^, (iv) CD4^+^ FOXP3^+^ and (v) CD4^+^ FOXP3^+^ IL-10^+^, and Bregs population including i) CD19^+^ CD24^+^ CD27^+^, ii) CD19^+^ CD24^+^ CD27^+^ IL-10^+^ alongside the concurrent suppression of pathogenic CD4^+^ IL-17A^+^ TH17 immune cells and Double negative (DN) CD3^+^ CD4^−^ CD8^−^ T cells following DW treatment. Comparative analyses with hydroxychloroquine (HCQ), a standard treatment for SLE, indicated similar immunomodulatory effects of DW, and the combination of the two exhibited better immunomodulation. Involvement of the TGF-β pathway in DW’s mode of action was substantiated by inhibition studies, elucidating the mechanistic underpinnings of DW’s effects. Lupus model was developed using female BALB/c mice. *In vivo* studies were performed and the mice were assessed for improvement of disease specific histopathology of kidney, spleen, lungs, liver and heart, autoantibody production, proteinuria, body weights, limb inflammation, adaptive immune system components; cytokines and TGF-β. DW further demonstrated clinical benefits, such as reversal of symptoms of seizure, alopecia and inflammation. *In vivo* PIL mouse model studies have highlighted therapeutic efficacy of DW as a potential disease-modifying intervention. Photoacoustic imaging (PAI) provided valuable insights into DW’s protective effects on vital organs including the kidneys, liver, and heart of lupus mice.

These results suggest DW to be a promising and comprehensive cell free biological therapeutic option for SLE treatment and associated autoimmune conditions. The diverse advantages observed using DW emphasize the need for clinical trials, to confirm its effectiveness and safety in humans.

## RESULTS

### Isolation of WJ-MSCs and preparation of Dex primed MSCs CM

We acquired WJ tissue samples from mothers undergoing either normal vaginal or caesarean delivery after obtaining informed consent. WJ-MSCs were isolated, cultured, and analyzed for proliferation, immunophenotypes, and differentiation capacities. Their trilineage differentiation capability towards osteogenic, chondrogenic and adipogenic cells lineages was evaluated. Isolated WJ-MSCs formed a homogenous monolayer of adherent, spindle-shaped cells and exhibited proliferation capacity of WJ-MSCs (Supplemental Figure 1, A and B). Immunophenotyping confirmed the purity of the isolated cells, and their surface antigen expression revealed the phenotypic properties of WJ-MSCs and surface marker profiling showed >95% positivity for CD105, CD90, CD73, CD29, and HLA-Class I, and negativity for HLA-class II and CD34/45 (Supplemental Figure 1, C and D). Isolated WJ-MSCs successfully underwent trilineage differentiation into osteocytes, adipocytes, and chondrocytes as evidenced by alizarin red staining, oil red ‘o’ staining and alcian blue staining respectively (Supplemental Figure 1 E). After the initial characterization the WJ-MSCs were primed with Dex for 24 h following which the serum-free culture supernatant was collected (DW) and preserved for subsequent use. Unprimed culture supernatant (W) was collected for comparative studies.

### Characteristics of study subjects

The characteristics of the patients with SLE enrolled in this study are included in Table 1. Total of 74 patients (males=4, females=70) were enrolled in the study (age range-15 to 60 years). Overall, 20 % of patients with SLE had active disease. The patients were on corticosteroids or immunosuppressive drugs, like HCQ (35), omnacortil (18) or both (17).

**Table 1.** Summary of patient’s Characteristics.

| Description |  | No. of Patients |
| --- | --- | --- |
| Total number (n) |  | 74 |
| Gender | Male | 4 |
|  | Female | 70 |
| Age | Median | 27.5 |
|  | Range | 60 – 15 |
| Autoantibodies | ANA | 72 |
|  | Anti-dsDNA | 49 |
|  | Anti-ENA | 8 |
|  | Anti-dsDNA + Anti-ENA | 17 |
| Positive for Coomb's test |  | 7 |
| Elevated RA factor |  | 8 |
| Condition of Raynaud phenomenon |  | 7 |
| Clinical manifestation | Lupus nephritis | 21 |
|  | Systemic sclerosis | 9 |
|  | Hemolytic anemia | 13 |
|  | Hypothyroidism | 11 |
|  | Hepatomegaly | 10 |
|  | Splenomegaly | 12 |
|  | Scleroderma | 2 |
|  | Koch's lung | 5 |
|  | Cardiomegaly with pericardial effusion | 6 |
|  | Seizure disorder | 4 |
|  | Mixed connective tissue disease | 11 |
|  | Alopecia | 9 |
|  | CNS lupus | 3 |
|  | Polyarticular pain | 8 |
|  | Oral ulcer | 16 |
|  | Photosensitivity | 14 |
|  | Malar rash | 7 |
| Medication | HCQ | 35 |
|  | Omnacortil | 18 |
|  | HCQ + Omnacortil | 17 |
|  | MMF | 8 |
|  | Rituximab | 1 |

### DW treatment promoted regulatory cells expansion, suppressed Th17, DN T cells, and neutrophils by upregulating IL-10 and downregulating IL-17A

#### Tregs

Treg lymphocytes are key cells that control the autoimmunity process by maintaining immune tolerance and secreting various immunosuppressive and anti-inflammatory cytokines (Sakaguchi et al., 1995). Treg deficiencies have been linked to immunological aberrations observed in SLE and other autoimmune diseases (Pan et al., 2020). Different populations of Tregs exhibit multiple phenotypic features and variable markers. We assessed five different Treg subtypes, phenotypically defined as (i) CD4^+^ CD25^+^ CD127^−^, (ii) CD4^+^ CD25^+^ CD127^−^ IL-10^+^, (iii) CD4^+^ IL-10^+^, (iv) CD4^+^ FOXP3^+^ and (v) CD4^+^ FOXP3^+^ IL-10^+^ by flow cytometry in the isolated PBMCs of SLE patients. The frequency of all evaluated Tregs significantly increased after 24 h of DW treatment compared to that of untreated control or W treated PBMCs (Figure 1 A).

**Figure 1:**
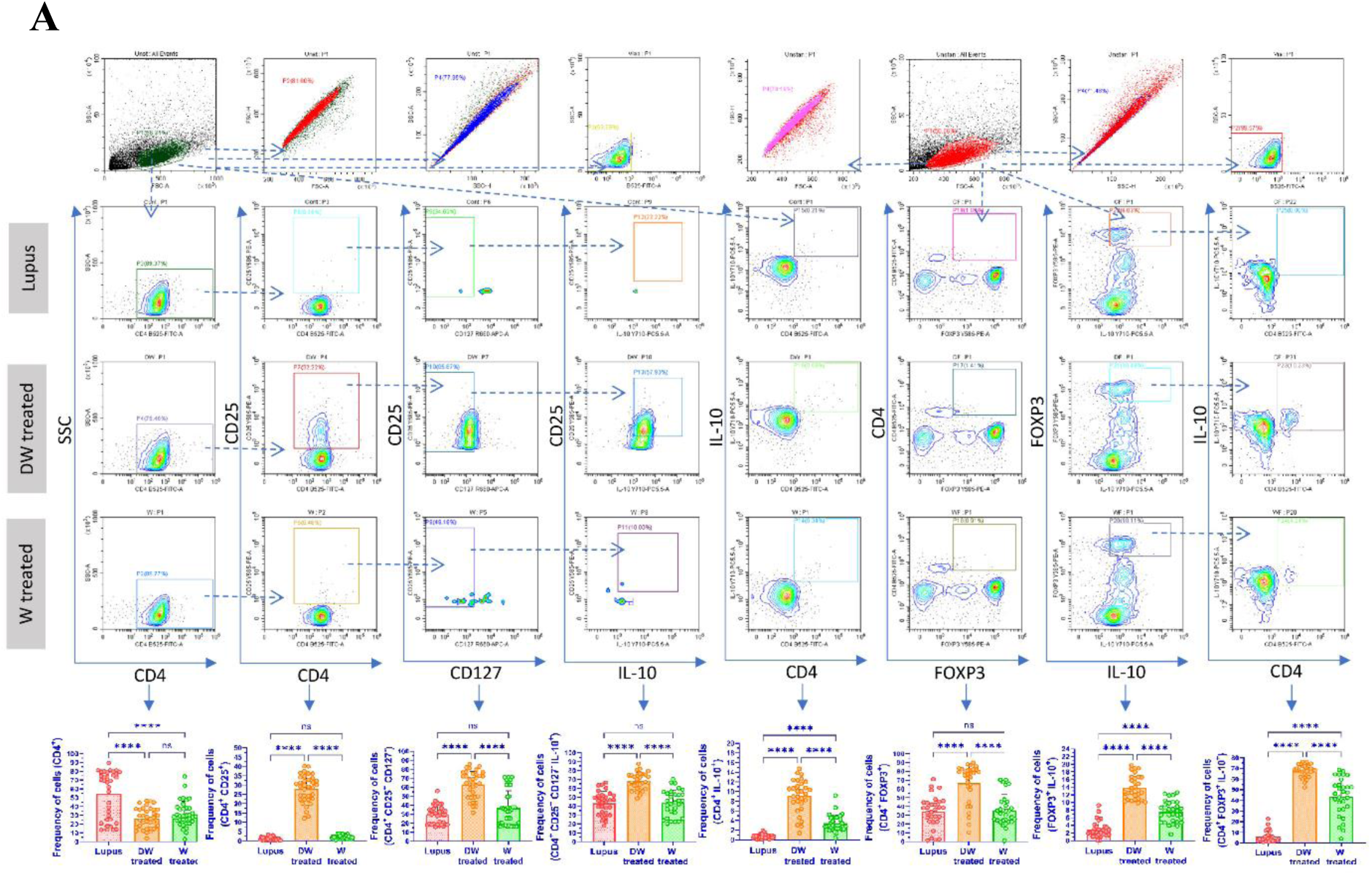
DW treatment promoted Tregs and Bregs expansion, suppressed Th17, DN T cells, and Inflammatory neutrophils by upregulating IL-10 and downregulating IL-17A. (A) Isolated PBMCs were stimulated and treated with DW and W for 24 hours and checked for the frequency of CD4^+^ CD25^+^, CD4^+^ CD25^+^ CD127^−^, CD4^+^ CD25^+^ CD127^−^ IL-10^+^, CD4^+^ IL-10^+^, CD4^+^ FOXP3^+^ and CD4^+^ FOXP3^+^ IL-10^+^ Tregs. The contour plots depict findings from a sample of 30 patients with SLE, and corresponding bar graphs are presented with a sample size of 30. Error bars show mean ± SD. p values indicate significant changes as follows: non-significant (ns) p > 0.05, *P < 0.05, **p < 0.01, ***P < 0.001 and ****P < 0.0001; One-way ANOVA.

#### Bregs

Bregs are also crucial for regulating autoimmune responses and primarily exert their effects by releasing immunosuppressive cytokine IL-10, which prevents immune cells from differentiating into effector/memory Breg subsets (Knippenberg et al., 2011). We examined two memory Breg populations (i) CD19^+^ CD24^+^ CD27^+^ and (ii) CD19^+^ CD24^+^ CD27^+^ IL-10. Further, the B cells of patients with SLE express PD-1 and correlate with the clinical progression of the disease (Figure 1 B). Hence, we also measured the frequency of (i) CD19^+^ IL-10^+^ B cells and (ii) CD19^+^ PD-1^+^ using SLE patients’ blood PBMCs and studied the effect of DW and W treatment on these cells. Untreated cells served as lupus controls. After 24 h of DW treatment, there was a noticeably decrease in frequency of CD19^+^ cells, and increase in frequency CD19^+^ CD24^+^ CD27^+^, CD19^+^ CD24^+^ CD27^+^ IL-10^+^ Breg memory cells and CD19^+^ IL-10^+^ Bregs compared with untreated cells (lupus). However, the CD19^+^ PD-1^+^ B cell populations were considerably lower in DW-treated cells as compared to untreated cells (Figure 1 B).

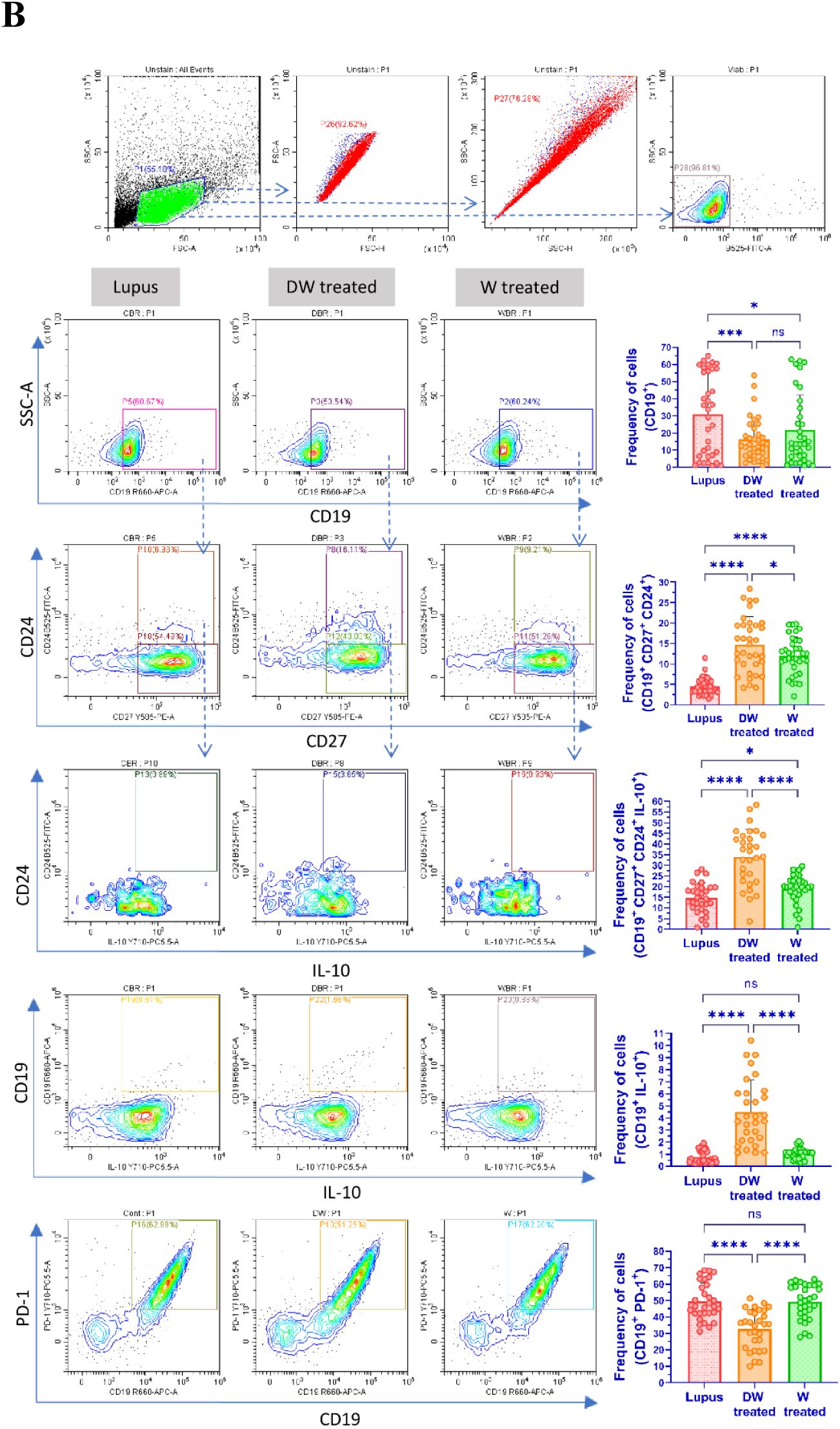
(B) Isolated PBMCs were stimulated and treated with DW and W for 24 hours and checked for the frequency of CD19^+^ B cells, IL-10-producing CD19^+^ B cells, PD-1 positive CD19^+^ B cells and Breg (CD19^+^ CD24^+^ CD27^+^). The contour plots depict findings from a sample of 30 patients with SLE, and corresponding bar graphs are presented with a sample size of 30. Error bars show mean ± SD. p values indicate significant changes as follows: non-significant (ns) p > 0.05, *P < 0.05, **p < 0.01, ***P < 0.001 and ****P < 0.0001; One-way ANOVA.

#### Th17 cells

T cell populations are altered in lupus, leaning more towards the inflammatory Th17 phenotype. There is an increase in the proportion of circulating Th17 cells in SLE patients. The imbalance between the two T cell subsets (Th17 and Tregs) that has been observed throughout the course of the disease as an intriguing problem to be addressed in the pathogenesis of SLE. The patients’ PBMCs were analyzed for the Th17 cell response. Additionally, CD4^+^ T cells bearing PD-1 accumulate at sites of inflammation in a variety of human autoimmune disorders, as evidenced by synovitis in rheumatoid arthritis and sialoadenitis in Sjogren’s syndrome (Hatachi et al., 2003). PD-1 plays an important role in the pathogenesis of SLE (Nishimura et al., 1999). Hence, we examined the effect of DW and W treatments on the differentiation of Th17 cells from CD4^+^ T cells and the frequency of Th17 and CD4^+^ T cells expressing PD-1 *in vitro* in PBMCs from SLE patients. Inflammatory CD4^+^ cells, expressing both IL-17A and PD-1, and Th17 cells expressing PD-1 were significantly reduced with DW treatment (Figure 1 C).

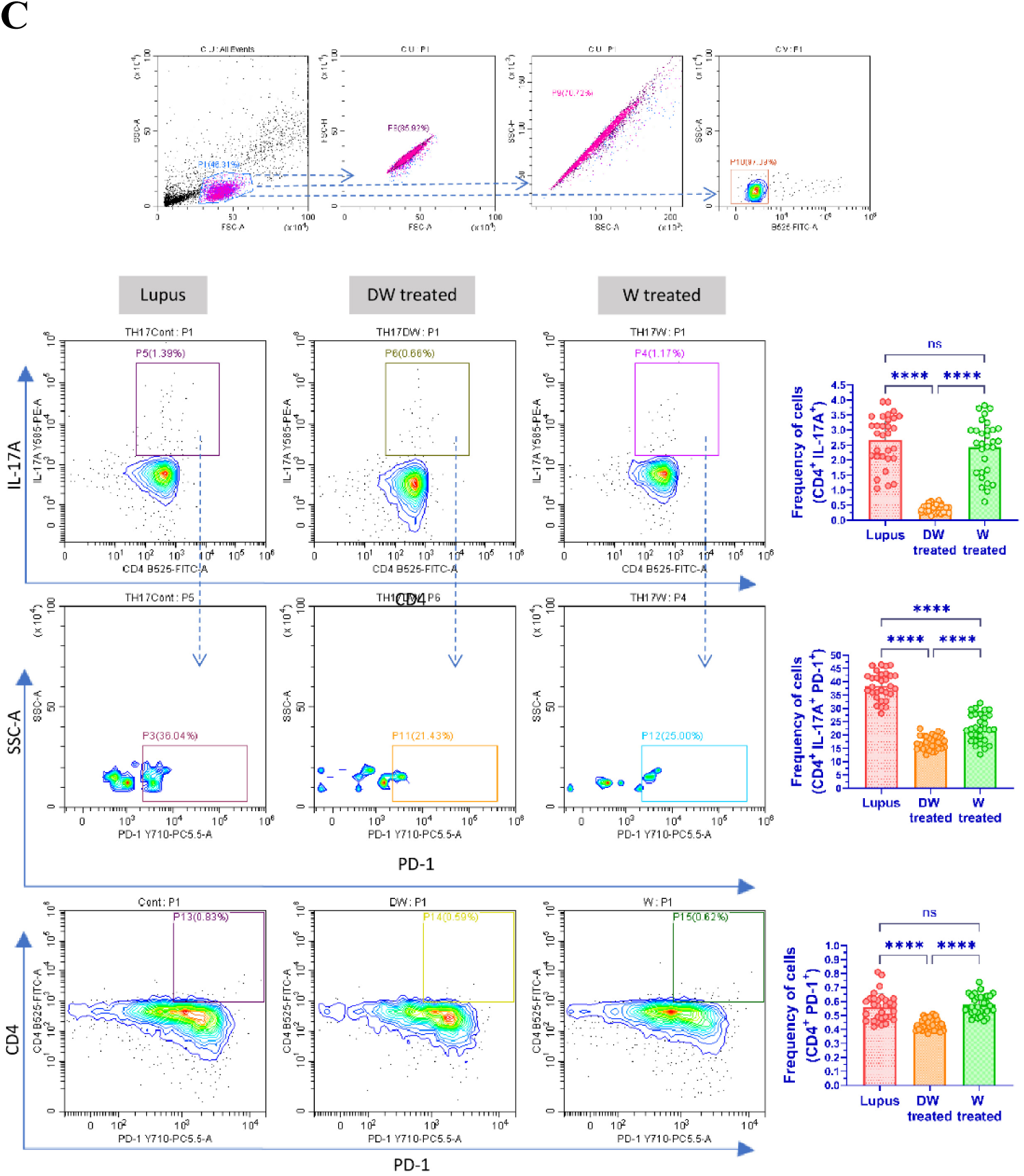
(C) Isolated PBMCs were stimulated and treated with DW and W for 24 hours and checked for the frequency of CD3^+^CD4^−^CD8^−^ DN population of T cells. The contour plots depict findings from a sample of 30 patients with SLE, and corresponding bar graphs are presented with a sample size of 30. Error bars show mean ± SD. p values indicate significant changes as follows: non-significant (ns) p > 0.05, *P < 0.05, **p < 0.01, ***P < 0.001 and ****P < 0.0001; One-way ANOVA.

#### DN T cells

In SLE patients and murine models of the disease, there is a significant increase in the CD3^+^ CD4^−^ CD8^−^ (DN) population of T cells, which normally accounts for less than 5% of the circulating CD3^+^ T cell population and is a key source of IL-17 (Tarbox et al, 2014). In our study, we found that DW treatment had a suppressive effect on the population of DN T cells. (Figure 1 D).

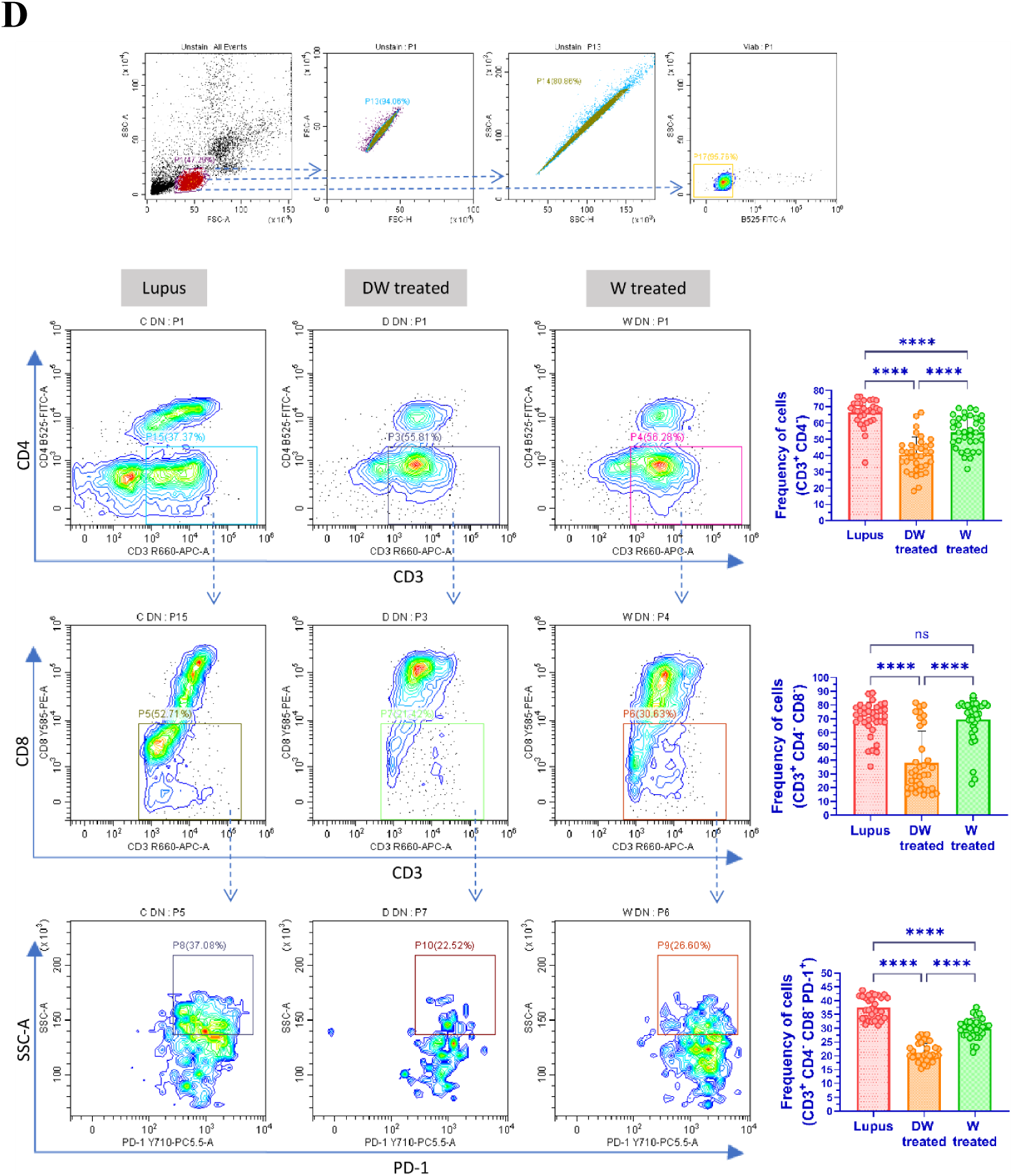
(D) Isolated PBMCs were stimulated and treated with DW and W for 24 hours and checked for the frequency of inflammatory CD4^+^ cells, expressing both IL-17A and PD-1. The contour plots depict findings from a sample of 30 patients with SLE, and corresponding bar graphs are presented with a sample size of 30. Error bars show mean ± SD. p values indicate significant changes as follows: non-significant (ns) p > 0.05, *P < 0.05, **p < 0.01, ***P < 0.001 and ****P < 0.0001; One-way ANOVA.

#### Inflammatory neutrophils

Compared to the healthy individuals, SLE patients’ neutrophils show increase in expression of PD1, PD-L1, Tim-3, CD40, and TIGIT and a notable rise in the prevalence of PD-L1-expressing neutrophils (Luo et al., 2016). This increased occurrence of CD15^+^ CD3^−^ PD-L1^+^ neutrophils aligns with the severity and disease activity in SLE patients (Luo et al., 2016). We found that treatment with DW considerably reduced the frequency of CD15^+^ CD3^−^ and CD15^+^ CD3^−^ PD-1^+^ neutrophils (Figure 1 E).

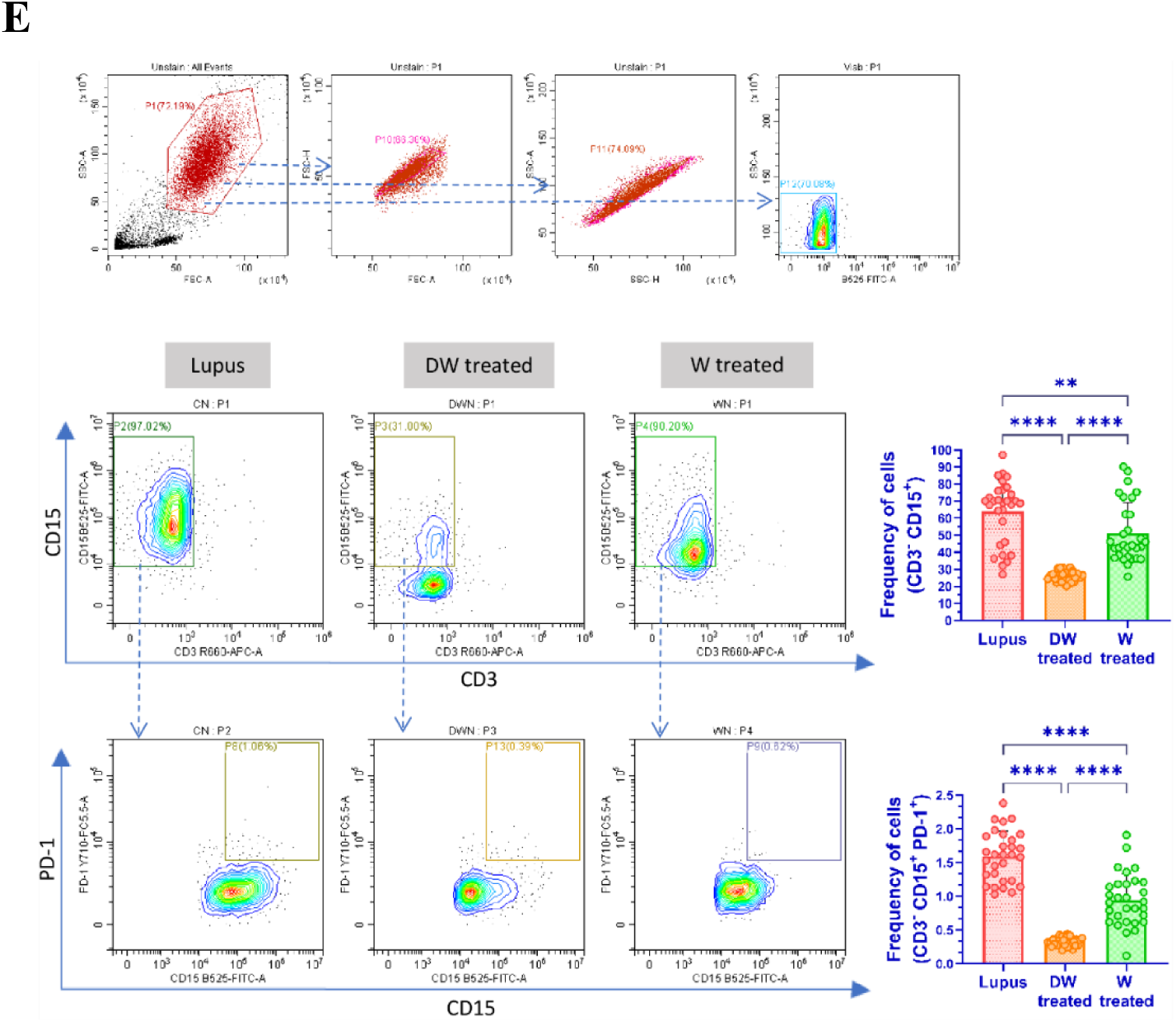
(E) Isolated PBMCs were stimulated and treated with DW and W for 24 hours and checked for the frequency of CD15^+^ CD3^−^ neutrophils and CD15^+^ CD3^−^ PD-1^+^ cells. The contour plots depict findings from a sample of 30 patients with SLE, and corresponding bar graphs are presented with a sample size of 30. Error bars show mean ± SD. p values indicate significant changes as follows: non-significant (ns) p > 0.05, *P < 0.05, **p < 0.01, ***P < 0.001 and ****P < 0.0001; One-way ANOVA.

Correlation analysis has elucidated that DW treatment has established a positive correlation between various subtypes of Bregs and Tregs. Additionally, there was a notable negative correlation observed with TH17, DN T cells and inflammatory neutrophils (Supplemental Figure 2). Consequently, the administration of DW facilitated the achievement of homeostasis *in vitro* in the SLE patients.

### Immunomodulation by DW treatment is similar to that of standard drug HCQ

HCQ is a standard therapy, which is currently the major long-term treatment recommended for all patients with SLE barring limitations or side effects. It reduces SLE activity, particularly in mild and moderate illness, avoid disease flare-ups, and reduce the long-term requirement for glucocorticoids (Bensreti et al., 2022). We therefore intended to compare the immunomodulatory effects of DW with that of the HCQ and Dex drug. We compared the effect of HCQ, Dex drug, DW and W treatments on the frequency of Tregs (CD4^+^ CD25^+^ CD127^−^ IL-10^+^), (CD4^+^ IL-17A^+^) Th17, DN T (CD3^+^ CD4^−^ CD8^−^) cells, Bregs (CD19^+^ CD24^+^ CD27^+^ IL-10^+^) and neutrophils (CD15^+^ CD3^−^). There was a significant increase in the number of Tregs (Figure 2 A) and Bregs (Figure 2 B), reduction in Th17 (Figure 2 C), DN T (Figure 2 D) cells and neutrophils (Figure 2 E) 24 h post-treatment with HCQ, DW, and W compared to the untreated group. Interestingly, the changes in frequency of all these cells were similar with both DW and HCQ-treatment but not with Dex drug and W treatment. Further, the combination of DW+HCQ yielded significantly better suppression of TH17 cells and neutrophils than the DW alone (Figure 2 C and E).

**Figure 2:**
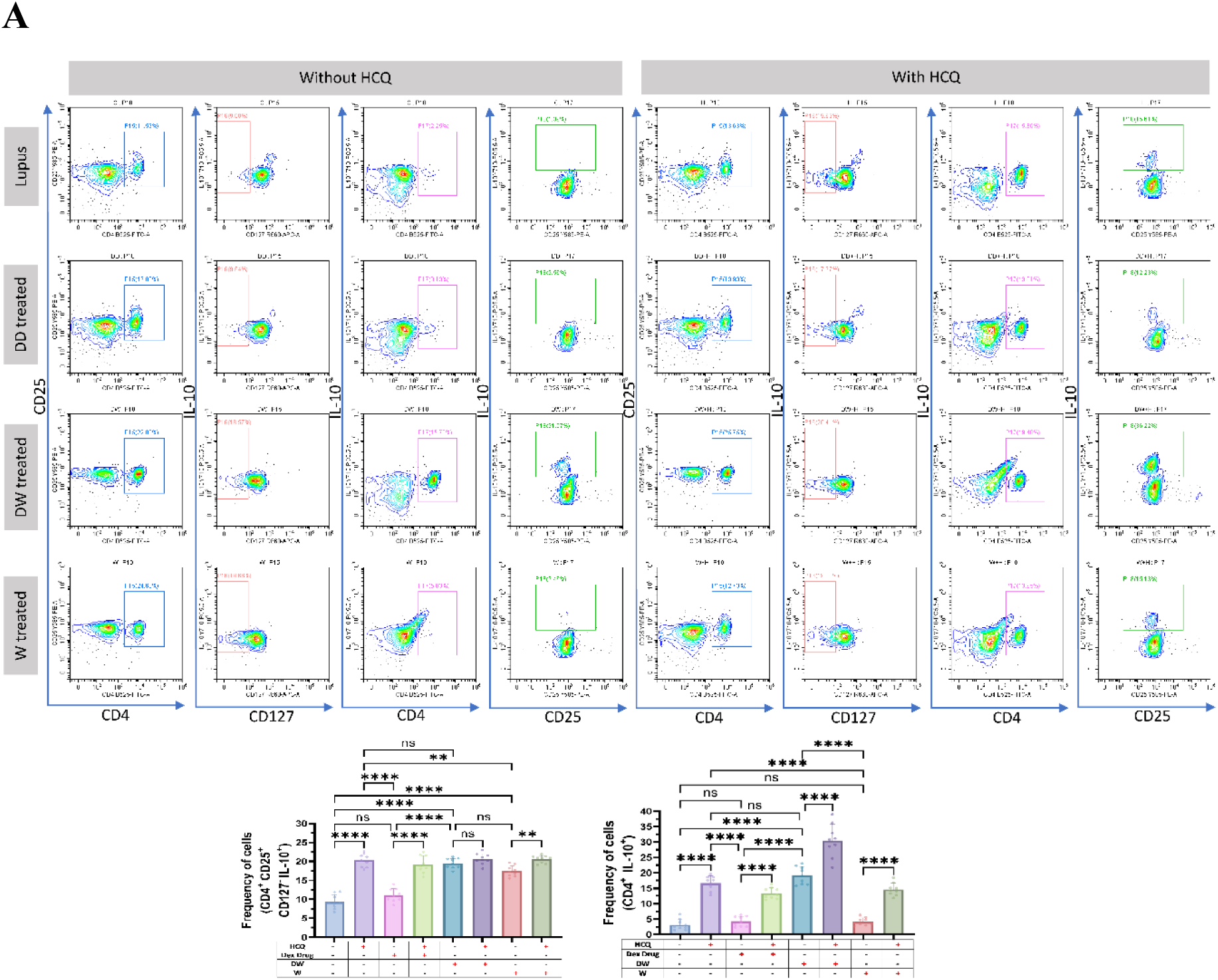
DW treatment has a similar immunomodulatory impact as HCQ as both of them promoted Tregs and Bregs expansion, suppressed Th17, DN T cells, and Inflammatory neutrophils by upregulating IL-10 and downregulating IL-17A. (A) Isolated PBMCs were stimulated and treated with HCQ, Dex drug, DW and W for 24 hours and checked for the frequency of CD4^+^ CD25^+^ CD127^−^ IL-10^+^ and CD4^+^ IL-10^+^ FOXP3^+^ Tregs. The contour plots depict findings from a sample of 9 patients with SLE, and corresponding bar graphs are presented with a sample size of 9. Error bars show mean ± SD. p values indicate significant changes as follows: non-significant (ns) p > 0.05, *P < 0.05, **p < 0.01, ***P < 0.001 and ****P < 0.0001; One-way ANOVA.

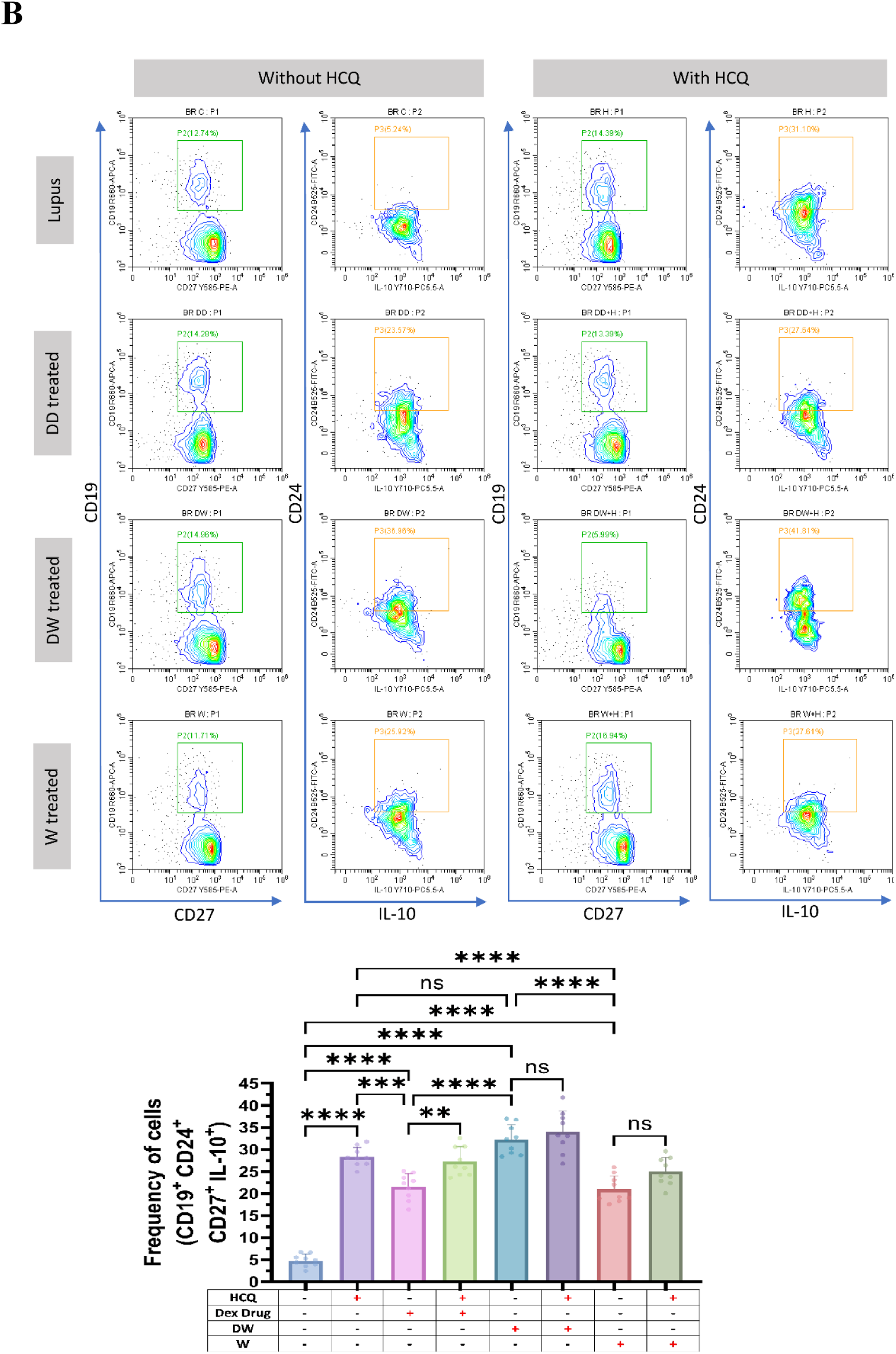
(B) Isolated PBMCs were stimulated and treated with HCQ, Dex drug, DW and W for 24 hours and checked for the frequency of inflammatory CD4^+^ cells, expressing IL-17A. The contour plots depict findings from a sample of 9 patients with SLE, and corresponding bar graphs are presented with a sample size of 9. Error bars show mean ± SD. p values indicate significant changes as follows: non-significant (ns) p > 0.05, *P < 0.05, **p < 0.01, ***P < 0.001 and ****P < 0.0001; One-way ANOVA.

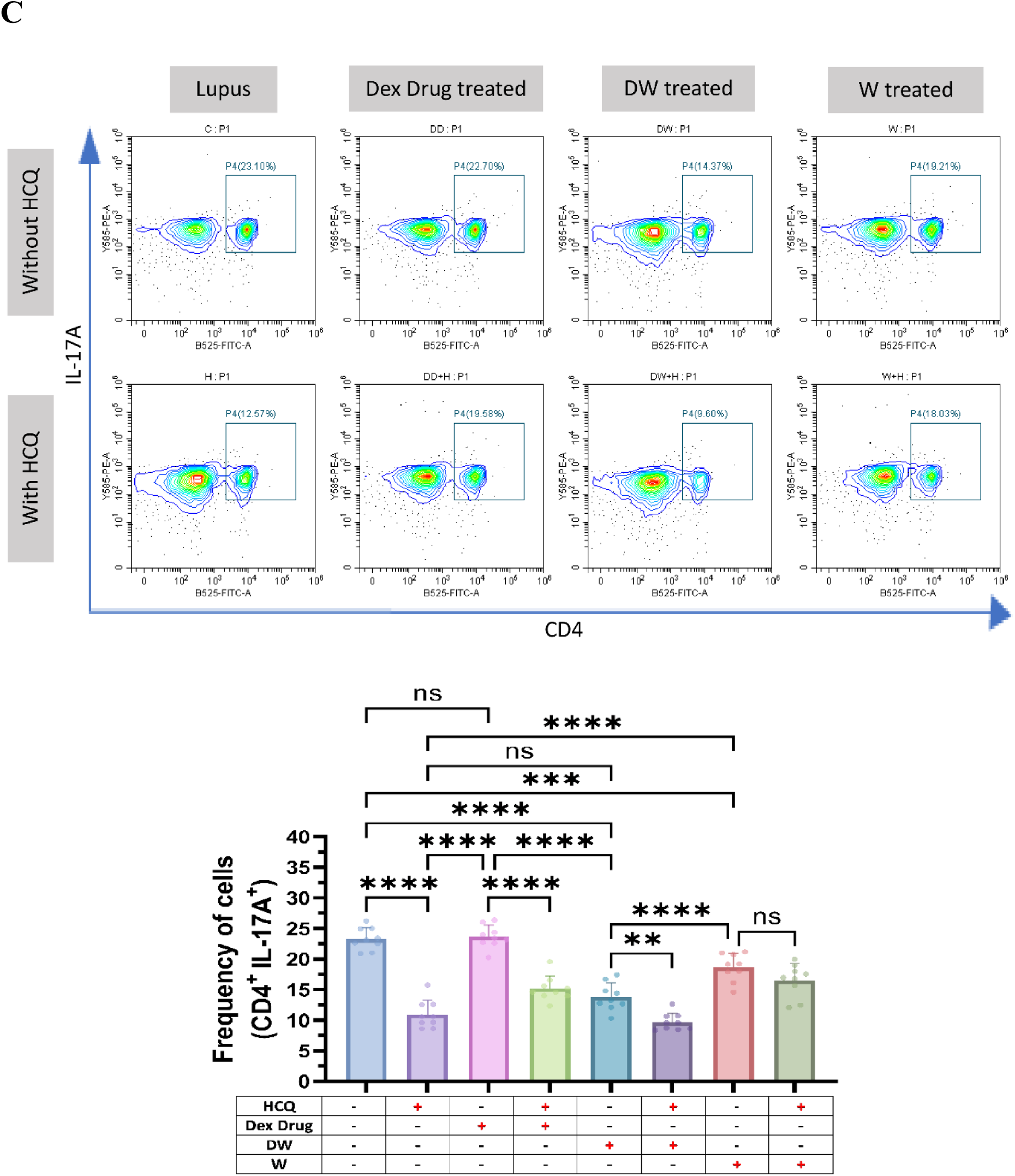
(C) Isolated PBMCs were stimulated and treated with HCQ, Dex drug, DW and W for 24 hours and checked for the frequency of CD3^+^CD4^−^CD8^−^ DN population of T cells. The contour plots depict findings from a sample of 9 patients with SLE, and corresponding bar graphs are presented with a sample size of 9. Error bars show mean ± SD. p values indicate significant changes as follows: non-significant (ns) p > 0.05, *P < 0.05, **p < 0.01, ***P < 0.001 and ****P < 0.0001; One-way ANOVA.

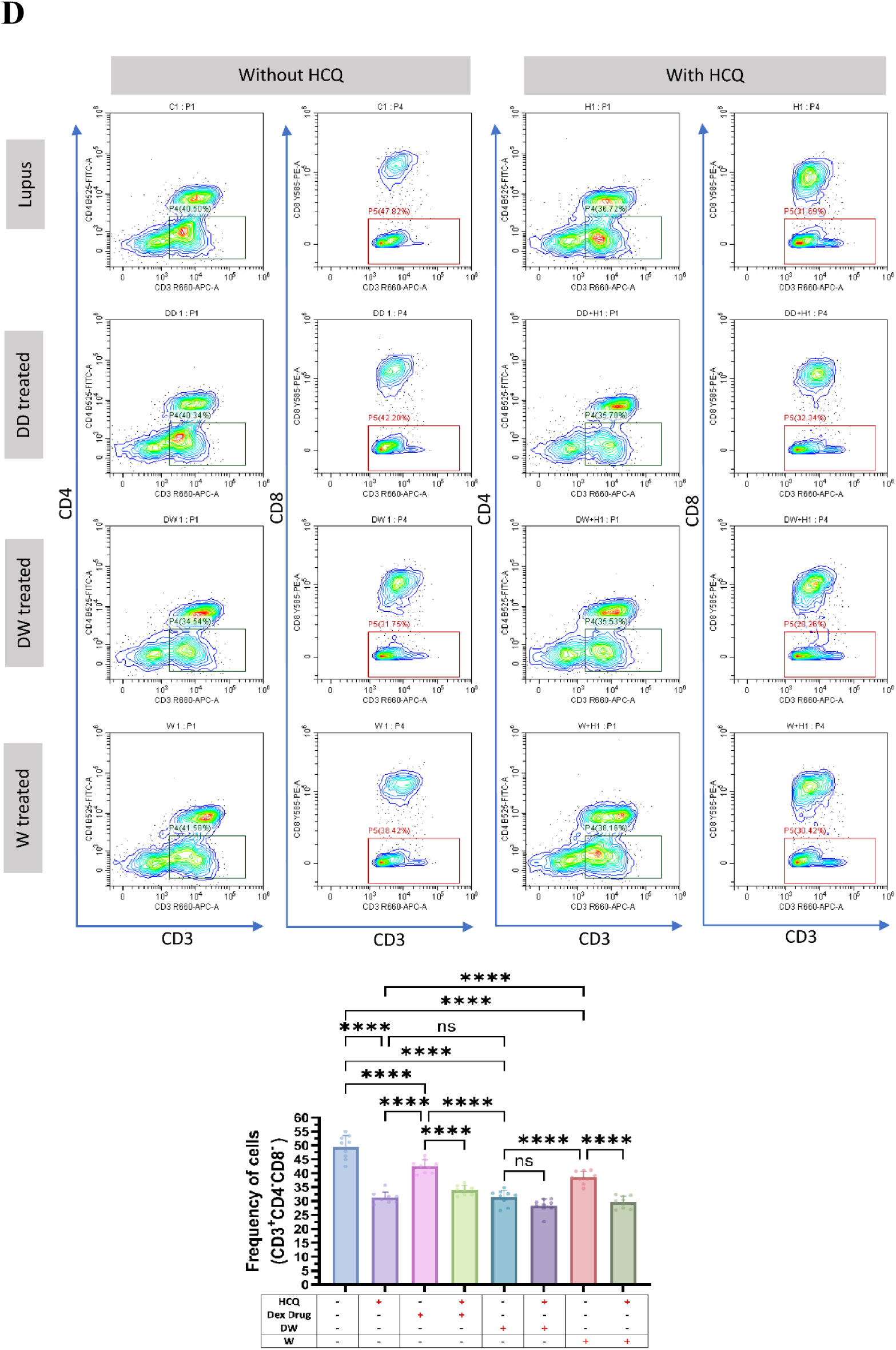
(D) Isolated PBMCs were stimulated and treated with HCQ, Dex drug, DW and W for 24 hours and checked for the frequency of frequency of CD19^+^ CD24^+^ CD27^+^ IL-10^+^ Breg. The contour plots depict findings from a sample of 9 patients with SLE, and corresponding bar graphs are presented with a sample size of 9. Error bars show mean ± SD. p values indicate significant changes as follows: non-significant (ns) p > 0.05, *P < 0.05, **p < 0.01, ***P < 0.001 and ****P < 0.0001; One-way ANOVA.

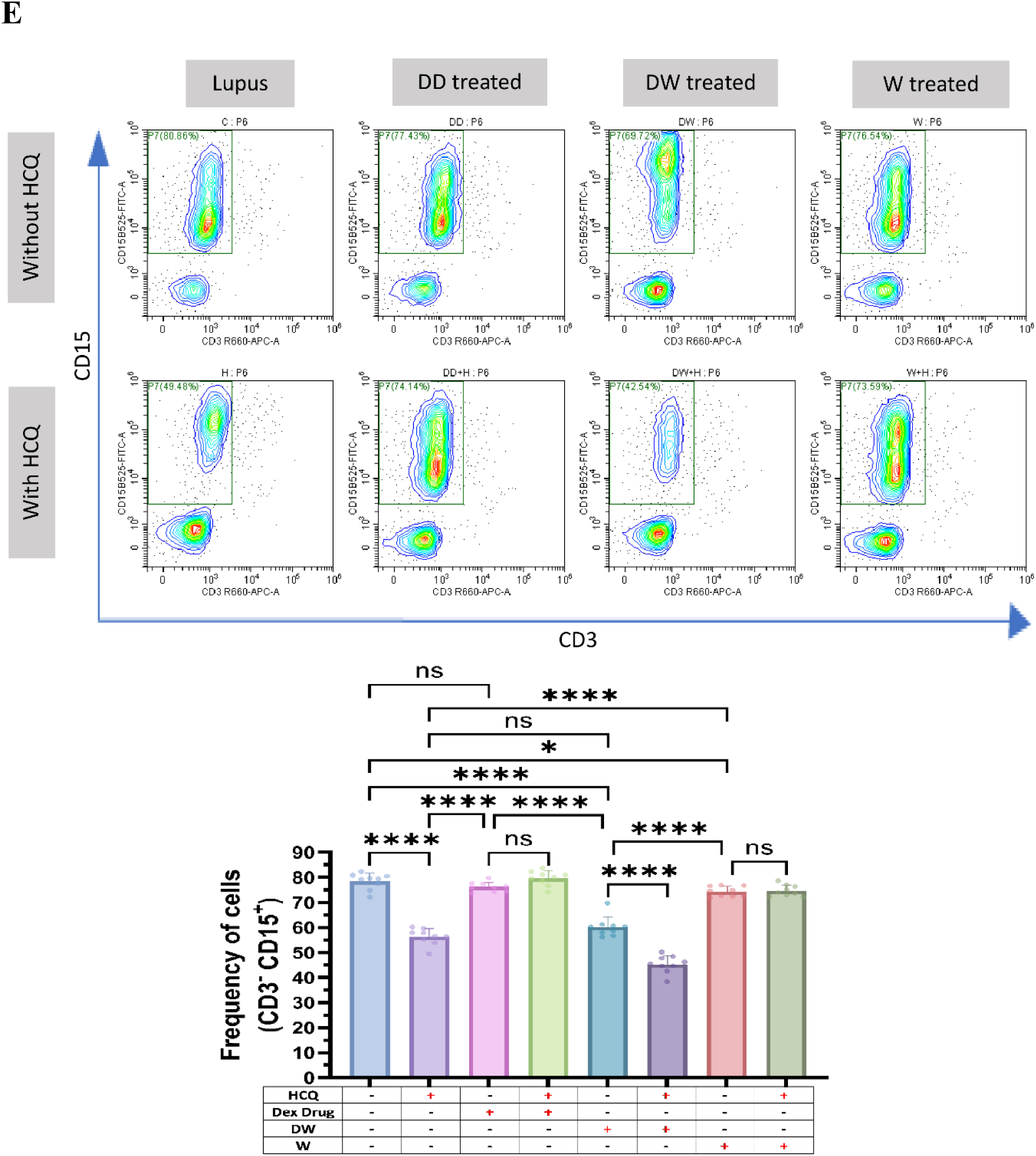
(E) Isolated PBMCs were stimulated and treated with HCQ, Dex drug, DW and W for 24 hours and checked for the frequency of CD15^+^ CD3^−^ inflammatory neutrophils. The contour plots depict findings from a sample of 9 patients with SLE, and corresponding bar graphs are presented with a sample size of 9. Error bars show mean ± SD. p values indicate significant changes as follows: non-significant (ns) p > 0.05, *P < 0.05, **p < 0.01, ***P < 0.001 and ****P < 0.0001; One-way ANOVA.

### Immunomodulatory activity of DW is mediated through TGF-β pathway

HCQ is known to work by inhibiting the release of pro-inflammatory cytokines, including IFN, TNF, and IL-12, increasing the release of anti-inflammatory cytokines like TGF-β1 and by promoting Treg differentiation and activity (Jin et al., 2022). HCQ is also known to restore the Th17/Treg balance, leading to significantly lower expression of IL-17 in Th17 cells and higher levels of Foxp3 and TGF-β (An et al., 2017). TGF-β plays a significant role in the regulation of autoimmunity by Breg and Treg cell development (Chen et al., 2003). To investigate whether DW mediated immunomodulation of Tregs and Bregs is also through TGF-β we used SB431542, an inhibitor of TGF-β activity (Cook et al., 2021) for inhibition assays. We observed that Tregs (Figure 3 A), Bregs (Figure 3 B) significantly decreased and Th17 (Figure 3 C), DN T cells (Figure 3 D), neutrophils (Figure 3 E) significantly increased when SB431542 was added to the DW treated culture for 24 h. This reversal of disease associated cell types was not as remarkable in other treatment groups.

**Figure 3:**
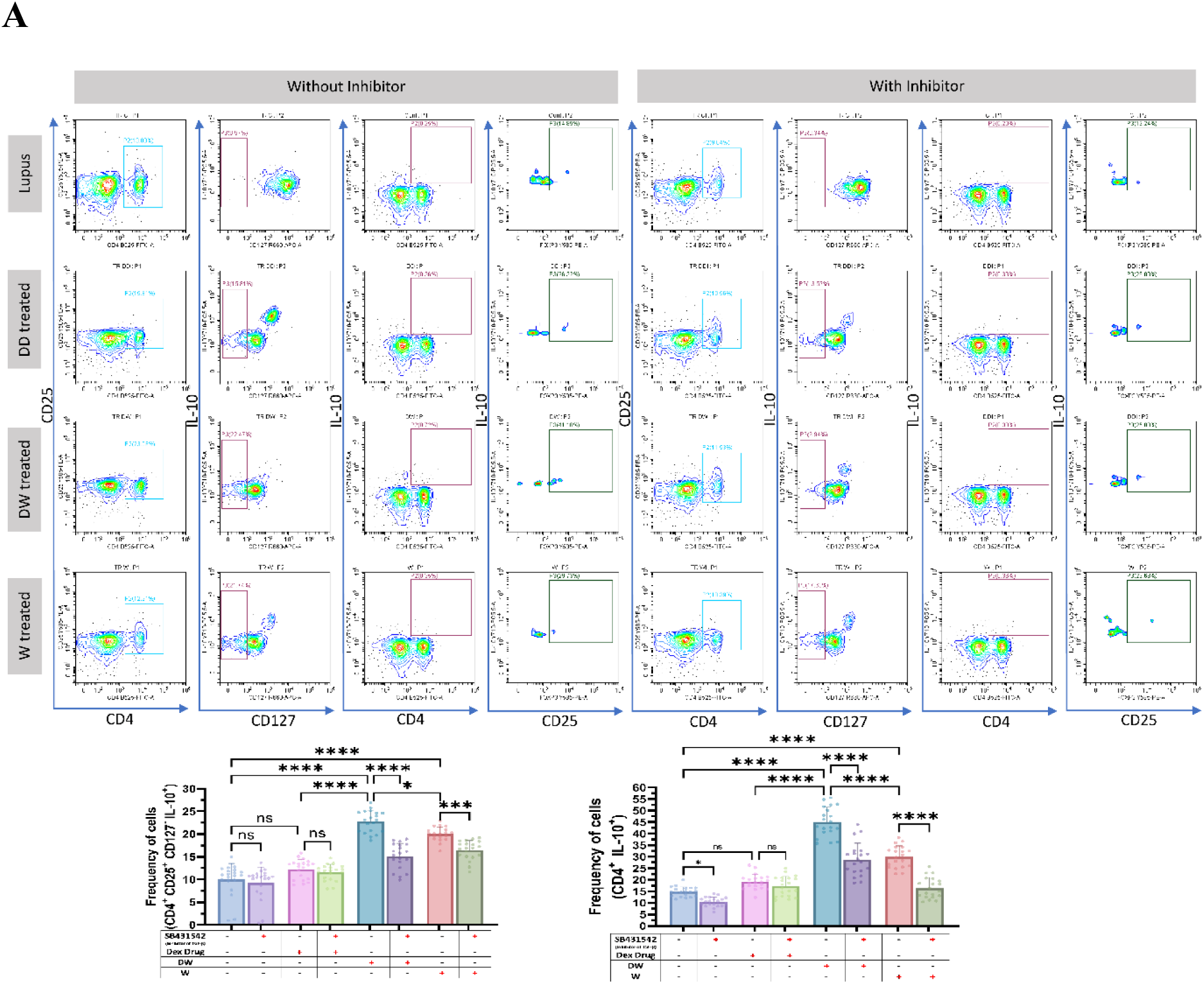
DW treatment mediates its immunomodulatory activity through TGF-β. SB431542 was used to inhibit TGF-β activity. It was added to the DW treated culture for 24 h and treated with Dex drug, DW and W for 24 hours and checked for the frequency of (A) Tregs. The contour plots depict findings from an individual sample of 20 patients with SLE, and corresponding bar graphs are presented with a sample size of 20. Error bars show mean ± SD. p values indicate significant changes as follows: non-significant (ns) p > 0.05, *P < 0.05, **p < 0.01, ***P < 0.001 and ****P < 0.0001; One-way ANOVA.

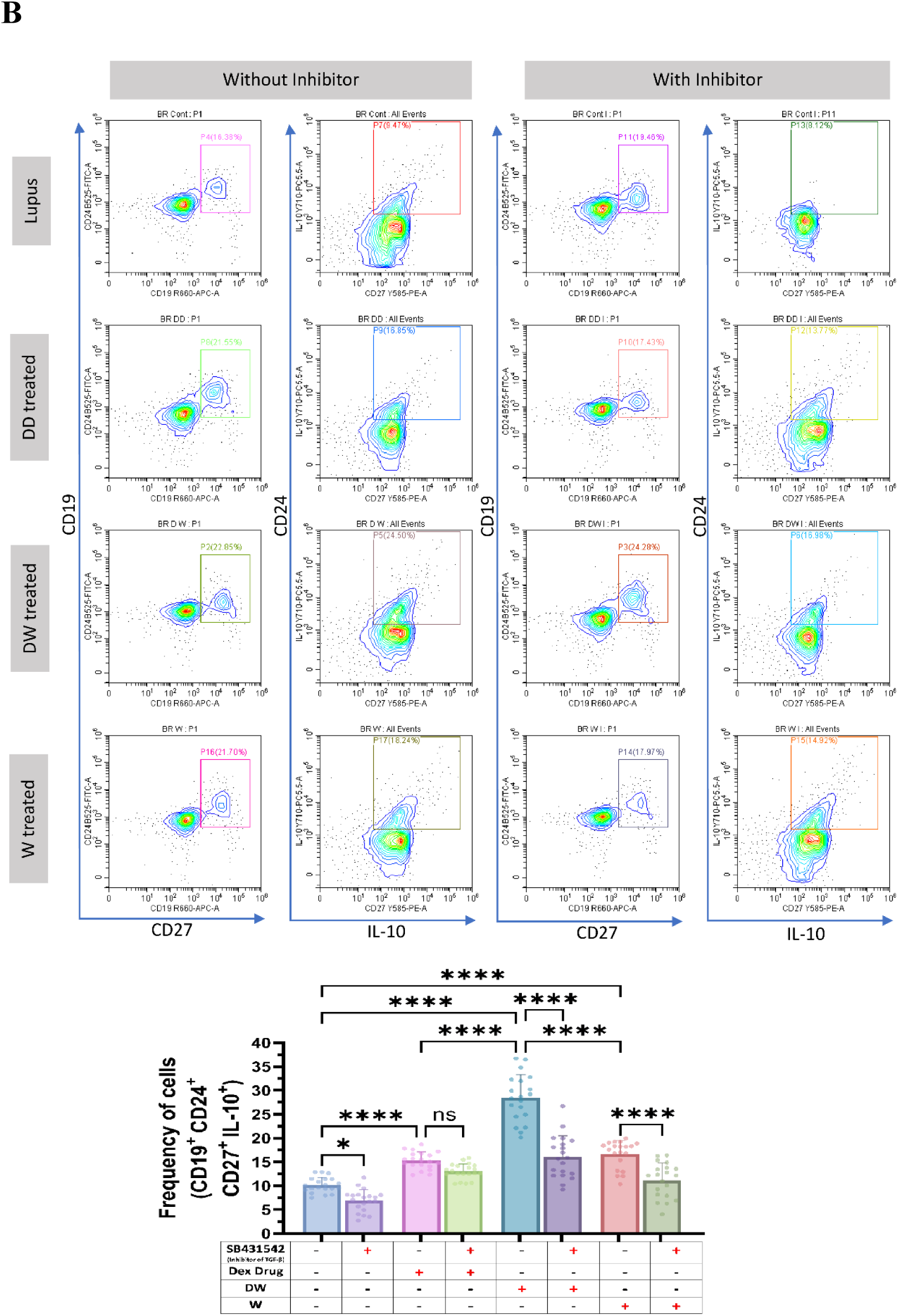
(B) SB431542 was used to inhibit TGF-β activity. It was added to the DW treated culture for 24 h and treated with Dex drug, DW and W for 24 hours and checked for the frequency of Bregs. The contour plots depict findings from an individual sample of 20 patients with SLE, and corresponding bar graphs are presented with a sample size of 20. Error bars show mean ± SD. p values indicate significant changes as follows: non-significant (ns) p > 0.05, *P < 0.05, **p < 0.01, ***P < 0.001 and ****P < 0.0001; One-way ANOVA.

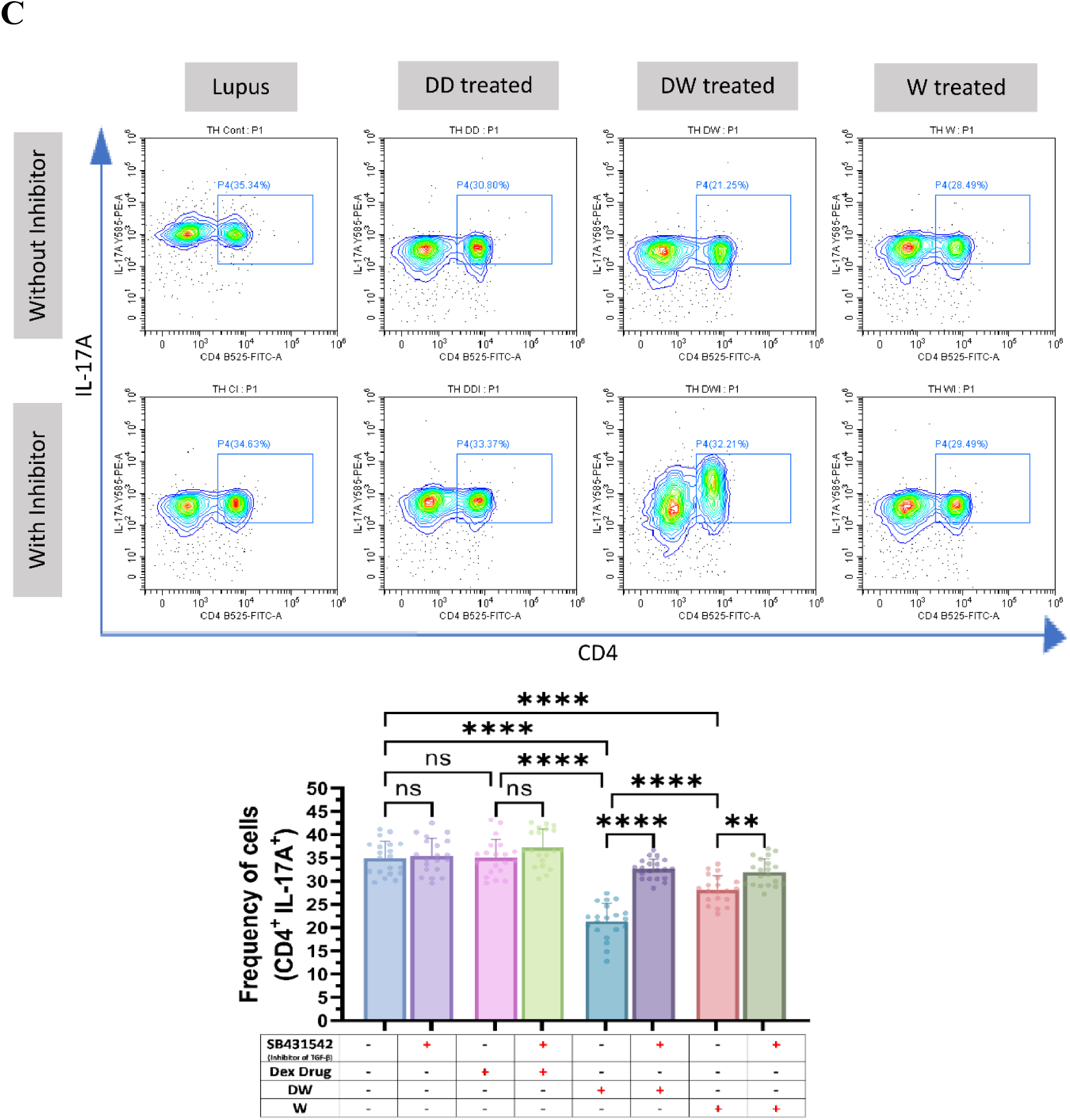
(C) SB431542 was used to inhibit TGF-β activity. It was added to the DW treated culture for 24 h and treated with Dex drug, DW and W for 24 hours and checked for the frequency of Th17. The contour plots depict findings from an individual sample of 20 patients with SLE, and corresponding bar graphs are presented with a sample size of 20. Error bars show mean ± SD. p values indicate significant changes as follows: non-significant (ns) p > 0.05, *P < 0.05, **p < 0.01, ***P < 0.001 and ****P < 0.0001; One-way ANOVA.

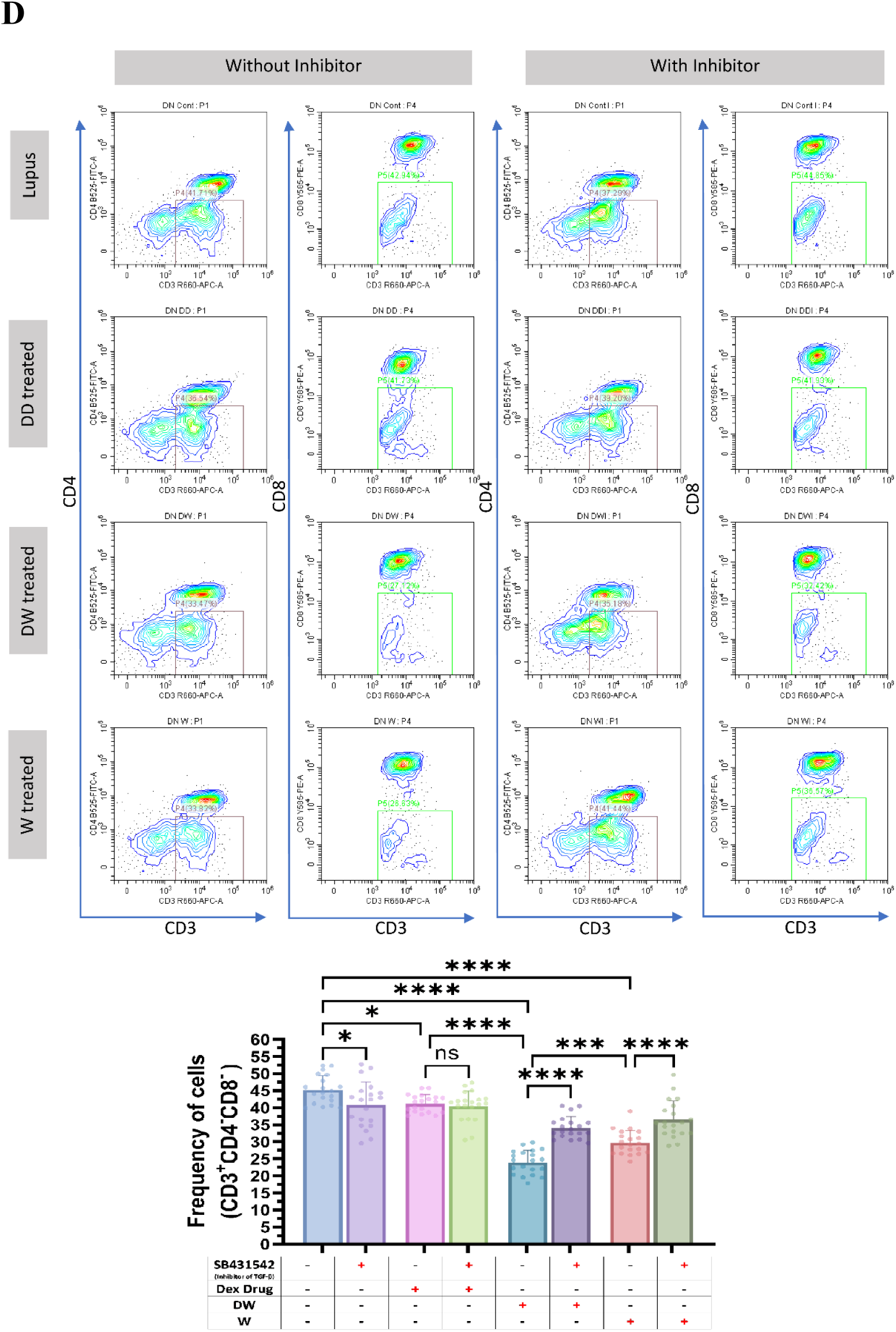
(D) SB431542 was used to inhibit TGF-β activity. It was added to the DW treated culture for 24 h and treated with Dex drug, DW and W for 24 hours and checked for the frequency of DN T cells. The contour plots depict findings from an individual sample of 20 patients with SLE, and corresponding bar graphs are presented with a sample size of 20. Error bars show mean ± SD. p values indicate significant changes as follows: non-significant (ns) p > 0.05, *P < 0.05, **p < 0.01, ***P < 0.001 and ****P < 0.0001; One-way ANOVA.

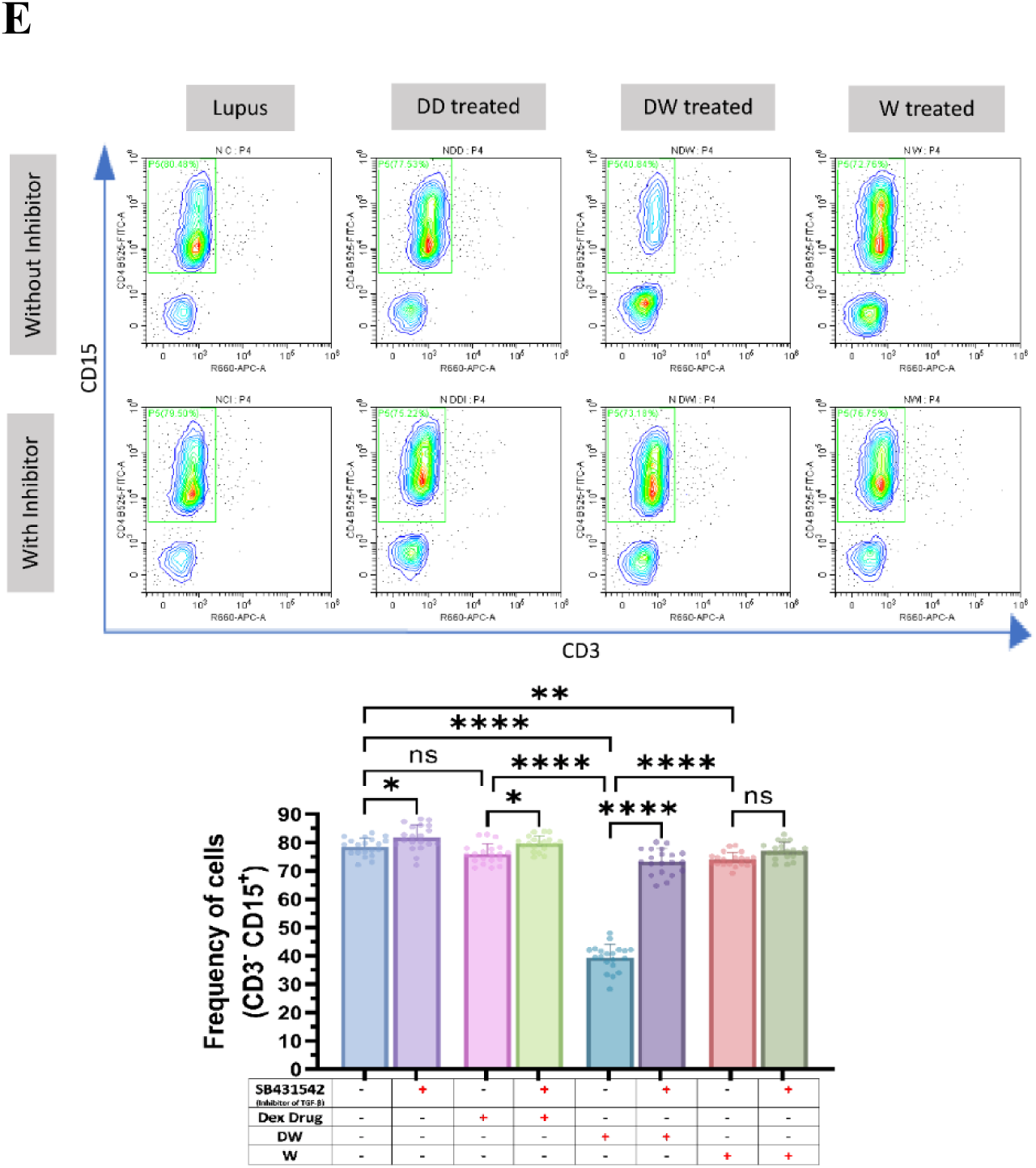
(E) SB431542 was used to inhibit TGF-β activity. It was added to the DW treated culture for 24 h and treated with Dex drug, DW and W for 24 hours and checked for the frequency of inflammatory neutrophils. The contour plots depict findings from an individual sample of 20 patients with SLE, and corresponding bar graphs are presented with a sample size of 20. Error bars show mean ± SD. p values indicate significant changes as follows: non-significant (ns) p > 0.05, *P < 0.05, **p < 0.01, ***P < 0.001 and ****P < 0.0001; One-way ANOVA.

### DW treatment suppressed autoantibodies, IL-17A and increases IL-10 and TGF-β expression

We evaluated the effect of DW treatment on autoantibodies generation from SLE patients PBMCs. Levels of anti-dsDNA and anti-ENA autoantibodies were detected in the cell supernatant using ELISA in different treatment groups including DW, W, Dex drug and untreated cells *in vitro*. Findings revealed that DW and W treatment were associated with a significant decrease in the supernatant levels of anti-dsDNA (Figure 4 A) and anti-ENA (Figure 4 B) as compared to that in non-treated cells (lupus). Levels of cytokines IL-10, and TGF-β were detected in PBMCs supernatant using ELISA, and mRNA expression of cytokine IL-10 and Th17 cell-mediated factor IL-17A were detected by Real Time (RT) PCR. IL-10 cytokines (Figure 4 C) and TGF-β (Figure 4 D) were found to be significantly increased in the DW and W treated groups compared to the non-treated group. In contrast, there was non-significant difference in the levels of TGF-β, and cellular expression of IL-10 (Figure 4 E) in Dex drug treated cells but there was a significant decrease in the IL-17A expression (Figure 4 F) when compared to non-treated group.

**Figure 4:**
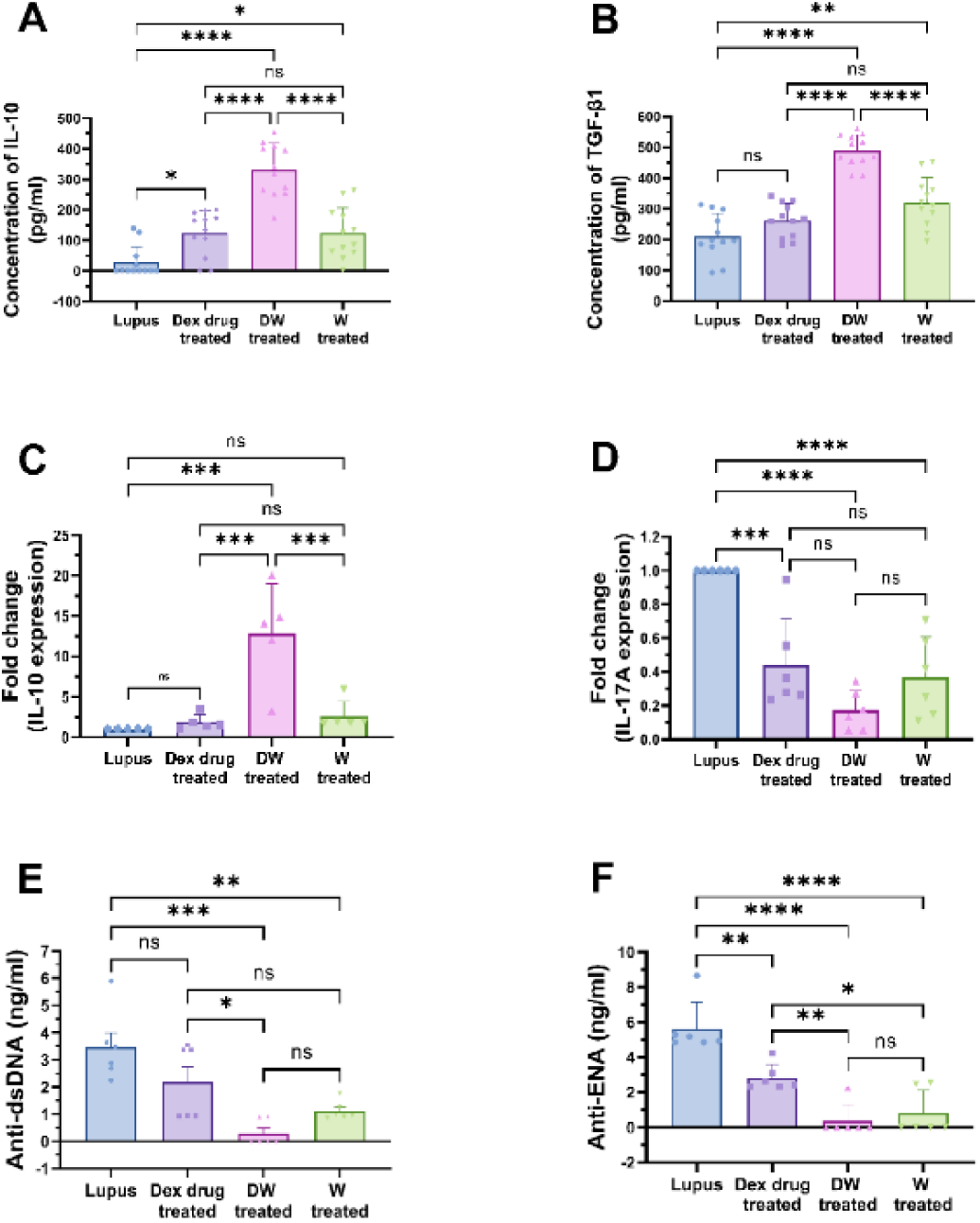
Immunomodulatory effect of DW treatment on IL-10, TGF-β, IL-17A expression and on the autoantibodies secretion. Bar graph showing the concentration of (A) IL-10 cytokines (n=12), (B) TGF-β (n=12) and the relative expression of (C) IL-10 (n=5), (D) IL-17A gene expression (n=6) in Dex drug, DW and W treated PBMC’s with that of non-treated cells (lupus). DW treatment significantly suppressed the (E) anti-dsDNA (F) anti-ENA autoantibody levels in the supernatant as compared to that in lupus (non-treated cells). The bar graphs data represent mean of 6 patients sample. All values are represented as mean ± SD. non-significant (ns) p > 0.05, ***P < 0.001 and ****P < 0.0001; One-way ANOVA.

### Preclinical studies

Over a period of two months, 75 female BALB/c mice were raised, with 65 mice received intraperitoneal pristane over the next month to induce lupus-like autoimmunity (Yun et al., 2023). Ten unimmunized mice served as controls. Following the development of autoantibodies, the pristane-induced mice were divided into four groups (15 mice per group): i) lupus (treated with normal saline), ii) Dex drug (received Dex drug treatment), iii) DW (received Dex-primed CM treatment), and W (received CM treatment). The groups were further categorized based on distinct autoantibody profiles, such as the presence of anti-dsDNA^+^, anti-ENA^+^, and anti-dsDNA^+^ + anti-ENA^+^ autoantibodies (n=5 in each category). After 35 days of pristane induction, serum samples were analyzed for autoantibody levels. Body weight was monitored, and urine samples were examined weekly for proteinuria (Figure 5 A). Blood samples were collected from the tail vein of each mouse (n=60) before and after 30 days of pristane injection. Autoantibodies levels were measured in the serum and a significant increase in the levels of anti-dsDNA and anti-ENA autoantibodies was observed over a span of one month (post-induction) (Figure 5 B and C). Twenty-five mice were reported positive for anti-dsDNA, 25 positive for anti-ENA and 10 had both the autoantibodies present.

**Figure 5:**
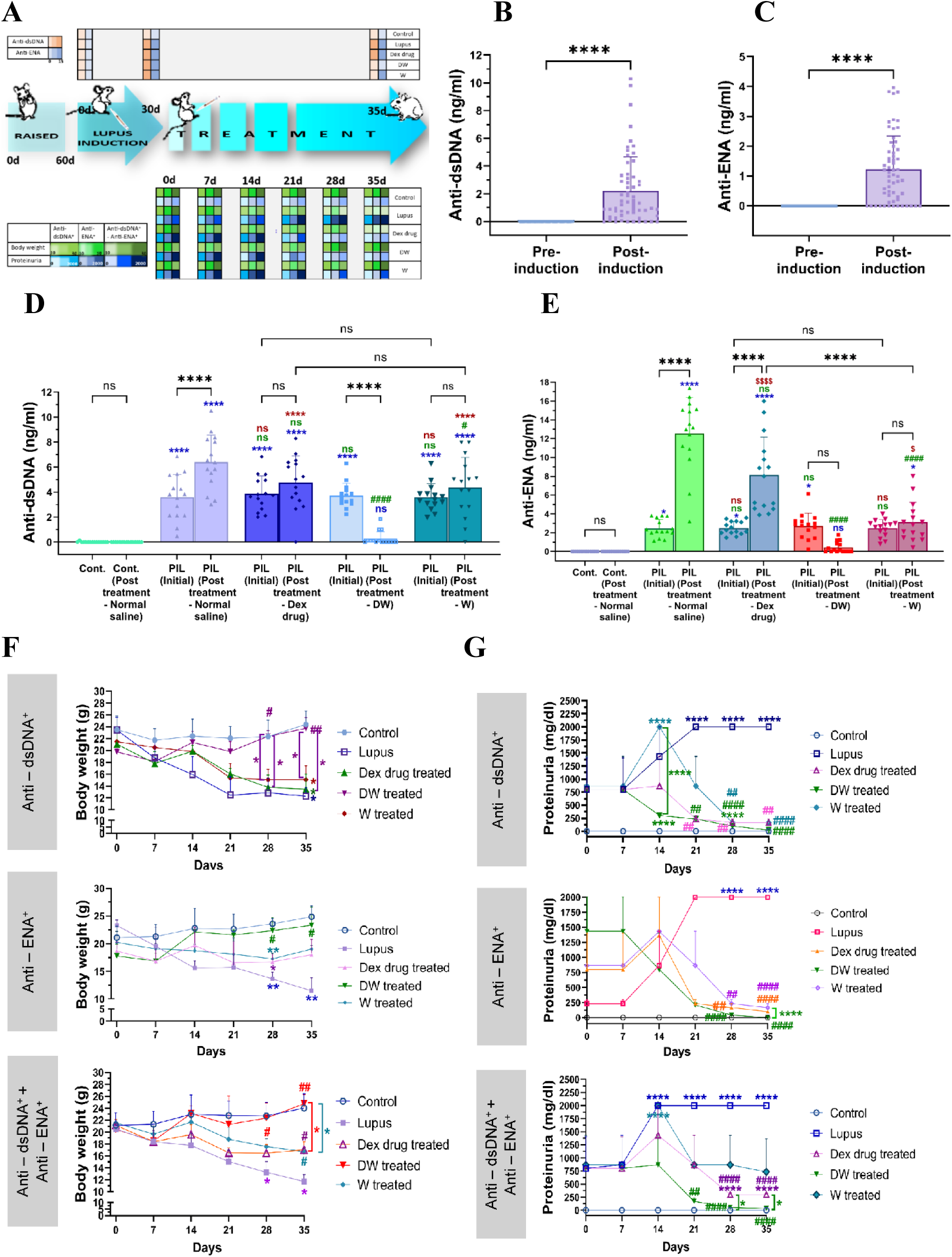
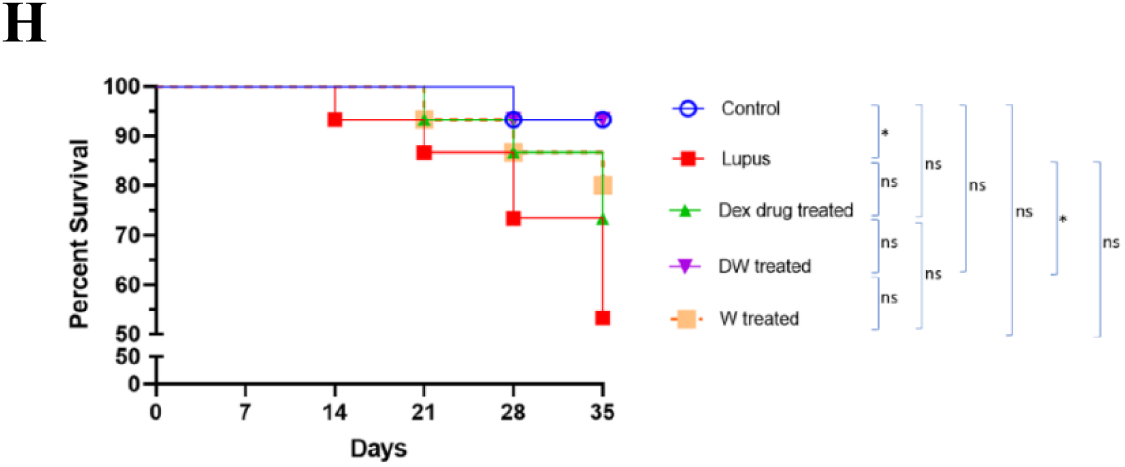
Generation of autoantibodies in PIL mice and the immunomodulatory effects of different treatments on the autoantibodies post-induction, proteinuria reduction, maintenance of optimal body weight and mortality in PIL mice. (A) Experimental outline. There was significant difference in the level of (B) anti-dsDNA and (C) anti-ENA autoantibodies over a span of one month (post-induction). The DW-treated group demonstrated a significant decrease in (D) anti-dsDNA and (E) anti-ENA compared to the pre-treatment levels and other groups post-induction. (F) Body weight and (G) proteinuria was measured weekly till 35^th^ day of various treatment and DW significantly suppressed proteinuria and helped in maintenance of optimum body weight in all the subgroups (anti-dsDNA^+^, anti-ENA^+^, and anti-dsDNA^+^ + anti-ENA^+^). (H) Survival rate (percentage) post 7^th^, 14^th^, 21^st^, 28^th^ and 35^th^ day of treatment. Kaplan–Meier survival curves, n = 15, analyzed by log-rank test, * p < 0.05. The numbers of PIL mice surviving to 35 weeks of age post-treatment. The line graphs illustrate mean data from a sample size of 5 mice. Error bars show mean ± SD. p values indicate significant changes as follows: non-significant (ns) p > 0.05, *P < 0.05, **p < 0.01, ***P < 0.001 and ****P < 0.0001; One-way ANOVA. In Figure C and D, **\*** depicts significance compared to control, **#** depicts significance compared to PIL (normal saline treated) counterparts, **$** depicts significance compared to DW.

### DW treatment reduced autoantibodies, maintained optimum body weight, reduced proteinuria and mortality in PIL mice

#### Autoantibodies

The DW-treated group demonstrated a significant decrease in autoantibodies (both anti-dsDNA and anti-ENA) compared to the pre-treatment levels and other treatment groups. The Dex drug and W treated groups of mice did not show significant reduction in autoantibody levels as compared to their pre-treatment levels. Hence, DW treatment was able to significantly lower the level of autoantibodies in a span of 35 days. In contrast, PIL mice receiving only normal saline exhibited a continued rise in autoantibody levels (anti-dsDNA and anti-ENA) compared to the pre-treatment levels or other treatment groups (Figure 5 D and E).

#### Body weight

Post treatment PIL mice were monitored for body weight to check if these treatments affected these parameters differentially in mice with distinct autoantibodies’ responses. There was a significant loss in body weight starting from 28^th^ day post treatment in normal saline treated PIL mice in all three distinct autoantibody subgroups (anti-dsDNA^+^, anti-ENA^+^ and anti-dsDNA^+^ + anti-ENA^+^) as compared to normal saline treated non PIL control mice. The initial body weights were 23.45, 23.29 and 20.52 g which reduced to 12.28, 11.46 and 11.71 g in anti-dsDNA^+^, anti-ENA^+^ and anti-dsDNA^+^ + anti-ENA^+^ respectively. The body weights of DW treated mice in different autoantibody specific subgroups (anti-dsDNA^+^: 22.40 g, anti-ENA^+^: 22.35 g and anti-dsDNA^+^ + anti-ENA^+^: 22.34 g) showed significant improvement post 28 days of treatment as compared to normal saline treated PIL mice (anti-dsDNA^+^: 12.85 g, anti-ENA^+^: 13.63 g and anti-dsDNA^+^ + anti-ENA^+^: 13.24 g). Whereas Dex drug and W treated mice showed delay in improvement of body weight starting only after 35 days post treatment for the anti-dsDNA^+^ + anti-ENA^+^ subgroup, and the increase in body weights in these two treatment groups was significantly lower than that of the DW-treatment group (Figure 5 F).

#### Proteinuria and mortality

Urinary protein was quantified from urine samples of individual mice in metabolic cages on 0, 7^th^, 14^th^, 21^st^, 28^th^ and 35^th^ day of treatment. Urinary protein excretion in normal saline treated lupus mice on 21^st^ to 35^th^, 28^th^ to 35^th^ and 14^th^ to 35^th^ days in anti-dsDNA^+^, anti-ENA^+^, and anti-dsDNA^+^ + anti-ENA^+^ autoantibodies groups respectively was significantly higher than that of the normal saline treated non PIL control mice. Dex drug, DW and W treated groups showed significant decrease in proteinuria on 21^st^, 14^th^ and 28^th^ day of treatment respectively in the mice positive for anti-dsDNA^+^ anti-ENA^+^ and anti-dsDNA^+^ + anti-ENA^+^ autoantibodies, suggesting that DW treatment kept a better check on urine protein levels than Dex drug or W treatment (Figure 5 G).

In the anti-ENA^+^ group, Dex drug, DW, and W treatment had significantly lowered proteinuria on 28^th^ day of treatment compared to PIL mice receiving normal saline. Dex drug and DW treated groups of mice kept the urine protein levels in check from 21^st^ day of treatment until the last day of treatment in anti-dsDNA^+^ + anti-ENA^+^ mice. However, this decrease in urine protein in the DW treated mice positive for anti-dsDNA, anti-ENA, and anti-dsDNA + anti-ENA autoantibodies were significantly higher than that of Dex drug treated mice. Therefore, DW treated proteinuria in this PIL model more effectively than the other two treatments W and Dex drug in all three autoantibody subgroups setups (Figure 5 G). PIL mice treated with DW exhibited significantly prolonged survival compared to normal saline treated PIL mice, Dex drug treated, or W treated littermates. On the 35^th^ day of treatment, the survival rate for normal saline treated PIL mice was 53.33%, whereas it improved significantly to 93.33% in DW treated mice. There was no significant difference in the survival percentages between the Dex drug (73.33%) and W (80%) treatment groups compared to both lupus and non-lupus control mice (Figure 5 H).

### Inhibition of SLE- associated splenic Th17 cells, upregulation of Tregs, Bregs and anti-inflammatory cytokine IL-10 through TGF-β with DW treatment

We studied the impact of DW treatment on Treg, Breg and Th17 population in PIL mice (n=3). PIL mice showed significantly decreased splenic regulatory cells (Treg and Breg) and increased Th17 (CD4^+^ IL-17A^+^) cells compared to non PIL control mice receiving normal saline. Dex drug and W treatment did not affect Tregs (CD4^+^ CD25^+^ FOXP3^+^) and Bregs (CD5^+^ CD1d^+^) in lupus mice. However, DW treatment increased regulatory cells and decreased Th17 cells (Supplemental Figure 3 A). The Th17/Treg ratio was shifted towards Treg differentiation, with better reduction than Dex drug or W treatment (Supplemental Figure 3 B). IL-10 cytokine production by PIL mice upon different treatments was measured. It was observed that the decreased IL-10 production in PIL mice was significantly reversed with DW treatment, and this improvement was higher than that achieved with Dex drug or W treatment (Supplemental Figure 3 C). Next, we determined the TGF-β concentrations in the serum samples of mice, as there is growing evidence that TGF-β can regulate T-cell immunity in inflammatory diseases and promote the emergence of Tregs contributing to the induction and maintenance of natural and induced immune tolerance (Huai et al., 2021). The concentration of TGF-β in the serum of lupus was significantly lower than that of the control mice which was improved with DW treatment. Dex drug or W treated mice, did not show any significant change (Supplemental Figure 3 D).

### Alopecia was reversed in DW treated mice

Of the 60 PIL mice, 5 mice developed alopecia (of which one died), hence 4 mice were isolated and treated for the next 42 days with either DW, Dex drug, W or normal saline. On the 42^nd^ day, observations were made to assess the effect of treatments on the clinical manifestation of alopecia. PAI was performed using power Doppler to assess the effect of treatments on the vasculature changes under the affected alopecia patch (Figure 6 A). PIL mice developed progressive dorsal alopecia that started dorsally and gradually spread across the body, resulting in localized baldness. The affected skin regions were very dry and had lost pliability. After 42 days of treatment there was a visible reversal of hair loss observed in DW treated mice (Figure 6 B), and the regrowth of hair was progressive starting from days 7, 14, 21, 28, 35 of DW treatment (Figure 6 C). Reduced peripheral and subcutaneous blood flow contributes to hair loss. The blood supply to the scalp resembles that of the skin elsewhere in the body, originating from the subcutaneous tissue. Hence, we checked for the subcutaneous blood flow by using a power Doppler tool overlaid with B-mode imaging in the area losing hairs and observed loss of vasculature in the PIL mice. Upon DW treatment restoration of vasculature and increased blood flow was observed but none to minimal restoration seen in Dex drug or W treated mice (Figure 6 D).

**Figure 6:**
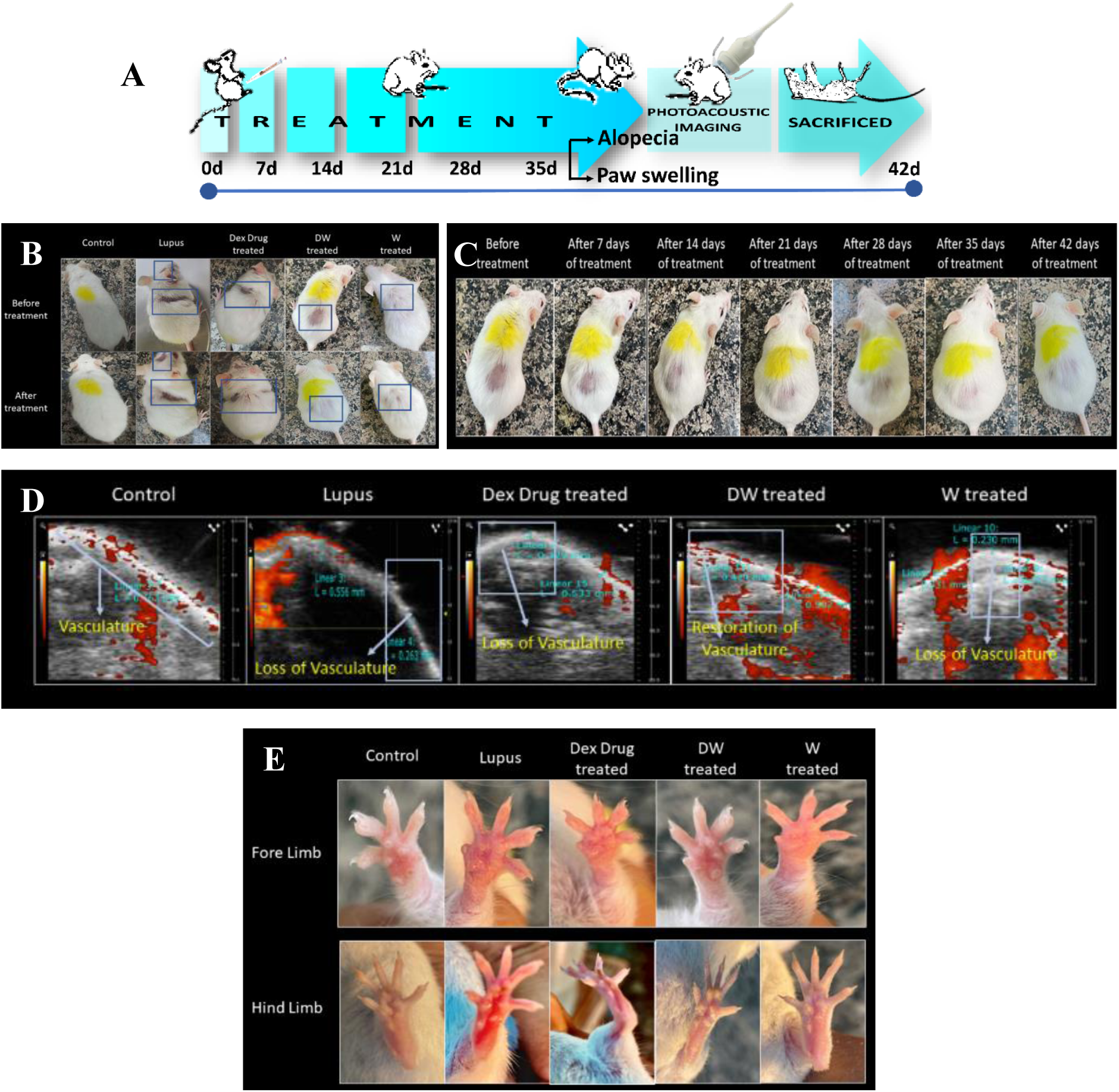
DW helps in the reversal of alopecia and reduced limb inflammation. (A) Experimental outline. Representative images of (B, C) alopecia, (D) power doppler image overlaid on B-mode image of area undergone hair loss and (E) limb inflammation in PIL mice before and after treatment with normal saline (lupus) and Dex drug, DW, W and normal saline (healthy, control) for duration of one month. Rectangles indicate the area impacted with hair loss.

### DW treatment reduced limb inflammation in lupus mice

Clinical evidence of arthritis, paw swelling, and loss of grip strength after pristane injection in BALB/c mice has been reported earlier (Leiss et al., 2013). Post 42 days of pristane induction, in concordance with the earlier reports we observed a marked increase in redness and swelling of the hind and fore limbs’ paws in PIL mice which reduced markedly with DW and W treatment but not with Dex drug-treatment (Figure 6 E).

### Seizure developed in PIL mouse was resolved with DW treatment

In 1999, the ACR established a standard nomenclature with case definitions for 19 neuropsychiatric conditions, 12 related to central nervous system manifestations mainly seizures, headache, stroke, depression, cognitive dysfunction, and psychosis (Zardi et al., 2014). In our study, among the 60 PIL mice, two mice developed seizures. Seizure refers to a sudden, uncontrolled electrical disturbance in the brain that results in abnormal behavior, sensations, or movements. Seizures in lupus may occur as a neurological manifestation of the disease, known as neuropsychiatric lupus. One mouse was administered normal saline, and the other was treated with DW. Mice that received normal saline died on the third day of treatment, whereas the DW treated mice recovered progressively from seizures in 7 days (Video 1-3).

### Reversal of organ damage and histopathological changes in DW treated mice

Post 35 days of treatment schedule, the mice were sacrificed under carbon dioxide, and their organs like kidneys, liver, lungs, heart, and spleen were harvested, morphological changes were observed, their size and weight was measured, and histological examination were performed (Figure 7 A).

**Figure 7:**
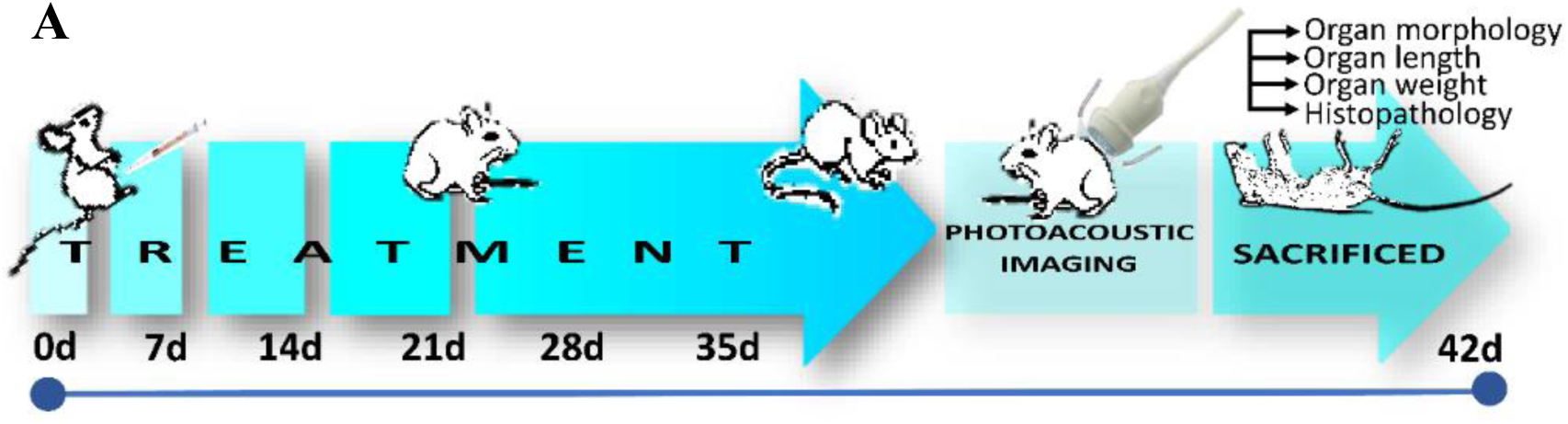
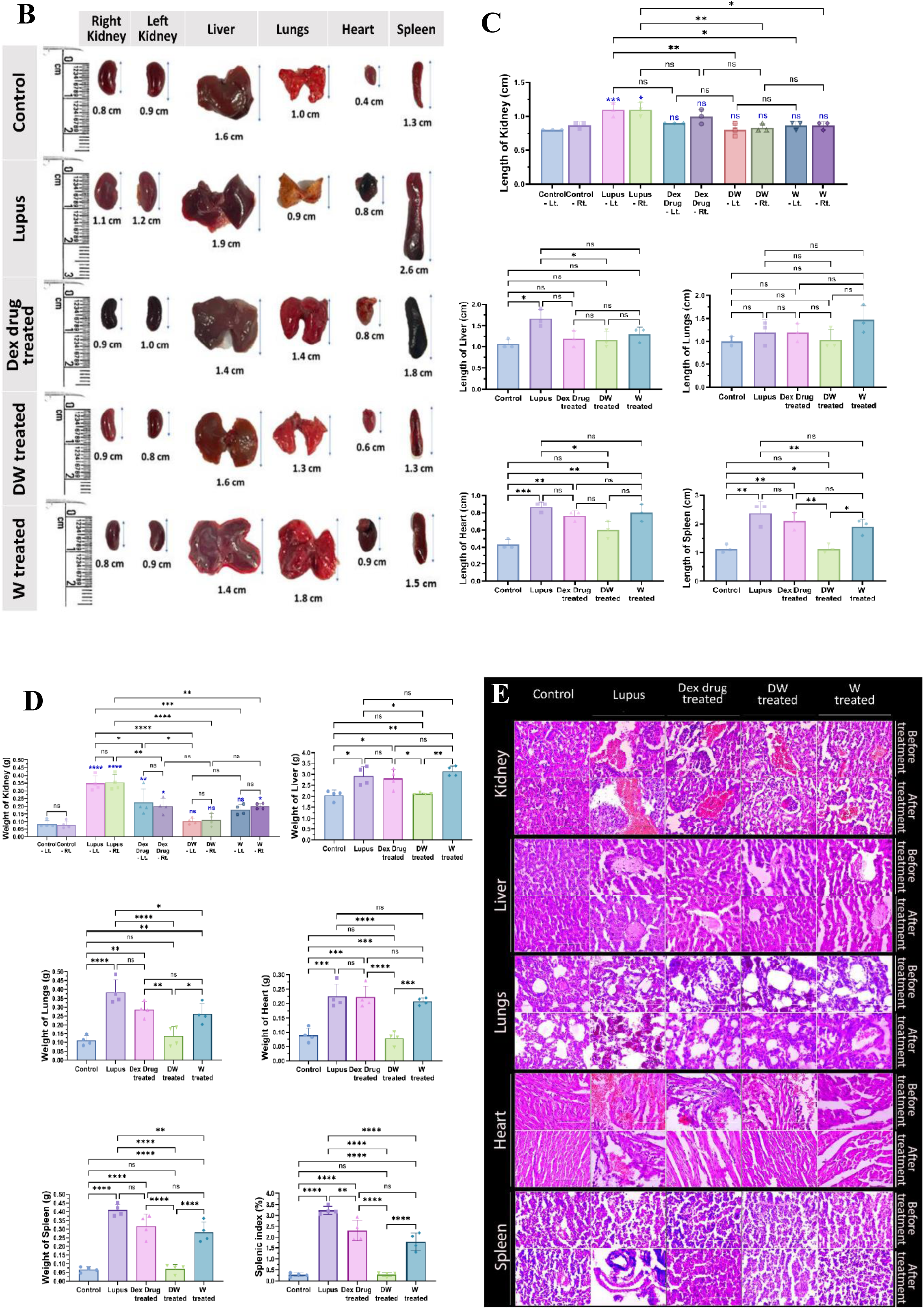
Reversal of organ damage and histopathological changes with DW treatment. (A) Experimental outline. Following a 35-day treatment regimen, mice were euthanized using carbon dioxide and organs like kidneys, liver, lungs, heart, and spleen were collected. (B) Morphological changes were observed, and (C) measurements of size (n=3) and (D) weight (n=4) were recorded. Additionally, (E) histological examinations (n=3) were conducted. Error bars show mean ± SD. p values indicate significant changes as follows: non-significant (ns) p > 0.05, *P < 0.05, **p < 0.01, ***P < 0.001 and ****P < 0.0001; One-way ANOVA.

#### Weight and length of organs

Gross observations of the systemic organs in all groups are shown in Figure 7 B. The PIL mice exhibited disease specific changes in morphology and swelling compared to the uninduced healthy control mice. Treatment with DW significantly normalized lengths and weights of kidney, liver, lungs, heart and spleen compared to other treatments (Figure 7 C and D).

#### Histopathological analyses

##### Kidney

Further, when we examined the histological images of hematoxylin and eosin (H & E) stained sections of the kidneys, the morphology and structure of the glomeruli and renal tubules in the mice of the control group were normal with no obvious abnormalities. In the PIL group, tubular epithelial cell shedding and basement membrane nudity, vacuolar degeneration of tubular epithelial cells, protein casting, tubular dilation, loss of tubular brush border, hyperplasia of mesangial cells, epithelial and endothelial cells, and edema-like degeneration of renal tubules were observed. However, after treatment with DW, the above pathological changes in the kidneys of PIL mice were alleviated majorly, though protein casts were present at a few sites. In contrast only a minor pathological improvement was seen in both Dex drug or W treated groups, as there still remained obvious intraparenchymal hemorrhage, red blood cell extravasation, dilated vascular spaces, fibrinous deposits, and protein cast in the renal section in the kidney section of lupus mice (Fig.7 E).

##### Liver

Liver H&E staining showed expansion of the portal tract and activity of the portal inflammatory infiltrate with extension to the lobule, causing direct damage to the hepatocytes. Morphological alterations were identified mainly in the hepatic lobule, with a storage spectrum of fine droplets. In several visual fields, histiocyte clusters were observed, conditioning the expansion of the sinusoid in the form of epithelioid granulomas, without necrosis or microgranulomas. Not only was intracytoplasmic accumulation of lipids noticeable, but its association with inflammatory changes was also observed, for instance, degeneration of hepatocytes, satelitosis (hepatocytes circumscribed by inflammatory cells, here with neutrophils and histiocytes), and acidophilic bodies (degenerated hepatocytes with condensed cytoplasm and inconspicuous nuclei). Such histopathological changes in liver have been reported (González-Regueiro et al., 2020). DW treatment significantly reduced the occurrence and severity of liver damage, though expanded portal tracts and acidophilic bodies were still present. In contrast, there was no improvement in the Dex drug and W treated groups (Fig.7 E).

##### Lungs

Histopathological findings suggested that the lungs of the control mice had normal alveolar walls and capillaries. Pneumocytes, alveolar lumina, and blood vessels were also normal. Few alveolar macrophages, which are normally present in the alveolar wall, have been observed earlier too (Rathore et al., 2017). Bronchial walls and epithelial cells were normal. On the other hand, PIL mice and normal saline treated PIL mice had collapsed alveolar lumina and accumulation of numerous inflammatory cells in and around the alveolar lumina and alveolar wall. The PIL mice treated with DW showed normal alveolar walls, alveolar lumina, minimal alveolar expansion, and pulmonary blood vessels which could not be observed with Dex drug or W treatment (Fig.7 E).

##### Heart

In SLE, cardiac manifestations such as coronary artery disease and myocarditis are the leading causes of morbidity and mortality. Heart problems commonly associated with lupus include pericarditis or pericardial effusion, valve abnormalities, myocarditis, arrhythmias, and accelerated atherosclerosis (Burkard et al., 2018). We studied gross histological changes in the heart of PIL mice and Dex drug, DW, W treated groups, and compared the results with normal saline treated PIL mice. The heart of the control group showed a normal architecture without edema or neutrophil infiltration. The histological study of lupus-induced myocardium showed edema, wavy myocardial fibers due to slippage of myofibrillar alignment, infarcted zones, necrosis of widespread area of muscle fiber, separation of cardiac muscle fibers, and neutrophil infiltration, which are characteristic of myocardial ischemia as compared to control. There was no improvement in cardiac histopathology in lupus mice after treatment with normal saline, but it was significantly improved in DW treated mice with reduction in hypertrophied cardiomyocytes, inflammatory cell infiltration, wavy myocardial fibers, and necrosis (Fig.7 E).

##### Spleen

Splenomegaly has been reported as a manifestation of active SLE (Yang et al., 2018). Splenomegaly can be caused by increased splenic function, congestion, or infiltration (Palmiere et al., 2019). Although the spleen is not considered a common target organ in SLE, its function in producing antibodies cannot be neglected. Thus, it is important to understand the effect of treatment on the pathogenesis and features of splenomegaly and inflammation of the spleen in lupus. Histological examination of the spleens of PIL mice revealed increased cellularity, severe degenerative changes as ill-defined lymphoid follicles with large necrotic foci by darkly stained cells accompanied with edema, and diffused degenerated lymphoid cells compared with that in control mice. Thirty-five days of DW treatment significantly improved splenic section whereas Dex and W treated mice still had an ill-defined spleen architecture with lost and loose cells accompanied by scattered cells (Fig.7 E).

### PAI of anatomical and physiological status of organs

#### Kidney

In PAI brightness Mode (B-Mode) refers to the ensuing two-dimensional grayscale picture of a region of interest (ROI), which was used to locate the kidneys, right and left in the abdomen of the mice (Figure 8 A; B-mode). A scale is located on the right side of each image in the first row, which shows the distance of structures, those farther away from the transducer at the bottom, and the tissue closest to it at the top, while also providing depth and size information. Both kidneys can be observed in white and light gray tones because as density increases, more echo is recorded, resulting in a brighter signal on the screen. B-Mode provides the best anatomical structure visualization; hence it was used to check the effect of treatments on anatomical differences of both the kidneys. The second row depicts the power doppler image that displays the traced boundary of the outer kidney wall within a red loop (Figure 8 A; PA signal laid on B-mode). This image showcases the vascularity of the kidney combined with 3D-Mode, provided volumetric information and spatial resolution of the kidney (Figure 8 A, rendered volume for 3D surface view with PA signal in wireframe and wireframe view of 3D volume). Consistent with Figure 7 B and C, Figure 8 A also showed a significant increase in the volume of kidney of the PIL mice. DW treatment normalized the kidney sizes and volumes significantly

**Figure 8:**
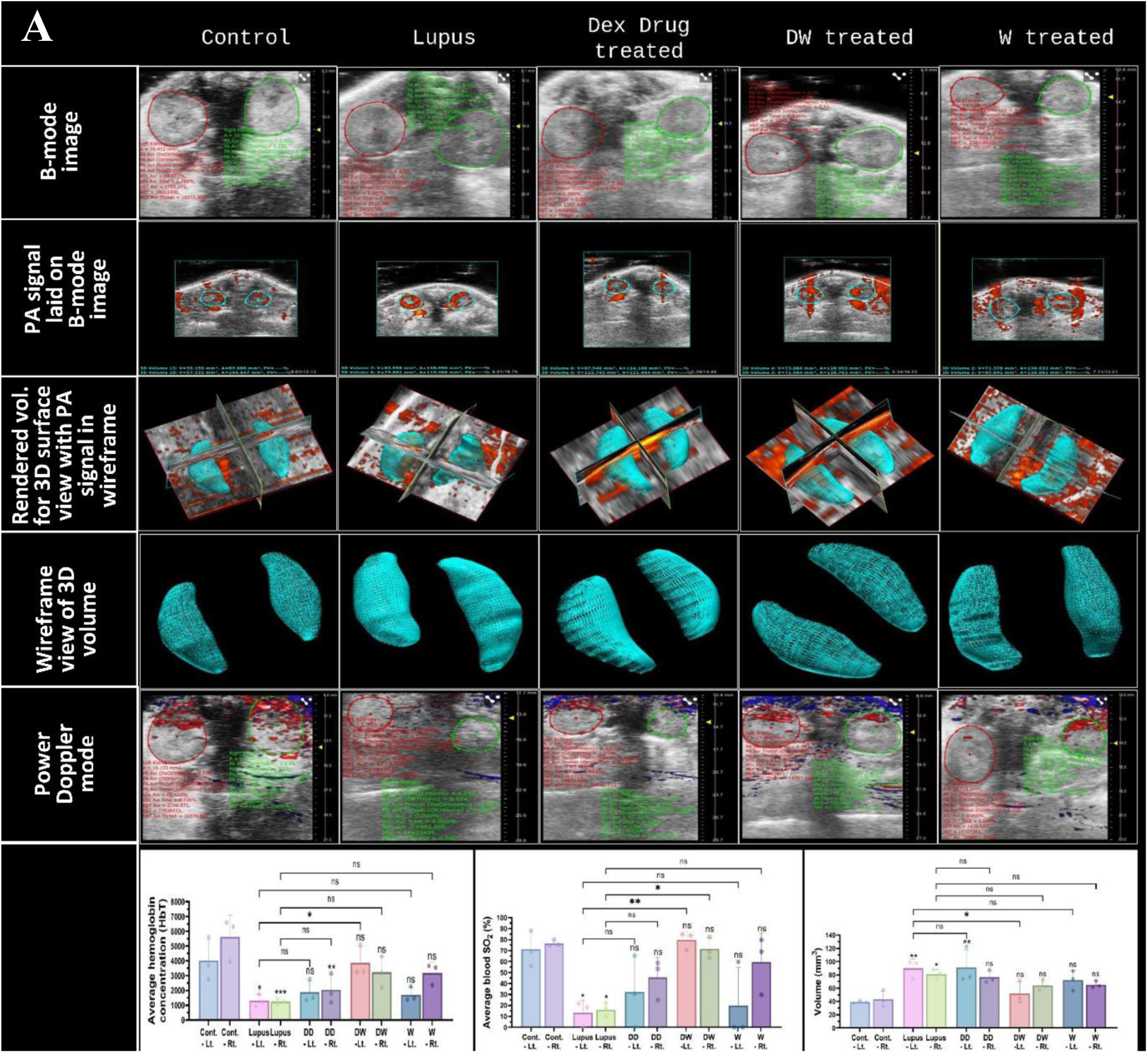
DW treatment improves anatomical and physiological status and function of vital organs. Representative US and PA overlay scanned images of kidneys from healthy (control), lupus, Dex drug, DW and W treated mice (n=3). (A) The confined red, green and blue lines denote the region of interest (ROI) registered during US, PA and power doppler and colour doppler imaging and quantitation for the PA images. Cross-view representation of the 3-dimensional volumetric reconstruction of PA imaging in the mice kidneys based on the ROI as shown in overlay images. The graph depicts the 3-D volume, average sO_2_ and HbT in the kidneys of different treatment groups. Error bars show mean ± SD. p values indicate significant changes as follows: non-significant (ns) p > 0.05, *P < 0.05, **p < 0.01, ***P < 0.001 and ****P < 0.0001; One-way ANOVA. Rt., Right; Lt., Left

Power Doppler mode, overlaid on top of B-mode image for anatomical reference, displays blood flow in the area of interest in red and blue gradients (Figure 8 A; Power Doppler mode). Blood flowing towards the transducer is represented by a gradient between red (lowest velocity) and white (highest velocity), and blood flowing away from the transducer is represented by a gradient between blue (lowest velocity) and white (highest velocity). The percent renal oxygen saturation (sO_2_) and average hemoglobin concentration (HbT) were significantly lower in the PIL group than in the control group. Analysis of average blood sO_2_ and HbT in different treatment groups revealed a significant increase in both these parameters only in the DW treated group (Figure 8 A; volume, average blood sO_2_ and HbT concentration graphs).

#### Liver

In Figure 8 B, the liver is visible in both ultrasound (US) (B-mode) and photoacoustic mode (PA signal laid on B-mode). The red loop in all the figures indicates the liver ROI, which was drawn manually based on the US image. The liver had homogeneous echotexture in DW treated mice and healthy control mice but hyperechoic in PIL and Dex drug-treated mice. Notably, the average hepatic oxygen saturation was found to be significantly decreased in the PIL, Dex drug, and W groups of mice compared to the control and DW groups. The average blood sO_2_ and HbT levels were not significantly changed in the DW group of mice when compared to the control (Figure 8 B; average blood sO_2_ and HbT concentration graphs).

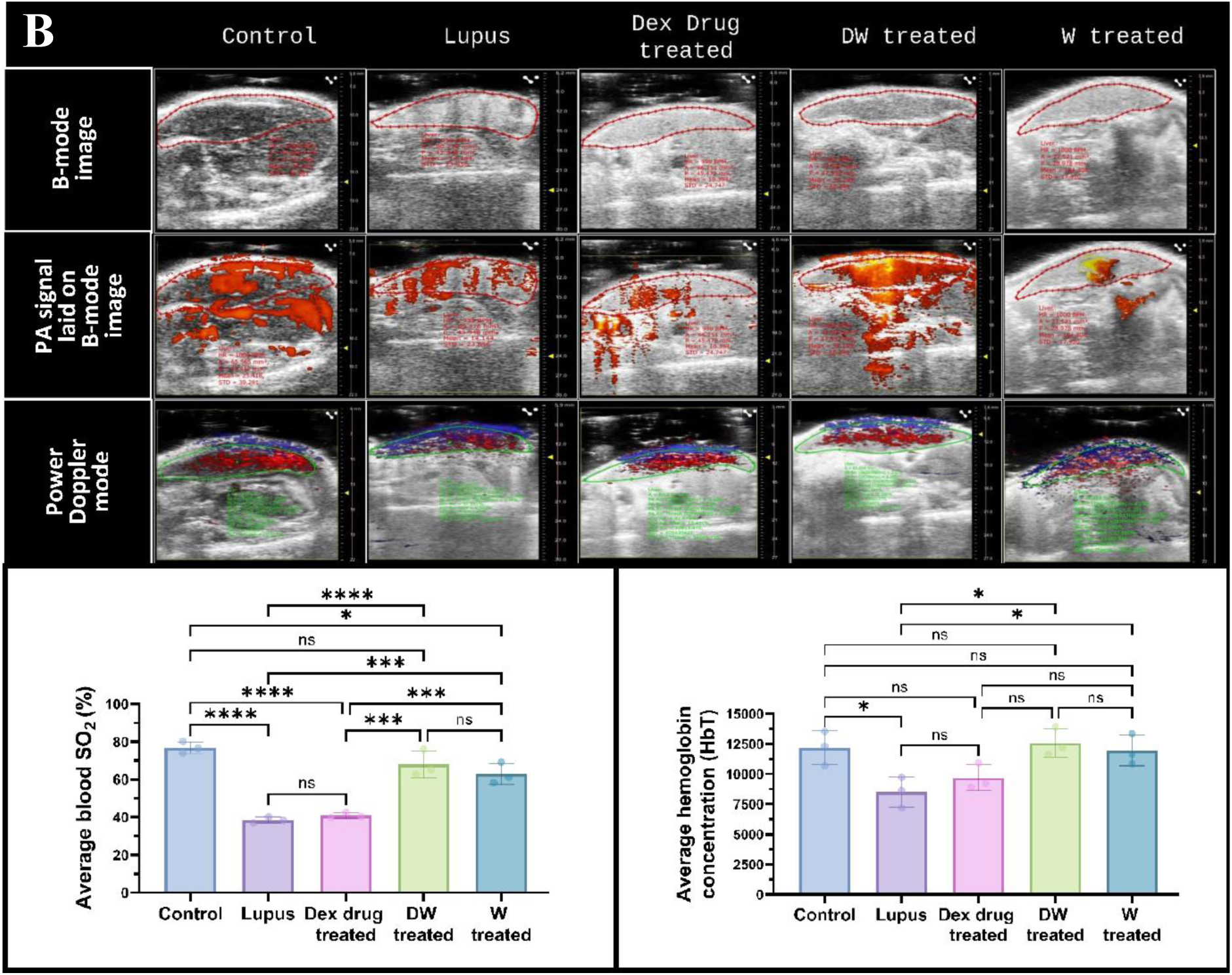
(B) Representative US and PA overlay scanned images of liver from healthy (control), lupus, Dex drug, DW and W treated mice (n=3). The confined red and green lines denote the ROI registered during PA, power doppler, colour doppler imaging and quantitation for the PA images. The graph depicts the average sO_2_ and HbT in the liver of different treatment groups. Error bars show mean ± SD. p values indicate significant changes as follows: non-significant (ns) p > 0.05, *P < 0.05, **p < 0.01, ***P < 0.001 and ****P < 0.0001; One-way ANOVA.

#### Heart

Various cardiac parameters were also evaluated, using US (Figure 8 C; B-mode) and PAI (Figure 8 C; Power doppler mode). The ROI that was manually generated based on the US image shows the heart in the mice treated with lupus, Dex, DW, W and control within the red loop. Interestingly, compared to control and DW-treated mice, the average blood sO_2_ and HbT levels were considerably lower in the lupus-, Dex-, and W groups of mice whereas in the DW group, the average blood sO_2_ and HbT levels were similar to those in the control group (Figure 8 C; average blood sO_2_ and HbT concentration graphs).

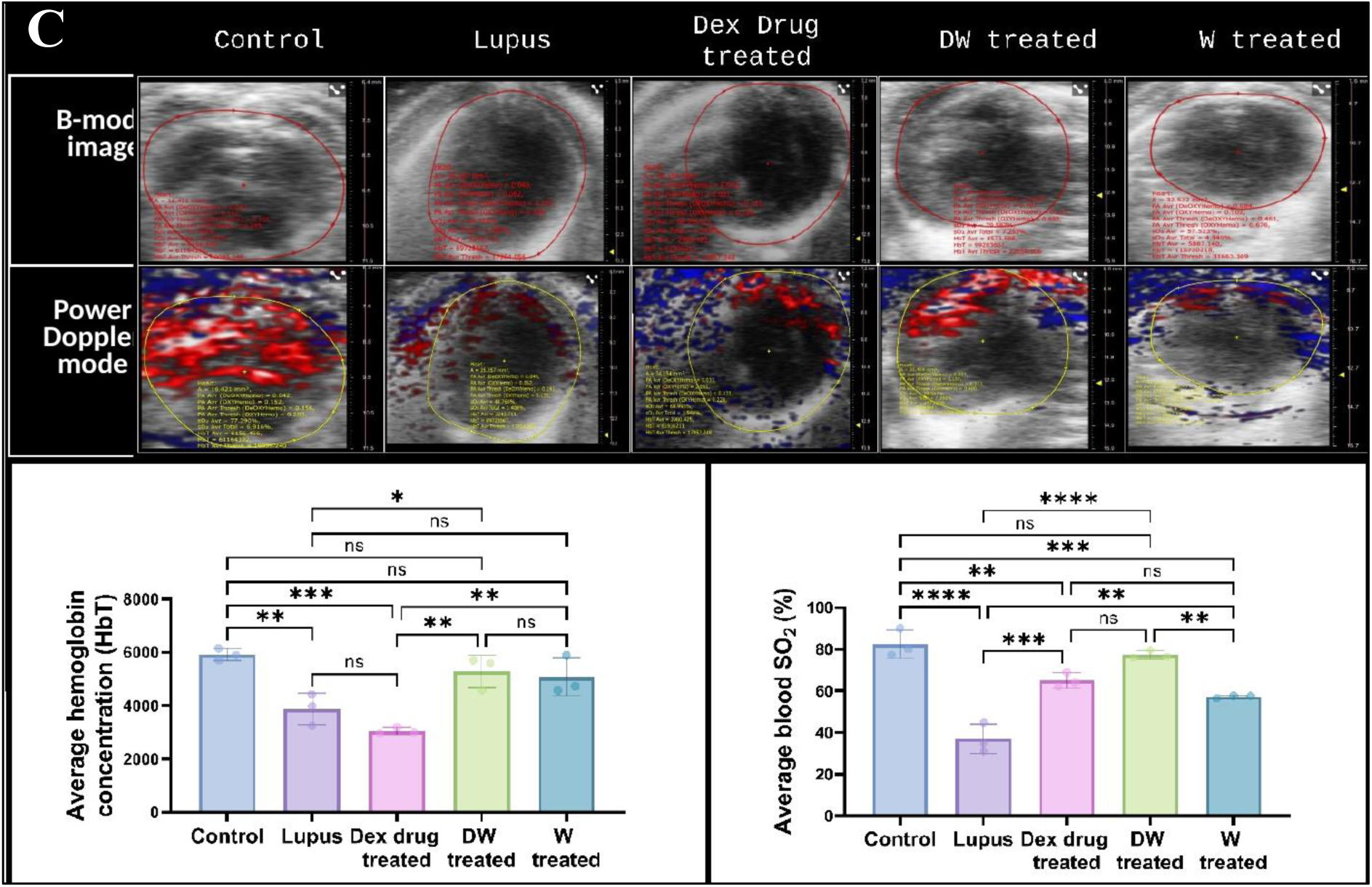
(C) Representative US and PA overlay scanned images of heart from healthy (control), lupus, Dex drug, DW and W treated mice (n=3). The confined red and yellow lines denote the ROI registered during PA imaging and quantitation for the PA images. B-mode image show parasternal long-axis view at the level of the papillary muscle. M-mode image of the LV displays dimensions of the ventricular walls, LV cavity, and cardiac function measurements. y-axis represents the distance (in mm) from the transducer; time (in ms) is on the x-axis. The graph depicts the average sO_2_ and HbT in the heart of different treatment groups. Error bars show mean ± SD. p values indicate significant changes as follows: non-significant (ns) p > 0.05, *P < 0.05, **p < 0.01, ***P < 0.001 and ****P < 0.0001; One-way ANOVA.

Echocardiography was performed to evaluate cardiac geometry and function. In lupus patients, the direct influence of heart rate and stroke volume on cardiac output has been well established. Reduced cardiac output can lead to inadequate blood supply, compromising essential physiological processes and potentially causing tissue and organ damage. Left unaddressed, diminished cardiac output is frequently associated with heart failure, which is a common diagnosis. In individuals with SLE, cardiac output undergoes modifications, accompanied by alterations in stroke volume, which represents the volume of blood expelled from the heart with each contraction (Alghareeb et al., 2022). Additionally, ejection fraction, indicating the proportion of blood ejected from the ventricle during each contraction, changes in response to SLE-related factors. Fractional shortening, an index reflecting the contractile performance of the heart, also exhibits variations in individuals affected by SLE. These intricate adjustments collectively contribute to the complex cardiovascular manifestations observed in SLE (Deng et al. 2020).

Multimode mode (M-mode) images and the accompanying graphs (Figure 8 D) demonstrate that compared to control, PIL and W treated mice had considerably lower stroke volume, ejection fraction, fractional shortening, cardiac output, and left ventricular mass. Stroke volume and left ventricular mass were considerably lower in the Dex drug-treated group than in the control group; however, cardiac output, fractional shortening, and ejection fraction were significantly greater. In contrast all parameters (except fractional shortening and ejection fraction) in the DW treated group were identical to that of the healthy control (Figure 8 D).

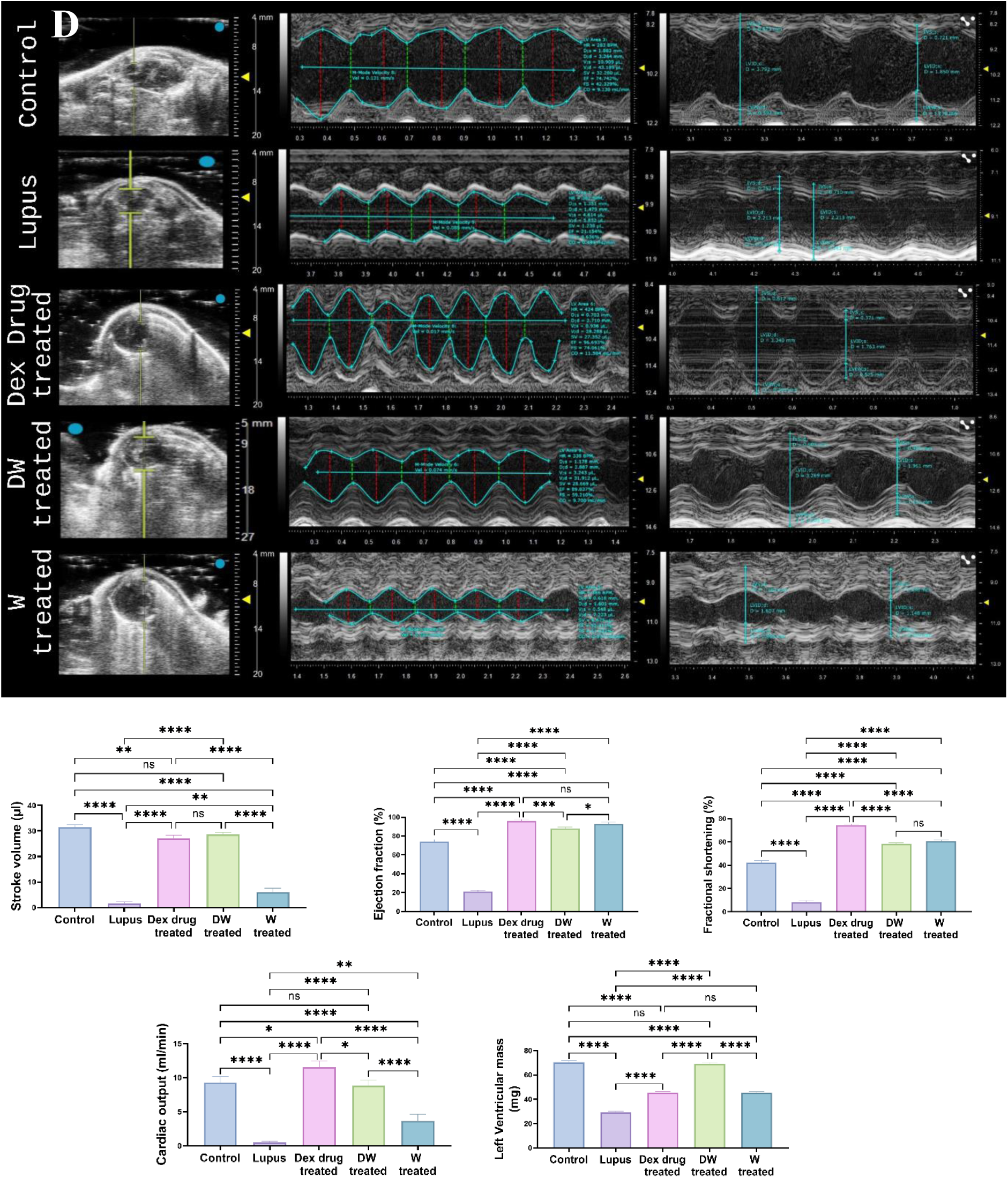
(D) The M-mode images show the LV AW, LV chamber, and LV PW throughout diastole (d) and systole (s). Echogenic peaks visible along the PW during systole represent the papillary muscle entering the field of view. The graph depicts the stroke volume, ejection fraction, fractional shortening, cardiac output and left ventricular mass in the heart of different treatment groups. Error bars show mean ± SD. p values indicate significant changes as follows: non-significant (ns) p > 0.05, *P < 0.05, **p < 0.01, ***P < 0.001 and ****P < 0.0001; One-way ANOVA. Rt., Right; Lt., Left; LVID, s, left ventricular internal diameter (systole); LVID, d, left ventricular internal diameter (diastole); SV, stroke volume; EF, ejection fraction; FS, fractional shortening; CO, cardiac output, and LV, left ventricle.

## Discussion

The *in vitro* and preclinical results obtained using DW in this study present a promising advancement in the field of CM based immunomodulation in autoimmunity. Most prevalent current treatments in SLE, majorly corticosteroids and immunosuppressants, have severe side effects and cause generalized immune suppression, warranting a need for a more effective therapy with minimal adverse effects. Studies have revealed that MSCs induce immunomodulation through their secretory factors and bioactive molecules like cytokines, growth factors, and chemokines (Yang et al., 2009). Various studies on stem cell-derived secreted factors have shown that they alone can cause immunomodulation (Weiss et al., 2019). The use of secretome-containing CM has several advantages compared to the use of stem cells, as CM can be manufactured, freeze-dried, packaged, and transported more easily. Being devoid of cells, there is no need to match the donor and recipient to avoid rejection problems. Therefore, stem cell-derived CM has promising prospects for the production of pharmaceuticals for immunomodulatory medicine. The priming of WJ-MSCs with Dex in this study and further analysis has provided insights into its immunomodulatory potential in SLE. Comparative studies with unprimed CM in parallel allowed for a direct comparison between the effects of Dex priming and unprimed secretory profile of WJ-MSCs. This helped elucidate the specific effects of Dex priming on the secretome of WJ-MSCs and its potential implications for therapeutic applications.

In this study the focus was on examining the immunomodulatory effects of DW treatment on various immune cell populations, including Tregs, Bregs, Th17, DN T cells, and inflammatory neutrophils in PBMCs from patients with SLE. DW treatment led to significant increase in the frequency of Tregs and Bregs, along with a reduction in TH17 cells, DN T cells, and inflammatory neutrophils were in concordance with the standard treatment outcomes in SLE. Significant increase observed in the frequency of various Treg and Bregs subtypes and levels of immunosuppressive cytokine IL-10 with DW treatment, indicated a restoration of immune balance and tolerance. The decrease in PD-1 expressing a B cells further suggested an encouraging role of DW in reduction in autoimmune responses and disease progression by promoting immune regulation. Additionally, the observed concurrent suppression of Th17 cells, DN T cells were attributed to the upregulation of IL-10 and downregulation of IL-17A production. The imbalance between Th17 cells and Tregs in SLE (Li et al., 2020) has been underscored, with Th17 cells being associated with inflammatory disorders. The increase in the population of DN T cells in PBMCs from SLE patients finds validation from earlier reports of high levels of DN T cells in SLE patients being associated with disease severity (Alexander et al., 2020) and also that the increased DN T cells act as a source of inflammatory IL-17 seen upregulated in SLE patients. (Crispín et al., 2008). Hence, achieving reduction in Th17 cell numbers and IL-17 with DW treatment in our study both *in vitro* and *in vivo* was an interesting finding. Correlation analysis revealed a positive correlation between Bregs and Tregs with DW treatment, indicating a coordinated regulatory response. Conversely, a negative correlation was observed with Th17 cells, DN T cells, and inflammatory neutrophils, highlighting the ability of DW treatment to restore immune homeostasis in SLE.

HCQ, a standard drug used for SLE (Dima et al., 2020) exhibited potent anti-inflammatory effects by inhibiting the release of pro-inflammatory cytokines, such as IL-17A, while concurrently promoting the release of anti-inflammatory cytokines, including IL-10 and TGF-β1. *In vivo* studies in MRL/lpr mice demonstrated HCQ’s ability to restore Th17/Treg balance, leading to a reduction in IL-17 expression in Th17 cells and an increase in Foxp3 and TGF-β levels (An et al., 2017). Comparison of the immunomodulatory effects of DW with standard drug HCQ, demonstrated that DW treatment exhibited similar effects as HCQ, with an increase in regulatory cells, decrease in Th17 cells, DN T cells, and neutrophils. Our investigation delves into the mechanisms underlying the impact of DW on the immune system, focusing on the TGF-β-mediated regulation and modulation of key immune cell populations. MSC-CM mostly mediates its suppressive effect through TGF-β, primarily when it pertains to inducing Tregs and preventing adaptive immunological responses (English et al., 2008) which was also observed for DW treatment using TGF-β inhibitor SB-431542. Findings of the inhibition studies as decreased populations of Tregs and Bregs and increase in Th17 cells, DN T cells, and neutrophils align with the hypothesis that DW contributes to Treg expansion via TGF-β secretion. Comparing the immunomodulatory effects of DW with standard therapies like HCQ and Dex drug, proposed similar efficacy of DW as that of HCQ in promoting regulatory cells and suppressing pro-inflammatory cells and the combination of DW with HCQ demonstrated enhanced suppression of Th17 cells and neutrophils than HCQ alone, suggesting a potential synergistic effect of DW. Interestingly, DW performed better than Dex drug in all immunomodulatory aspects.

As autoantibodies play a pivotal role in lupus pathogenesis, our study investigated the impact of DW on their production. The results revealed a significant decrease in the levels of anti-dsDNA and anti-ENA antibodies in the DW-treated group, suggesting potential attenuation of the autoimmune response. Additionally, DW treatment was associated with a significant increase in IL-10 and TGF-β cytokine levels, indicating an immunomodulatory shift towards anti-inflammatory responses. In contrast, Dex treatment resulted in an increase in autoantibodies and a decrease in IL-10 and TGF-β expression, highlighting the differential effects of Dex compared to DW. Examination of IL-17A and IL-10 gene expression further supports the immunomodulatory potential of DW. The treatment was effective in restoring the balance between elevated IL-10, TGF-β expression and reduced IL-17A.

In our *in vivo* studies using PIL mouse model, DW treatment demonstrated positive impact on overall health by reducing mortality, suppressing autoantibodies, managing proteinuria and maintaining body weight in different autoantibody-specific subgroups of the PIL model, highlighting the potential of DW in improving complete survival outcomes in lupus-like autoimmunity. DW treatment increased Treg and Breg populations while lowering Th17 cells, reversed the decrease in anti-inflammatory cytokine IL-10 production, and improved TGF-β levels, crucial for regulating T-cell immunity and promoting Tregs. Hence, the findings suggest that DW treatment in PIL mice exerts multi-faceted therapeutic effects by modulating immune cell populations and cytokine production, ultimately improving disease outcome. The study highlights and provide evidence for the efficacy of DW in reducing limb inflammation and treating alopecia, a dermatological symptom of lupus, by restoring vasculature and blood flow that is crucial for the reversal of alopecia. It also effectively resolved seizures in PIL mice, leading to progressive recovery within 7 days, highlighting its potential neuroprotective effects in addressing lupus-related neuropsychiatric manifestations.

In SLE, as a result of disease activity, damage to target organs and unfavourable events, renal involvement affects up to two-thirds of SLE patients, quite early in the course of the disease (Obrişcă et al., 2021). A range of vascular, glomerular, and tubulointerstitial lesions can be seen in the morphologic alterations in a kidney biopsy taken from a patient suffering from SLE (Weening et al., 2004). A concomitant condition often explains the vast morphological range seen in the liver of an SLE patient which includes fatty liver, portal inflammation, and vascular abnormalities like hemangioma, congestion, and arteritis (Grover et al., 2014). Immune-inflammatory process in SLE may have distinct pathogenetic processes that impact not just the lung parenchyma but also the pleural layers, respiratory muscles, and pulmonary vascular system (Palm et al., 2013). Commonly associated heart problems with lupus include pericarditis or pericardial effusion, valve abnormalities, myocarditis, arrhythmias, and accelerated atherosclerosis (Burkard et al., 2018). These organ malfunction are the primary cause of the patients’ morbidity and death. Hence, we studied gross morphological, histological, anatomical and functional changes in the respective organs. The observed improvements in vasculature, organ morphology and function in kidneys, liver, lungs, heart, and spleen with the DW treatment highlight its potential on reversing organ damage and histopathological changes in a murine model of SLE. DW helped in mitigating pathological changes and abnormalities associated with SLE.

The results of this study provide valuable insights into the impact of DW treatment on renal, hepatic, and cardiac parameters in a murine model of SLE. Imaging modalities, including B-mode, M-mode, color Doppler, US, and photoacoustic imaging, have been instrumental in characterizing anatomical and functional changes in the kidneys, liver, and heart. Interestingly, a significant improvement in renal, hepatic and cardiac oxygenation as well as hemoglobin concentration along with improved compromised organ function was observed with DW treatment, strongly suggesting its potential therapeutic effect.

Overall, through a combination of *in vitro* experiments, murine lupus model studies, and advanced imaging techniques, our research demonstrated the multifaceted immunomodulatory benefits of DW treatment on various aspects of lupus pathophysiology. Notably, DW exhibited significant immunomodulatory effects by expanding regulatory B-,T-cell subtypes and suppressing pathogenic immune cell populations from patients with SLE *in vitro*. Comparative analyses with HCQ, a standard SLE treatment, revealed that DW shares similar immunomodulatory effects with HCQ, suggesting comparable efficacy. In a PIL mouse model, DW showed therapeutic efficacy by improving survival rates, suppressing autoantibody production, and modulating immune cell population. Advanced imaging highlighted organ-protective effects on the kidneys, liver, and heart, while the reversal of alopecia and anti-inflammatory effects were added to DW’s comprehensive therapeutic potential. These findings strongly indicate DW as a potentially effective SLE-ameliorating strategy, either used in combination with a standard drug or alone warranting further exploration and potential for clinical development, to validate its efficacy and safety in human subjects.

## Materials and methods

### Isolation, expansion and characterization of WJ-MSCs

#### Isolation and expansion of WJ-MSCs

Umbilical cord derived WJ-MSCs were collected and processed within 24 h of normal or cesarean delivery from the Department of Obstetrics and Gynecology, AIIMS, New Delhi. Ethical clearance was obtained from the Institutional Committee for Stem Cell Research, All India Institute of Medical Science, Delhi for the collection of MSCs derived conditioned and pre-conditioned media (Ref no. : IC-SCR/116/20(R), Dated: 14.06.2021). Briefly, the umbilical cord was collected in a 50 ml schott bottle containing phosphate buffered saline (PBS) with 1% antibiotics (Penicillin, Streptomycin and Gentamycin). Upon arrival, the samples were extensively washed with PBS containing 1% antibiotics. The artery part of the cord was exposed using a sharp surgical blade and chopped into smaller pieces (approx. ∼2mm). The exposed jelly part of the cord was placed in a 35 mm culture dish and kept undisturbed. The cultures were incubated overnight at 37°C with 1 ml complete medium in 5% CO_2_ that was changed every three-four day. When the cells started growing and migrating out of the explant and reached 80% confluence, they were harvested using 0.05% trypsin-EDTA (In*vitro*gen-Gibco) and transferred into a 60 mm culture dish.

#### Characterization of WJ-MSCs

##### Morphological analysis of cultured WJ-MSCs

Cell cultures were monitored using phase-contrast microscopy (Olympus) to evaluate cell morphology and confluency. All assays were performed using WJ-MSCs at passages three and five.

##### Measurement of Metabolic Activity by MTT assay

The proliferation rate of MSCs (n=3) was measured on days 1, 3, 5, 7, and 14 using 3-(4, 5-Dimethylthiazol-2-yl)-2,5-diphenyltetrazolium bromide (MTT) (5 mg/ml HiMedia). After four hours of addition of MTT, 0.04M HCl isopropanol was added to the medium and incubated at 37°C for 1h in dark. The absorbance was measured at 570 nm using a 96 well microplate reader (Synergy^TM^ HT, Bio-Tek Instruments, Inc.).

##### Immunophenotyping for MSCs

MSCs were characterized at passage three for surface markers using flow cytometry before being used for any experimental analysis. Briefly, the cells were incubated with the labelled antibodies at room temperature (RT) in the dark for 1 h. The following anti-human antibodies were used: CD73 (PE; Becton Dickinson), CD90 (PECy5; Becton Dickinson), HLA Class I (APC; Becton Dickinson), HLA Class II (FITC; Becton Dickinson), CD29 (FITC; eBioscience) and CD105 (APC; eBioscience). After incubation, the cells were washed twice with flow staining buffer, centrifuged at 800×g for 5 min, and the final cell pellet was resuspended in 300μl of cold flow staining buffer. Unlabeled cells were used as experimental controls. An isotype control was included in each experiment and specific staining was measured from the cross point of the isotype with a specific antibody graph. Cell fluorescence was evaluated using a BD FACS LSR II (Becton Dickinson) instrument, and data were analyzed using BDFACS DIVA software.

#### Trilineage differentiation

##### Osteogenic Differentiation of WJ-hMSCs

MSCs at third passage and 70-80% confluency was exposed to osteogenic differentiation medium containing 1X DMEM-LG (GIBCO, Invitrogen), 10% FBS (Hyclone), 50 μM ascorbic acid-2-phosphate, 0.1 μM dexamethasone and 10 mM ß-glycerophosphate (Sigma) for 4 weeks with intermittent change of media every 2 days. Uninduced hMSCs were used as the experimental controls. Differentiation was confirmed by Alizarin Red S staining (HiMedia).

##### Adipogenic Differentiation of WJ-hMSCs

MSCs in the third passage were exposed to adipogenic differentiation media containing 1X DMEM-LG, 100 μM indomethacin, 1 μmol/L dexamethasone, 500 μM 3-isobutyl-1-methylxanthine, 1 μg/ml insulin (Sigma), and 10% FBS (Hyclone) for 21 days. Uninduced MSCs were used as experimental controls. The medium was changed every three days. Differentiation was confirmed using Oil Red O staining (HiMedia).

##### Chondrogenic Differentiation of WJ-hMSCs

A commercially available kit (GIBCO, Invitrogen) was used for chondrogenic differentiation. Briefly, a single cell suspension of third passage MSCs was prepared at a concentration of 1.6×10^7^ viable cells/ml using TrypLE Express to detach cells from the monolayer. Micromass culture was generated by seeding 1X10^5^ cells/10 μl of cell solution in a 35 mm tissue culture plate and incubated for 2 h at 37°C with 5% CO_2_. After this, chondrogenic differentiation medium was added gently, without disturbing the cell micromass. The medium was changed every 3^rd^ day for 14 days before termination. Uninduced MSCs were used as the experimental controls. Differentiation was confirmed using Alcian Blue staining (Spectrochem).

#### Preconditioning of MSCs with Dex

Harvested cells were incubated in serum-free medium containing Dex (Sigma Aldrich) (3000 ng/ml; Sigma-Aldrich). Dex-preconditioned media were collected after 24 h of preconditioning.

### Human population and collection of samples

The study subjects were recruited from Sir Sunderlal Hospital, Banaras Hindu University (BHU), Varanasi, Uttar Pradesh, India. Ethical clearance was obtained from the Institutional Ethics Committee of the Institute of Medical Science, BHU (Ref. Dean/2020/ EC/Ro32, dated: 27.06.2020) for the collection of SLE patients (n=74) and related studies and provided written informed consent for participation in this study. All participants were deemed systemically SLE patient based on their detailed medical history and selected laboratory blood work. Detailed information regarding age, sex, medical history, symptoms, medications, and clinical manifestations was collected on a predesigned proforma. The inclusion criteria for this study were as follows: age ≥ 18 years, SLE or suspected SLE established by ACR criteria, and glomerulonephritis and pericarditis. The exclusion criteria were as follows: history of hepatitis B or C, history of HIV, cancer, pregnancy or lactation, diagnosis of diabetes and/or HbA1C level >6%, and any comorbidity of medical, psychological/psychiatric condition, or treatment.

### Isolation and culture of peripheral blood mononuclear cells (PBMCs)

Blood (10 ml) from SLE patients was collected in heparinized tubes. PBMCs were isolated by density gradient centrifugation using a HiSep™ LSM 1077 (Himedia). All procedures were performed at RT. The whole anticoagulated blood was mixed with an equal volume of sterile PBS solution in a 50 ml conical tube. HiSep was pipetted into a separate 15 ml conical tube onto which PBS diluted blood was gently layered on one third of it by volume. The sample was centrifuged at 400 g for 30 min with no brake at RT (25°C). Most of the plasma and platelet-containing supernatants were discarded, and mononuclear cells above the interface band (granulocytes and erythrocytes) were aspirated and transferred to a separate tube. Isotonic PBS (10 ml) was added to the mononuclear cell layer in a centrifuge tube and mixed via gentle aspiration. The mixture was centrifuged at 200 g with brake off at RT for 10 min. Washing with isotonic PBS removed the HiSep LSM and reduced the number of platelets. The cells were washed again with isotonic PBS and resuspended in RPMI medium.

### Immunomodulatory impact of W and DW

Isolated PBMCs (1×10^6^ cells/well) were incubated with 40% non-preconditioned MSC-derived conditioned media (W) or 40% Dex (7.6 µM) primed preconditioned media (DW) for 24 h. Cells incubated only in RPMI media served as controls. Further, the cell suspensions were stimulated with 2 µg/mL cytosine-phosphate-guanosine oligodeoxynucleotide (CpG ODN; InvivoGen), 50 ng/ml phorbol myristate acetate (PMA; InvivoGen), and 1 μg/ml ionomycin (InvivoGen) during the last 6 h of culture.

### Comparison of Immunomodulatory potential of DW and HCQ

To compare the immunomodulatory effect of DW with standard drug HCQ and of their combination, 1×10^6^ PBMCs were incubated with 7.6 µM HCQ (Sigma-Aldrich), Dex drug (7.6 µM), DW, and W for 24 h and stimulated as described above.

### Inhibition study

SB 431542 hydrate (Sigma Aldrich) was used as an inhibitor of TGF-β activity to study the functional relevance of TGF-β in DW mediated immunomodulation. For this, PBMCs were incubated with SB 431542 hydrate at a concentration of 5 µM, then after Dex drug, DW and W was added for 24 h and stimulated with CpG ODN, PMA and ionomycin for last 6 h.

### Immunophenotyping for SLE associated Immune Cells

#### Cell surface staining

For flow-cytometric analysis, PBMCs were extracted from peripheral blood samples of patients with SLE and co-incubated with either Dex drug, DW or W. To assess cell viability PBMCs were stained with Fixable Viability Stain (BD Biosciences). For cell staining PBMCs were suspended in PBS containing 0.1% sodium azide (NaN_3_) and 1% bovine serum albumin (BSA). For immunophenotyping of Tregs cells (1×10^6^/sample) PBMCs were stained with CD4 (FITC; BD Biosciences), CD25 (PE; eBioscience), CD127 (APC; eBioscience), FOXP3 and for Bregs immunophenotyping, CD19 (APC; eBioscience), CD24 (FITC; eBioscience), and CD27 (PE; eBioscience) antibodies were used to stain for surface antigens for 30 min at 4°C in dark. DN T cells were stained with antibodies against surface markers CD3 (APC; BD Biosciences), CD4 (FITC; BD Biosciences), and CD8 (PE; BD Biosciences). Neutrophils were stained for surface markers CD3 (APC; BD Biosciences) and CD15 (FITC; BD Biosciences). CD4^+^ T cells, CD19^+^ B cells, Th17 cells, DN T cells, and neutrophils were also stained with PD1 antibody (PerCP; BD Biosciences) for 30 min at 4°C in the dark. Following incubation, cells were stored in 1% paraformaldehyde (PFA, Sigma-Aldrich) at 4°C in the dark until they were acquired by flow cytometer (CytoFLEX, Beckman Coulter). CytExpert software (Beckman Coulter) was used for data analysis.

#### Intracellular staining

Intracellular staining was done for FOXP3 (PE; eBiosciences), IL-10 (PerCP; eBiosciences) and IL-17A (PE; eBiosciences). Briefly, Brefeldin A (5 ng/ml) was added to the PBMCs cultures during last 4 h of culture. The culture supernatant was isolated and stored until further use. Then after, the cells were washed with washing buffer [PBS, BSA (1%) and NaN_3_ (0.1%)], permeabilized with FACS Permeabilizing Solution (BD Biosciences) for 10 min according to manufacturer’s protocol and stained with respective antibodies. Finally, cells were fixed with PFA; 1% and stored in the dark at 4°C until further use. The samples were acquired using a CytoFLEX cytometer (Beckman Coulter). Data analysis was performed using the CytExpert software (Beckman Coulter).

### IL-10 and TGF-β1 Immunoassay

The supernatant of W or DW treated PBMCs from SLE patients were assayed for the active form of TGF-β1 using a Human TGF-β1 specific ELISA kit (Elabscience), as per the manufacturer’s protocol. This immunoassay system was designed for the sensitive and specific detection of biologically active TGF-β1. The antibody in this system did not recognize the TGF-β1 precursor. To determine total TGF-β1 in the cell supernatant, samples were pretreated with 1N HCl for 15 min at RT before neutralization with 1N NaOH, as suggested by the manufacturer. This procedure converted latent TGF-β1 to its active form.

### Detection of autoantibodies

Autoantibodies against dsDNA and ENA in supernatant of DW or W treated PBMCs from SLE patients were determined using ELISA (MyBioSource) as per manufacturer’s instruction.

### Quantitative real time PCR (qRT PCR)

IL-10 and IL-17A gene expressions in Dex drug, DW or W treated PBMCs from SLE patients were quantified using qRT PCR. Total RNA was extracted using Trizol Reagent (Invitrogen) and phenol/chloroform extraction method. RNA concentration was quantified using a NanoDrop spectrophotometer (ThermoScientific) and RNA integrity was tested by electrophoresis on a 1.5% agarose gel. One-step qRT PCR (Applied Biosystems) was carried out according to the manufacturer’s recommendations. The results were normalized to those of the β-actin control and are reported as ΔCT values and fold changes. Relative gene quantification was performed using the 2^−*ΔΔ*Ct^ method following normalization to β-actin in the respective groups. The primer sequences used were as follows:

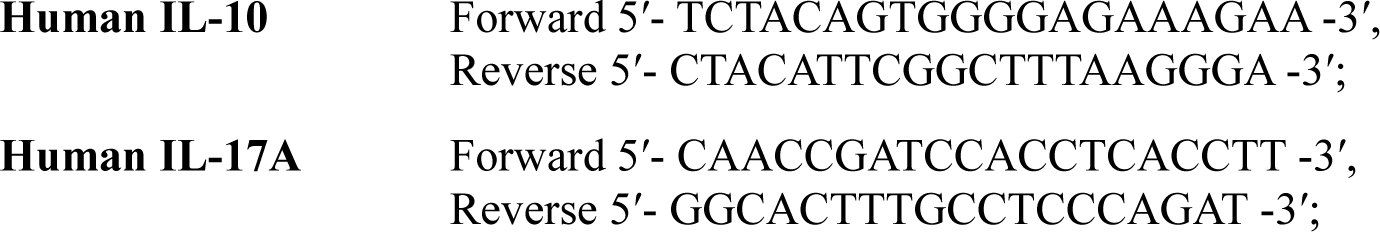

The housekeeping gene β-actin was used as an endogenous control,

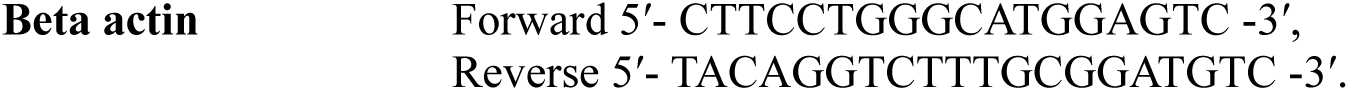

### Pre-Clinical studies using PIL Model

#### Sex as a biological variable

Our study exclusively examined female mice because the disease modeled is only relevant in females.

#### Mice

Female BALB/c mice (n=50, 8 weeks old, 20 ± 2 g) were purchased from the Institute of Medical Science, BHU (Varanasi, India). The animals were maintained in a barrier system with an alternating 12 h light/dark cycle, a relative humidity of 50 ± 5%, and a constant temperature of 25°C. All experimental protocols involving the care and use of animals were reviewed and approved by the Institutional Animal Ethics Committee (IAEC) of the Institute of Science, BHU, India (Ref no.: BHU/DoZ/IAEC/ 2022-2023/003).

Mice (n=40) were administered an intraperitoneal injection of pristane (0.5 ml) (Sigma-Aldrich). Following 4 weeks of pristane treatment, development of PIL model was determined by measuring the generation of anti-dsDNA and anti-ENA autoantibodies. The PIL mice were randomly divided into four groups (n=10 per group) and, intraperitoneal injections of various treatments (1.3 ml) were administered daily for next 6 weeks (group 1: No treatment Lupus group (normal saline); group 2: Dex drug treated, group 3: DW treated, 4: W treated and healthy mice (n=10) were used as control and administered normal saline (1.3 ml) for 6 weeks.

#### Determination of autoantibodies

For detection of anti-dsDNA and anti-ENA autoantibodies post pristane injection, blood was collected from the tail vein at 4 and 16 weeks, and serum was isolated. Using ELISA kits (MyBioSource) the serum levels of IgG autoantibodies against dsDNA and ENA were measured following the manufacturer’s instructions.

#### Body weight, renal function and lifespan analysis

For every mouse, body weight (in grams) was measured at day 7, 14, 21, 28, and 35 after lupus induction. Urine was collected on day 7, 14, 21, 28, and 35 from the control, and the four PIL groups of mice in the morning into a clean microcentrifuge tube. Urine was dropped into a urine analysis strip (Uristix, Siemen) and analyzed according to the manufacturer’s instructions. All groups described above were closely monitored for 35 weeks after lupus induction. To calculate the percent survival rate, Kaplan-Meier survival analysis was performed, and the animals were euthanized when moribund.

#### Immunophenotyping for disease associated immune cells

##### Mice splenocyte preparation

Single-cell suspensions of the spleen were prepared and collected in 10 ml of staining buffer in a conical tube and filtered through a cell strainer to remove any remaining debris or clumps. After centrifugation for 5 min (400×g) at 4°C the supernatant was removed and RBC lysis was performed. The remaining splenocytes were reconstituted in staining buffer, and live cell counting was performed using the Trypan Blue exclusion assay.

##### Flow cytometry analysis

###### Cell surface staining

For flow-cytometric analysis, cells prepared from the spleen of different PIL groups and control mice were used for cell staining by suspending them in PBS containing 0.1% NaN_3_ and 1% BSA. For immunophenotyping of Tregs cells (1×10^6^/sample) splenocytes were stained with CD4 (PerCP; BioLegend), CD25 (APC; BioLegend) and for Bregs immunophenotyping, CD5 (PerCP; BioLegend) and CD1d (PE; BioLegend) antibodies were used to stain for surface antigens for 30 min at 4°C in dark. Following incubation, cells were stored in 1% PFA (Sigma-Aldrich) at 4°C in the dark until they were acquired by flow cytometer (CytoFLEX, Beckman Coulter). CytExpert software (Beckman Coulter) was used for data analysis.

###### Intracellular staining

Intracellular staining was done for FOXP3 (PE; BioLegend) and IL-17A (FITC; BioLegend). Then after, the cells were washed with washing buffer, permeabilized with FACS Permeabilizing Solution (BD Biosciences) for 10 min according to manufacturer’s protocol and stained with respective antibodies. Finally, cells were fixed with PFA; 1% and stored in the dark at 4°C until further use. The samples were acquired using a CytoFLEX cytometer (Beckman Coulter). Data analysis was performed using the CytExpert software (Beckman Coulter).

#### ELISA Analysis

The venous blood samples were allowed to clot at RT for 15–30 min, serum separation was done by centrifugation at 10,000×g for 10 min. Serum levels of IL-10 and TGF-β were measured using ELISA kits (Elabscience) according to the manufacturer’s instructions. The cytokine concentrations were quantified using standard curves.

#### Clinical assessment in PIL mice

##### Alopecia

Mice with lupus exhibited progressive dorsal alopecia, and the afflicted areas of skin were very dry and lost pliability. Following PIL induction, they were randomly divided into four groups as described in 3.2.3.1 and received treatments during the following 42 days. After observing hair loss, the marked site of alopecia was observed for 42 days for hair regrowth.

For *in vivo* imaging, Fujifilm VisualSonics’ Vevo 3100 system was utilised, and all images were obtained using an MX-550D linear-array transducer (25–55 MHz centre, 15 mm scan depth). The mice were scanned while lying supine. The dermal layer was seen by positioning and holding stationary the RMV-706 scanhead utilising VisualSonics Vevo Integrated Rail System II and B-mode imaging. At the site of hair loss, an ultrasonic transducer (Fujifilm Visual Sonics, Inc.) with a jacket that contained a short (14 mm) optical fibre bundle was placed. A transparent ultrasonic gel (OXD, Spain) without bubbles was utilised to seal the 5 mm space that existed between the mouse skin and the transducer surface. For PAI, the following imaging parameters were chosen: PA gain = 37 dB, power 100%, step size 0.219 mm, and high sensitivity were maintained. The rats were sedated with a 2% isoflurane induction dosage and a 1.5–2% maintenance dose prior to imaging. After that, the animals were put to sleep in the prone position on a table that was kept at 37^◦^C for the purpose of performing PAI and ultrasonography. A baseline recording of the dermal layer’s power Doppler blood flow was made after a brief time of stabilisation. All images were processed using the Vevo LAB software (FUJIFILM VisualSonics).

##### Paw swelling and inflammation

After lupus induction and treatment with preconditioned and non-preconditioned media, mice were assessed for paw edema and redness in the forelimb and hindlimb six weeks after pristane injection. Animals were observed blindly for swelling and redness by examining the inflammatory joints in each paw; swelling and redness are indicators of inflammation. Unaware of the identities of the mice, two investigators (specialists in anatomical pathology) conducted blinded clinical examinations.

##### Seizure

Seizure development was monitored and video graphed. The seizure-prone mice were continually monitored for improvement.

#### Measurement of size and weight of organ

Mice were used as experimental subjects, and all procedures were conducted in compliance with ethical guidelines approved by the IAEC. At week 35 (5 weeks following treatment), euthanasia was carried out using an approved method to minimize suffering, and organs were harvested to measure the size and weight to assess the effect of Dex, preconditioned, and non-preconditioned media. The mice were placed in a supine position for optimal access to abdominal organs. Abdominal organs, including the kidneys, liver, lungs, heart, and spleen, were dissected using sterile surgical instruments. Each organ was placed in a clean labelled Petri dish. Organ length (cm) was measured using a ruler. With the organ gently held, measurements were taken from one end to the other, ensuring an accurate recording of each length.

A precision balance was used to separately weigh each organ. Prior to each measurement, the balance was calibrated and a Petri dish was placed and tared to ensure accuracy. Organ weight was recorded by placing the organ on a Petri dish on a balance and noting the measurement. For the spleen, the splenic index was calculated to assess the relative organ size compared with the overall body weight. Splenic index was determined by dividing the weight of the spleen by the body weight of the mouse and multiplying the result by 100.

#### Histological Analysis

Standard histological procedures were followed for the processing, embedding, sectioning, and staining of different organs. Furthermore, the cells were analyzed and evaluated for cellular composition, organization, and identification of any abnormalities or pathological changes. At the conclusion of the experimental period, the mice were euthanized following approved procedures. Organs, including the kidneys, liver, lungs, heart, and spleen, were carefully dissected from each mouse. Freshly harvested organs were immediately fixed in a suitable fixative solution (10% formalin) to preserve cellular structures. The fixed organs were then embedded in paraffin. Paraffin embedding was performed according to the established protocols to ensure proper tissue infiltration. Embedded organs were sliced into thin sections with a thickness of approximately 5 μm using a microtome. Sections were collected on glass slides for histological analysis. Thin rehydrated tissue sections were stained with H & E to enhance cellular detail and highlight the tissue structures. Staining was conducted following standard protocols (Gunawan et al., 2017). Blinded evaluation of the stained sections was performed to minimize bias. The sections were coded and randomized before evaluation to ensure objectivity. Morphological changes and inflammatory responses were assessed based on established criteria. Slides were observed and images were obtained using a live-cell imager fluorescence microscope (EVOS FL Cell Imaging System, Life Technologies, USA).

#### Organ Damage Assessment using PAI

Abdominal and cardiac region images were obtained in live mice using high-resolution US imaging using a Vevo3100 LAZR-X small animal US in conjunction with a PAI system equipped with an MX-550D (25-50 MHz, scan depth of 15 mm) and 250S (15-30 MHz, scan depth of 30 mm) linear-array transducer. A depilator and hair removal lotion were used to remove the hair from the abdomen, and alcohol pads were used to properly clean the scalp. A transparent, bubble-free ultrasonic gel was used to bridge the 5 mm gap that existed between the mouse skin and transducer surface. For PAI, the following imaging settings were used: power 100%, step size 0.279 mm, PA gain = 37 dB, and high sensitivity. The mice were given anesthesia with a 2% induction dose of isoflurane and a 1.5–2% maintenance dose prior to imaging. After that, the animals were placed supine on an operating table kept at 37^◦^C so that the ultrasonography and PAI could be carried out on a heated platform while the animals’ body temperatures, heart rates, and respiration rates were tracked. B-mode imaging was one of the main imaging techniques used to see a cross-sectional image of the internal anatomy of the kidneys, liver, and heart in real time. The size, shape, and location of the organ may also be seen in this mode. Furthermore, the method made it possible to create a 3D model of the kidney, which allowed for a more thorough visualisation and comprehension of its structure. To gather the area; 2D data, depth volume (3D), and blood flow in the relevant organ, a 3D-mode overlay over the power Doppler image was utilised. It was possible to evaluate the blood flow in the kidneys, liver, and heart by using colour Doppler imaging. The oxygen saturation was measured using the following settings once the system was initialised: depth, 21–29 mm; width, 13.75 mm; and wavelength, 750 and 850 nm for the total Hbt and sO2, respectively. The probe was positioned precisely over the abdomen to obtain all of the photographs. The probe was positioned precisely over the abdomen to obtain all of the photographs. The PAI system’s image processing software was then used to determine the mean value of the PA signals in each B-mode frame, from which the total PA signals was derived.

A Vevo LAZR-X Imaging System (Fujifilm VisualSonics Inc., Toronto, Canada) was used to do echocardiography. A high-frequency linear array transducer that is appropriate for imaging tiny animals was installed in the imaging system. One day before the experiments, the mice’s anterior chests were depilated. The mice were anaesthetized with 2.5–3.0% isoflurane at a flow rate of 0.8 L/min and maintained with 1-2.5% isoflurane on the day of image capture. The mice were then placed supine on a pad with ECG electrodes, a temperature sensor, and a heater. Following anaesthesia, it was made sure that the temperature remained at 37°C and that the maximum heart rate was 500 beats per minute. To provide the best possible acoustic connection between the transducer and skin, US gel was placed to the chest. Throughout the imaging procedure, echocardiographic data, such as heart rate, were continually observed. Initially, 2D imaging was used to evaluate overall heart function and visualise the cardiac architecture. Measurements of cardiac structures, including stroke volume, ejection percent, fractional shortening, cardiac output, and left ventricular mass, were obtained using M-mode imaging. To measure blood flow patterns and velocities, Doppler imaging—including colour and pulsed-wave Doppler imaging—was used. Aortic and mitral flow velocities, among other characteristics, were measured using pulsed-wave Doppler. Pulsed-wave Doppler was used to measure blood flow velocities during both systole and diastole. The systolic and diastolic velocities were measured and averaged across a number of cardiac cycles. Using the M-mode, the left ventricle’s systolic diameter was determined by placing the cursor perpendicular to the ventricular walls during systole. During diastole, the diastolic diameter was also measured in M-mode. The aortic valve’s peak diastolic and systolic velocities were measured using pulsed-wave Doppler. Doppler and M-mode echocardiographic images were recorded and stored for subsequent offline analysis using Vevo LAB software.

#### Statistical analysis

Statistical analyses were performed using Prism (GraphPad version 10.1.2). The mean ± SEM of at least three independent experiments was calculated for all the experiments. The unpaired two-tailed Student’s t-test was used to compare the means between two independent groups, one-way ANOVA and two-way ANOVA was used to compare the means between two or more groups followed by Tukey’s multiple comparison post-hoc test. Specific P values are detailed in the figure legends. Correlations were assessed using the Spearman’s rank correlation coefficient. Differences were considered significant when p-values were < 0.05.

## Supporting information

Isolated WJ-MSCs formed a homogenous monolayer of adherent, spindle-shaped cells and exhibited proliferation capacity of WJ-MSCs

. Mice that received normal saline died on the third day of treatment, whereas the DW treated mice recovered progressively from seizures in 7 days

. Mice that received normal saline died on the third day of treatment, whereas the DW treated mice recovered progressively from seizures in 7 days

. Mice that received normal saline died on the third day of treatment, whereas the DW treated mice recovered progressively from seizures in 7 days

## Data availability

All supporting data are provided in the Supporting Data Values file.

## Author contributions

GR conceptualized and designed the study. KP performed the experiments, and, GR and KP analyzed the data, drafted the manuscript. SR and SM provided the conditioned media, performed and analysed the characterization of mesenchymal stem cells. KG provided the animal housing facility had helped with the animal handling. DD, HT and SS critically reviewed and revised the manuscript. MC and MR provided SLE patients’ blood samples for the study and did clinical analysis. GR critically reviewed and revised the manuscript, provided overall guidance for the project and approved the final manuscript as submitted. All authors approved the final manuscript as submitted and agree to be accountable for related aspects of the work.

## Acknowledgments

The authors sincerely acknowledge Interdisciplinary School of Life Sciences (ISLS), Interdisciplinary School of Life Sciences (SATHI) and Central Discovery Center (CDC), BHU, Varanasi, India for equipment support. This study was supported by Science and Engineering Research Board, India (CRG/2022/002561/BHS).

## Notes

### Competing Interest Statement

The authors have declared no competing interest.

## References

1. Alexander JJ, Jacob A, Chang A, Quigg RJ, Jarvis JN. Double negative T cells, a potential biomarker for systemic lupus erythematosus. Precis Clin Med. 2020 Mar;3(1):34–43. doi: 10.1093/pcmedi/pbaa001. Epub 2020 Jan 20. PMID: 32257532; PMCID: PMC7093895.

2. Alghareeb R, Hussain A, Maheshwari MV, Khalid N, Patel PD. Cardiovascular Complications in Systemic Lupus Erythematosus. Cureus. 2022 Jul 8;14(7):e26671. doi: 10.7759/cureus.26671. PMID: 35949751; PMCID: PMC9358056.

3. An N, Chen Y, Wang C, Yang C, Wu ZH, Xue J, Ye L, Wang S, Liu HF, Pan Q. Chloroquine Autophagic Inhibition Rebalances Th17/Treg-Mediated Immunity and Ameliorates Systemic Lupus Erythematosus. Cell Physiol Biochem. 2017;44(1):412–422. doi: 10.1159/000484955. Epub 2017 Nov 14. PMID: 29141242.

4. Bensreti H, Alhamad DW, Gonzalez AM, Pizarro-Mondesir M, Bollag WB, Isales CM, McGee-Lawrence ME. Update on the Role of Glucocorticoid Signaling in Osteoblasts and Bone Marrow Adipocytes During Aging. Curr Osteoporos Rep. 2023 Feb;21(1):32–44. doi: 10.1007/s11914-022-00772-5. Epub 2022 Dec 24. PMID: 36564571; PMCID: PMC9936962.

5. Bertsias GK, Tektonidou M, Amoura Z, Aringer M, Bajema I, Berden JH, Boletis J, Cervera R, Dörner T, Doria A, Ferrario F, Floege J, Houssiau FA, Ioannidis JP, Isenberg DA, Kallenberg CG, Lightstone L, Marks SD, Martini A, Moroni G, Neumann I, Praga M, Schneider M, Starra A, Tesar V, Vasconcelos C, van Vollenhoven RF, Zakharova H, Haubitz M, Gordon C, Jayne D, Boumpas DT; European League Against Rheumatism and European Renal Association-European Dialysis and Transplant Association. Joint European League Against Rheumatism and European Renal Association-European Dialysis and Transplant Association (EULAR/ERA-EDTA) recommendations for the management of adult and paediatric lupus nephritis. Ann Rheum Dis. 2012 Nov;71(11):1771–82. doi: 10.1136/annrheumdis-2012-201940. Epub 2012 Jul 31. PMID: 22851469; PMCID: PMC3465859.

6. Blanco S, Bandiera R, Popis M, Hussain S, Lombard P, Aleksic J, Sajini A, Tanna H, Cortés-Garrido R, Gkatza N, Dietmann S, Frye M. Stem cell function and stress response are controlled by protein synthesis. Nature. 2016 Jun 16;534(7607):335–40. doi: 10.1038/nature18282. PMID: 27306184; PMCID: PMC5040503.

7. Burkard T, Trendelenburg M, Daikeler T, Hess C, Bremerich J, Haaf P, Buser P, Zellweger MJ. The heart in systemic lupus erythematosus - A comprehensive approach by cardiovascular magnetic resonance tomography. PLoS One. 2018 Oct 1;13(10):e0202105. doi: 10.1371/journal.pone.0202105. PMID: 30273933; PMCID: PMC6167090.

8. Chang CP, Chio CC, Cheong CU, Chao CM, Cheng BC, Lin MT. Hypoxic preconditioning enhances the therapeutic potential of the secretome from cultured human mesenchymal stem cells in experimental traumatic brain injury. Clin Sci (Lond). 2013 Feb;124(3):165–76. doi: 10.1042/CS20120226. PMID: 22876972.

9. Chen W, Jin W, Hardegen N, Lei KJ, Li L, Marinos N, McGrady G, Wahl SM. Conversion of peripheral CD4+CD25-naive T cells to CD4+CD25+ regulatory T cells by TGF-beta induction of transcription factor Foxp3. J Exp Med. 2003 Dec 15;198(12):1875–86. doi: 10.1084/jem.20030152. PMID: 14676299; PMCID: PMC2194145.

10. Cook L, Reid KT, Häkkinen E, de Bie B, Tanaka S, Smyth DJ, White MP, Wong MQ, Huang Q, Gillies JK, Ziegler SF, Maizels RM, Levings MK. Induction of stable human FOXP3+ Tregs by a parasite-derived TGF-β mimic. Immunol Cell Biol. 2021 Sep;99(8):833–847. doi: 10.1111/imcb.12475. Epub 2021 Jun 3. PMID: 33929751; PMCID: PMC8453874.

11. Crispín JC, Oukka M, Bayliss G, Cohen RA, Van Beek CA, Stillman IE, Kyttaris VC, Juang YT, Tsokos GC. Expanded double negative T cells in patients with systemic lupus erythematosus produce IL-17 and infiltrate the kidneys. J Immunol. 2008 Dec 15;181(12):8761–6. doi: 10.4049/jimmunol.181.12.8761. PMID: 19050297; PMCID: PMC2596652.

12. Deckers J, Bougarne N, Mylka V, Desmet S, Luypaert A, Devos M, Tanghe G, Van Moorleghem J, Vanheerswynghels M, De Cauwer L, Thommis J, Vuylsteke M, Tavernier J, Lambrecht BN, Hammad H, De Bosscher K. Co-Activation of Glucocorticoid Receptor and Peroxisome Proliferator-Activated Receptor-γ in Murine Skin Prevents Worsening of Atopic March. J Invest Dermatol. 2018 Jun;138(6):1360–1370. doi: 10.1016/j.jid.2017.12.023. Epub 2017 Dec 27. PMID: 29288652; PMCID: PMC7611015.

13. Deng W, Xie M, Lv Q, Li Y, Fang L, Wang J. Early left ventricular remodeling and subclinical cardiac dysfunction in systemic lupus erythematosus: a three-dimensional speckle tracking study. Int J Cardiovasc Imaging. 2020 Jul;36(7):1227–1235. doi: 10.1007/s10554-020-01816-6. Epub 2020 Mar 19. PMID: 32193773.

14. Dima A, Jurcut C, Chasset F, Felten R, Arnaud L. Hydroxychloroquine in systemic lupus erythematosus: overview of current knowledge. Ther Adv Musculoskelet Dis. 2022 Feb 14;14:1759720X211073001. doi: 10.1177/1759720X211073001. PMID: 35186126; PMCID: PMC8848057.

15. Duijvestein M, Molendijk I, Roelofs H, Vos AC, Verhaar AP, Reinders ME, Fibbe WE, Verspaget HW, van den Brink GR, Wildenberg ME, Hommes DW. Mesenchymal stromal cell function is not affected by drugs used in the treatment of inflammatory bowel disease. Cytotherapy. 2011 Oct;13(9):1066–73. doi: 10.3109/14653249.2011.597379. Epub 2011 Aug 17. PMID: 21846292.

16. English K, Barry FP, Mahon BP. Murine mesenchymal stem cells suppress dendritic cell migration, maturation and antigen presentation. Immunol Lett. 2008 Jan 15;115(1):50–8. doi: 10.1016/j.imlet.2007.10.002. Epub 2007 Oct 31. PMID: 18022251.

17. Fontaine J, Chagnon-Choquet J, Valcke HS, Poudrier J, Roger M; Montreal Primary HIV Infection and Long-Term Non-Progressor Study Groups. High expression levels of B lymphocyte stimulator (BLyS) by dendritic cells correlate with HIV-related B-cell disease progression in humans. Blood. 2011 Jan 6;117(1):145–55. doi: 10.1182/blood-2010-08-301887. Epub 2010 Sep 24. PMID: 20870901.

18. González-Regueiro JA, Cruz-Contreras M, Merayo-Chalico J, Barrera-Vargas A, Ruiz-Margáin A, Campos-Murguía A, Espin-Nasser M, Martínez-Benítez B, Méndez-Cano VH, Macías-Rodríguez RU. Hepatic manifestations in systemic lupus erythematosus. Lupus. 2020 Jul;29(8):813–824. doi: 10.1177/0961203320923398. Epub 2020 May 9. PMID: 32390496.

19. Grover S, Rastogi A, Singh J, Rajbongshi A, Bihari C. Spectrum of Histomorphologic Findings in Liver in Patients with SLE: A Review. Hepat Res Treat. 2014;2014:562979. doi: 10.1155/2014/562979. Epub 2014 Jul 21. PMID: 25136456; PMCID: PMC4130189.

20. Gunawan M, Her Z, Liu M, Tan SY, Chan XY, Tan WWS, Dharmaraaja S, Fan Y, Ong CB, Loh E, Chang KTE, Tan TC, Chan JKY, Chen Q. A Novel Human Systemic Lupus Erythematosus Model in Humanised Mice. Sci Rep. 2017 Nov 30;7(1):16642. doi: 10.1038/s41598-017-16999-7. PMID: 29192160; PMCID: PMC5709358.

21. Hatachi S, Iwai Y, Kawano S, Morinobu S, Kobayashi M, Koshiba M, Saura R, Kurosaka M, Honjo T, Kumagai S. CD4+ PD-1+ T cells accumulate as unique anergic cells in rheumatoid arthritis synovial fluid. J Rheumatol. 2003 Jul;30(7):1410–9. PMID: 12858435.

22. Huai G, Markmann JF, Deng S, Rickert CG. TGF-β-secreting regulatory B cells: unsung players in immune regulation. Clin Transl Immunology. 2021 Apr 2;10(4):e1270. doi: 10.1002/cti2.1270. PMID: 33815797; PMCID: PMC8017464.

23. Jin C, Gao BB, Zhou WJ, Zhao BJ, Fang X, Yang CL, Wang XH, Xia Q, Liu TT. Hydroxychloroquine attenuates autoimmune hepatitis by suppressing the interaction of GRK2 with PI3K in T lymphocytes. Front Pharmacol. 2022 Sep 15;13:972397. doi: 10.3389/fphar.2022.972397

24. Knippenberg S, Peelen E, Smolders J, Thewissen M, Menheere P, Cohen Tervaert JW, Hupperts R, Damoiseaux J. Reduction in IL-10 producing B cells (Breg) in multiple sclerosis is accompanied by a reduced naïve/memory Breg ratio during a relapse but not in remission. J Neuroimmunol. 2011 Oct 28;239(1-2):80–6. doi: 10.1016/j.jneuroim.2011.08.019. Epub 2011 Sep 21. PMID: 21940055.

25. Lan YW, Choo KB, Chen CM, Hung TH, Chen YB, Hsieh CH, Kuo HP, Chong KY. Hypoxia-preconditioned mesenchymal stem cells attenuate bleomycin-induced pulmonary fibrosis. Stem Cell Res Ther. 2015 May 20;6(1):97. doi: 10.1186/s13287-015-0081-6. PMID: 25986930; PMCID: PMC4487587.

26. Leiss H, Niederreiter B, Bandur T, Schwarzecker B, Blüml S, Steiner G, Ulrich W, Smolen JS, Stummvoll GH. Pristane-induced lupus as a model of human lupus arthritis: evolvement of autoantibodies, internal organ and joint inflammation. Lupus. 2013 Jul;22(8):778–92. doi: 10.1177/0961203313492869. PMID: 23817510.

27. Li C, Jiang P, Wei S, Xu X, Wang J. Regulatory T cells in tumor microenvironment: new mechanisms, potential therapeutic strategies and future prospects. Mol Cancer. 2020 Jul 17;19(1):116. doi: 10.1186/s12943-020-01234-1. PMID: 32680511; PMCID: PMC7367382.

28. Li D, Liu Q, Qi L, Dai X, Liu H, Wang Y. Low levels of TGF-β1 enhance human umbilical cord-derived mesenchymal stem cell fibronectin production and extend survival time in a rat model of lipopolysaccharide-induced acute lung injury. Mol Med Rep. 2016 Aug;14(2):1681–92. doi: 10.3892/mmr.2016.5416. Epub 2016 Jun 21. PMID: 27357811.

29. Lu Z, Chen Y, Dunstan C, Roohani-Esfahani S, Zreiqat H. Priming Adipose Stem Cells with Tumor Necrosis Factor-Alpha Preconditioning Potentiates Their Exosome Efficacy for Bone Regeneration. Tissue Eng Part A. 2017 Nov;23(21-22):1212–1220. doi: 10.1089/ten.tea.2016.0548. Epub 2017 Mar 23. PMID: 28346798.

30. Luo Q, Huang Z, Ye J, Deng Y, Fang L, Li X, Guo Y, Jiang H, Ju B, Huang Q, Li J. PD-L1-expressing neutrophils as a novel indicator to assess disease activity and severity of systemic lupus erythematosus. Arthritis Res Ther. 2016 Feb 11;18:47. doi: 10.1186/s13075-016-0942-0. PMID: 26867643; PMCID: PMC4751645.

31. Nishimura H, Nose M, Hiai H, Minato N, Honjo T. Development of lupus-like autoimmune diseases by disruption of the PD-1 gene encoding an ITIM motif-carrying immunoreceptor. Immunity. 1999 Aug;11(2):141–51. doi: 10.1016/s1074-7613(00)80089-8. PMID: 10485649.

32. Numasawa Y, Kimura T, Miyoshi S, Nishiyama N, Hida N, Tsuji H, Tsuruta H, Segawa K, Ogawa S, Umezawa A. Treatment of human mesenchymal stem cells with angiotensin receptor blocker improved efficiency of cardiomyogenic transdifferentiation and improved cardiac function via angiogenesis. Stem Cells. 2011 Sep;29(9):1405–14. doi: 10.1002/stem.691. PMID: 21755575.

33. Obrișcă B, Vornicu A, Jurubiță R, Achim C, Bobeică R, Andronesi A, Sorohan B, Herlea V, Procop A, Dina C, Ismail G. Corticosteroids are the major contributors to the risk for serious infections in autoimmune disorders with severe renal involvement. Clin Rheumatol. 2021 Aug;40(8):3285–3297. doi: 10.1007/s10067-021-05646-2. Epub 2021 Feb 17. PMID: 33595739.

34. Palm O, Garen T, Berge Enger T, Jensen JL, Lund MB, Aaløkken TM, Gran JT. Clinical pulmonary involvement in primary Sjogren’s syndrome: prevalence, quality of life and mortality--a retrospective study based on registry data. Rheumatology (Oxford). 2013 Jan;52(1):173–9. doi: 10.1093/rheumatology/kes311. Epub 2012 Nov 28. PMID: 23192906.

35. Palmiere C, Tettamanti C, Scarpelli MP, Tse R. The forensic spleen: Morphological, radiological, and toxicological investigations. Forensic Sci Int. 2018 Oct;291:94–99. doi: 10.1016/j.forsciint.2018.08.012. Epub 2018 Aug 18. Erratum in: Forensic Sci Int. 2019 Apr;297:384-387. PMID: 30173072.

36. Pan L, Lu MP, Wang JH, Xu M, Yang SR. Immunological pathogenesis and treatment of systemic lupus erythematosus. World J Pediatr. 2020 Feb;16(1):19–30. doi: 10.1007/s12519-019-00229-3. Epub 2019 Feb 22. PMID: 30796732; PMCID: PMC7040062.

37. Paolino S, Cutolo M, Pizzorni C. Glucocorticoid management in rheumatoid arthritis: morning or night low dose? Reumatologia. 2017;55(4):189–197. doi: 10.5114/reum.2017.69779. Epub 2017 Aug 31. PMID: 29056774; PMCID: PMC5647534.

38. Rathore SS, Isravel M, Vellaisamy S, Chellappan DR, Cheepurupalli L, Raman T, Ramakrishnan J. Exploration of Antifungal and Immunomodulatory Potentials of a Furanone Derivative to Rescue Disseminated Cryptococosis in Mice. Sci Rep. 2017 Nov 13;7(1):15400. doi: 10.1038/s41598-017-15500-8. PMID: 29133871; PMCID: PMC5684196.

39. Rawat S, Dadhwal V, Mohanty S. Dexamethasone priming enhances stemness and immunomodulatory property of tissue-specific human mesenchymal stem cells. BMC Dev Biol. 2021 Nov 4;21(1):16. doi: 10.1186/s12861-021-00246-4. PMID: 34736395; PMCID: PMC8567134.

40. Ray DW, Davis JR, White A, Clark AJ. Glucocorticoid receptor structure and function in glucocorticoid-resistant small cell lung carcinoma cells. Cancer Res. 1996 Jul 15;56(14):3276–80. PMID: 8764121.

41. Sakaguchi S, Sakaguchi N, Asano M, Itoh M, Toda M. Immunologic self-tolerance maintained by activated T cells expressing IL-2 receptor alpha-chains (CD25). Breakdown of a single mechanism of self-tolerance causes various autoimmune diseases. J Immunol. 1995 Aug 1;155(3):1151–64. PMID: 7636184.

42. Sellers AR, Roddy MR, Darville KK, Sanchez-Teppa B, McKinley SD, Sochet AA. Dexamethasone for Pediatric Critical Asthma: A Multicenter Descriptive Study. J Intensive Care Med. 2022 Nov;37(11):1520–1527. doi: 10.1177/08850666221082540. Epub 2022 Mar 3. PMID: 35236174.

43. Song Y, Dou H, Li X, Zhao X, Li Y, Liu D, Ji J, Liu F, Ding L, Ni Y, Hou Y. Exosomal miR-146a Contributes to the Enhanced Therapeutic Efficacy of Interleukin-1β-Primed Mesenchymal Stem Cells Against Sepsis. Stem Cells. 2017 May;35(5):1208–1221. doi: 10.1002/stem.2564. Epub 2017 Feb 5. PMID: 28090688.

44. Sun J, Wei ZZ, Gu X, Zhang JY, Zhang Y, Li J, Wei L. Intranasal delivery of hypoxia-preconditioned bone marrow-derived mesenchymal stem cells enhanced regenerative effects after intracerebral hemorrhagic stroke in mice. Exp Neurol. 2015 Oct;272:78–87. doi: 10.1016/j.expneurol.2015.03.011. Epub 2015 Mar 20. PMID: 25797577.

45. Tarbox JA, Keppel MP, Topcagic N, Mackin C, Ben Abdallah M, Baszis KW, White AJ, French AR, Cooper MA. Elevated double negative T cells in pediatric autoimmunity. J Clin Immunol. 2014 Jul;34(5):594–9. doi: 10.1007/s10875-014-0038-z. Epub 2014 Apr 24. PMID: 24760111; PMCID: PMC4047151.

46. Weening JJ, D’Agati VD, Schwartz MM, Seshan SV, Alpers CE, Appel GB, Balow JE, Bruijn JA, Cook T, Ferrario F, Fogo AB, Ginzler EM, Hebert L, Hill G, Hill P, Jennette JC, Kong NC, Lesavre P, Lockshin M, Looi LM, Makino H, Moura LA, Nagata M; International Society of Nephrology Working Group on the Classification of Lupus Nephritis; Renal Pathology Society Working Group on the Classification of Lupus Nephritis. The classification of glomerulonephritis in systemic lupus erythematosus revisited. Kidney Int. 2004 Feb;65(2):521–30. doi: 10.1111/j.1523-1755.2004.00443.x. Erratum in: Kidney Int. 2004 Mar;65(3):1132. PMID: 14717922.

47. Weiss ARR, Dahlke MH. Immunomodulation by Mesenchymal Stem Cells (MSCs): Mechanisms of Action of Living, Apoptotic, and Dead MSCs. Front Immunol. 2019 Jun 4;10:1191. doi: 10.3389/fimmu.2019.01191. PMID: 31214172; PMCID: PMC6557979.

48. Whelan R, Apfel CC. Pharmacology of postoperative nausea and vomiting. In: Pharmacology and Physiology for Anesthesia. 2013, 503–22. doi: 10.1016/B978-1-4377-1679-5.00029-6.

49. Yang QB, He YL, Peng CM, Qing YF, He Q, Zhou JG. Systemic lupus erythematosus complicated by noncirrhotic portal hypertension: A case report and review of literature. World J Clin Cases. 2018 Nov 6;6(13):688–693. doi: 10.12998/wjcc.v6.i13.688. PMID: 30430127; PMCID: PMC6232573.

50. Yang SH, Park MJ, Yoon IH, Kim SY, Hong SH, Shin JY, Nam HY, Kim YH, Kim B, Park CG. Soluble mediators from mesenchymal stem cells suppress T cell proliferation by inducing IL-10. Exp Mol Med. 2009 May 31;41(5):315–24. doi: 10.3858/emm.2009.41.5.035. PMID: 19307751; PMCID: PMC2701980.

51. Yun Y, Wang X, Xu J, Jin C, Chen J, Wang X, Wang J, Qin L, Yang P. Pristane induced lupus mice as a model for neuropsychiatric lupus (NPSLE). Behav Brain Funct. 2023 Feb 10;19(1):3. doi: 10.1186/s12993-023-00205-y. PMID: 36765366; PMCID: PMC9921421.

52. Zardi EM, Taccone A, Marigliano B, Margiotta DP, Afeltra A. Neuropsychiatric systemic lupus erythematosus: tools for the diagnosis. Autoimmun Rev. 2014 Aug;13(8):831–9. doi: 10.1016/j.autrev.2014.04.002. Epub 2014 Apr 3. PMID: 24704869.

53. Zhang M, Methot D, Poppa V, Fujio Y, Walsh K, Murry CE. Cardiomyocyte grafting for cardiac repair: graft cell death and anti-death strategies. J Mol Cell Cardiol. 2001 May;33(5):907–21. doi: 10.1006/jmcc.2001.1367. PMID: 11343414.

