## Supplementary material for "Autoimmunity and clinical pathology amelioration in SLE by Dexamethasone primed Mesenchymal Stem Cell derived conditioned media": Isolated WJ-MSCs formed a homogenous monolayer of adherent, spindle-shaped cells and exhibited proliferation capacity of WJ-MSCs

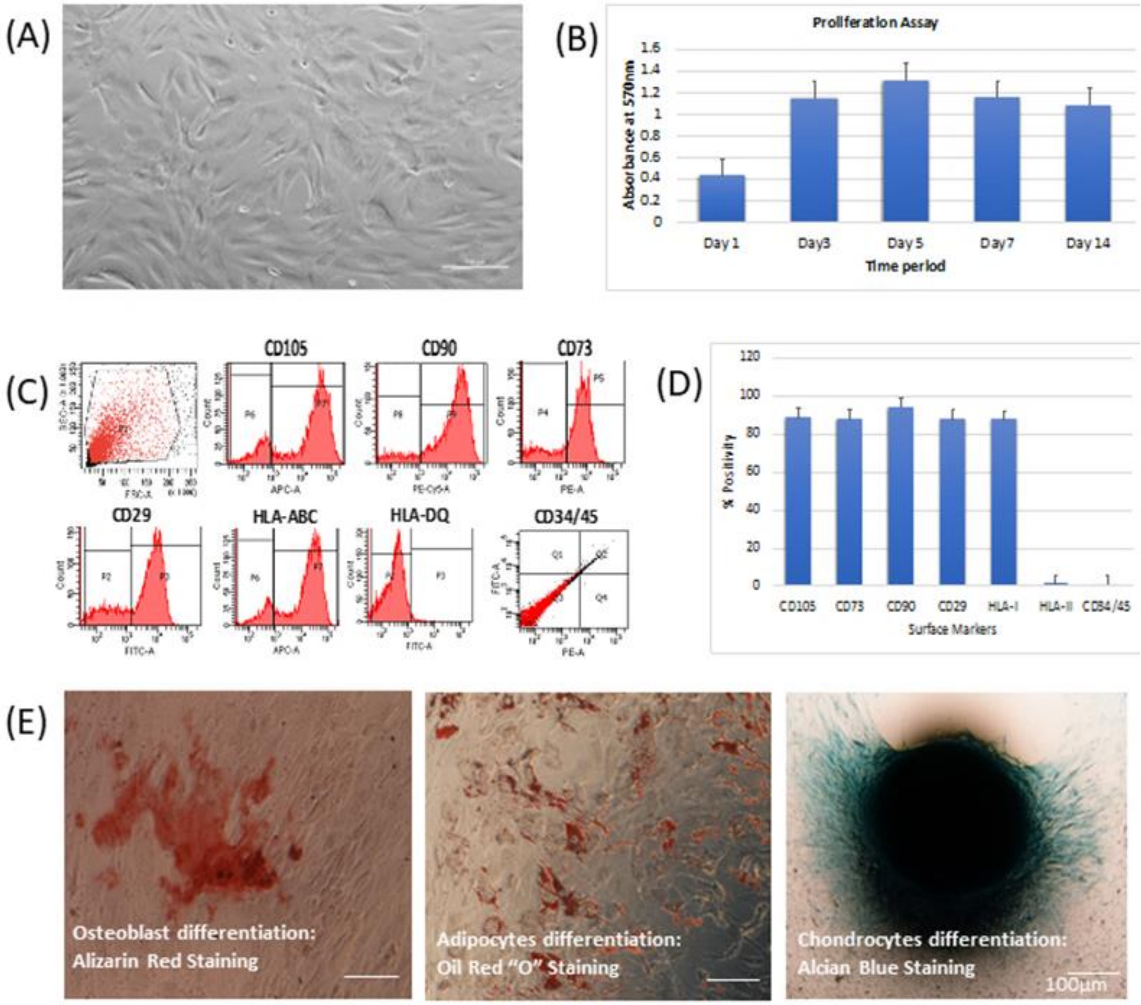


**Figure S1: The representative image shows the basic characterization of WJ-MSCs.** (A) The micrographs shows the cellular typical spindle shaped morphology of WJ-MSCs, (B) the bar graph represents the proliferation rate of WJ-MSCs. (C) FACs histograms shows the marker profile of WJ-MSCs and (D) the bar graph represents the % positivity for WJ-MSCs. (E) The representative micrograph shows the trilineage differentiation capability of WJ-MSCs through alizarin staining for osteoblast, Oil Red "O" staining for adipocytes and Alcian blue staining for chondrocytes.

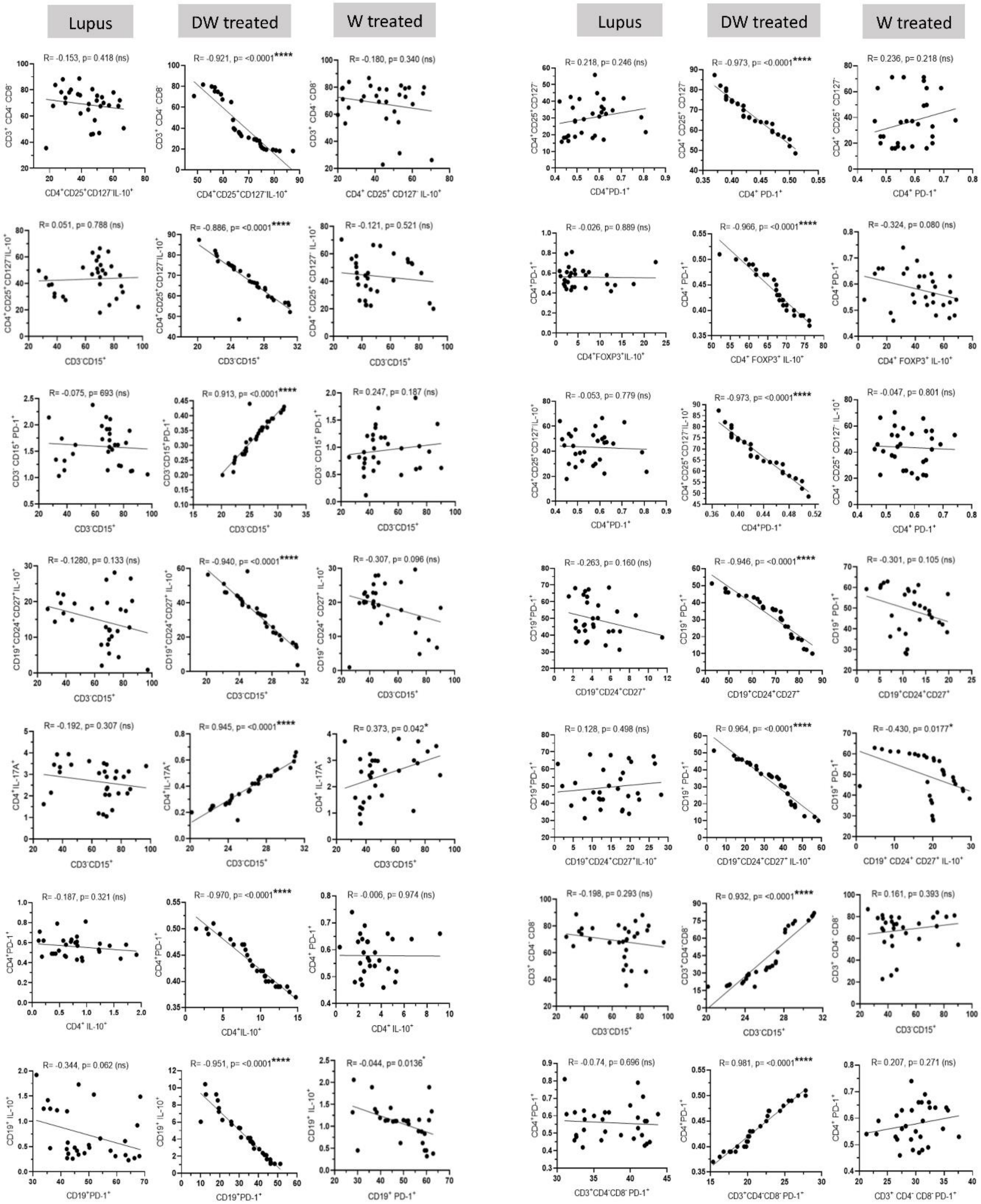

Lupus
DW treated
W treated
R= -0.153, p= 0.418 (ns)
R= -0.921, p= <0.0001****
R= -0.180, p= 0.340 (ns)
CD4+CD25+CD127-IL-10+
CD3+ CD4- CD8-
R= 0.051, p= 0.788 (ns)
R= -0.886, p= <0.0001****
R= -0.121, p= 0.521 (ns)
CD3-CD15+
CD4+CD25+CD127-IL-10+
R= -0.075, p= 693 (ns)
R= 0.913, p= <0.0001****
R= 0.247, p= 0.187 (ns)
CD3- CD15+ PD-1+
R= -0.1280, p= 0.133 (ns)
R= -0.940, p= <0.0001****
R= -0.307, p= 0.096 (ns)
CD19+CD24+CD27+IL-10+
R= -0.192, p= 0.307 (ns)
R= 0.945, p= <0.0001****
R= 0.373, p= 0.042*
CD4+IL-17A+
R= -0.187, p= 0.321 (ns)
R= -0.970, p= <0.0001****
R= -0.006, p= 0.974 (ns)
CD4+ IL-10+
CD4+PD-1+
R= -0.344, p= 0.062 (ns)
R= -0.951, p= <0.0001****
R= -0.044, p= 0.0136*
CD19+PD-1+
CD19+ IL-10+
Lupus
DW treated
W treated
R= 0.218, p= 0.246 (ns)
R= -0.973, p= <0.0001****
R= 0.236, p= 0.218 (ns)
CD4+PD-1+
CD4+CD25+CD127-
R= -0.026, p= 0.889 (ns)
R= -0.966, p= <0.0001****
R= -0.324, p= 0.080 (ns)
CD4+FOXP3+IL-10+
CD4+PD-1+
R= -0.053, p= 0.779 (ns)
R= -0.973, p= <0.0001****
R= -0.047, p= 0.801 (ns)
CD4+PD-1+
CD4+CD25+CD127-IL-10+
R= -0.263, p= 0.160 (ns)
R= -0.946, p= <0.0001****
R= -0.301, p= 0.105 (ns)
CD19+CD24+CD27+
CD19+PD-1+
R= 0.128, p= 0.498 (ns)
R= 0.964, p= <0.0001****
R= -0.430, p= 0.0177*
CD19+CD24+CD27+IL-10+
CD19+PD-1+
R= -0.198, p= 0.293 (ns)
R= 0.932, p= <0.0001****
R= 0.161, p= 0.393 (ns)
CD3-CD15+
CD3+ CD4- CD8-
R= -0.74, p= 0.696 (ns)
R= 0.981, p= <0.0001****
R= 0.207, p= 0.271 (ns)
CD3+CD4-CD8- PD-1+
CD4+PD-1+

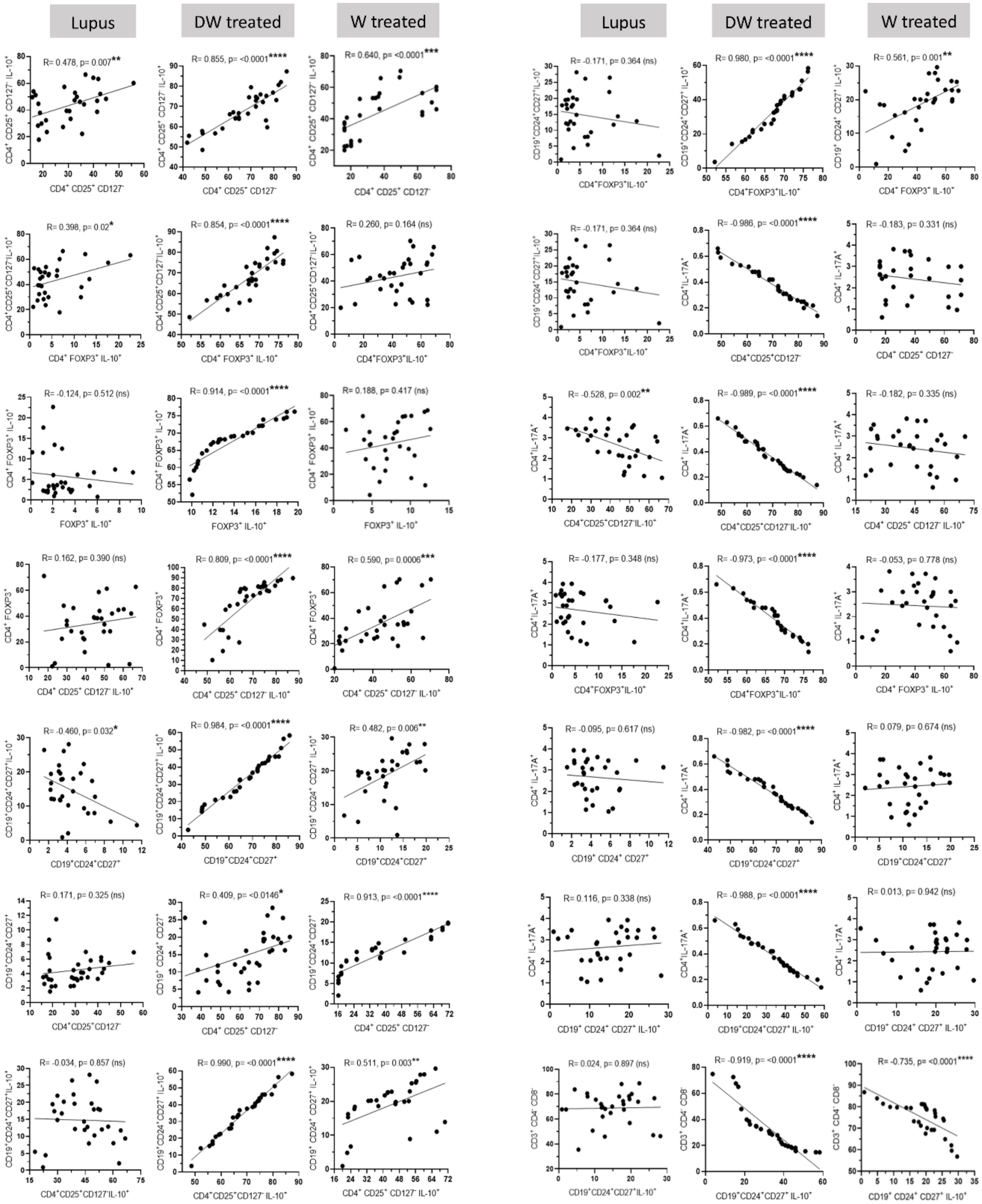
Lupus
DW treated
W treated
Lupus
DW treated
W treated
R= 0.478, p= 0.007**
R= 0.855, p= <0.0001****
R= 0.640, p= <0.0001***
R= -0.171, p= 0.364 (ns)
R= 0.980, p= <0.0001****
R= 0.561, p= 0.001**
R= 0.398, p= 0.02*
R= 0.854, p= <0.0001****
R= 0.260, p= 0.164 (ns)
R= -0.171, p= 0.364 (ns)
R= -0.986, p= <0.0001****
R= -0.183, p= 0.331 (ns)
R= -0.124, p= 0.512 (ns)
R= 0.914, p= <0.0001****
R= 0.188, p= 0.417 (ns)
R= -0.528, p= 0.002**
R= -0.989, p= <0.0001****
R= -0.182, p= 0.335 (ns)
R= 0.162, p= 0.390 (ns)
R= 0.809, p= <0.0001****
R= 0.590, p= 0.0006***
R= -0.177, p= 0.348 (ns)
R= -0.973, p= <0.0001****
R= -0.053, p= 0.778 (ns)
R= -0.460, p= 0.032*
R= 0.984, p= <0.0001****
R= 0.482, p= 0.006**
R= -0.095, p= 0.617 (ns)
R= -0.982, p= <0.0001****
R= 0.079, p= 0.674 (ns)
R= 0.171, p= 0.325 (ns)
R= 0.409, p= <0.0146*
R= 0.913, p= <0.0001****
R= 0.116, p= 0.338 (ns)
R= -0.988, p= <0.0001****
R= 0.013, p= 0.942 (ns)
R= -0.034, p= 0.857 (ns)
R= 0.990, p= <0.0001****
R= 0.511, p= 0.003**
R= 0.024, p= 0.897 (ns)
R= -0.919, p= <0.0001****
R= -0.735, p= <0.0001****

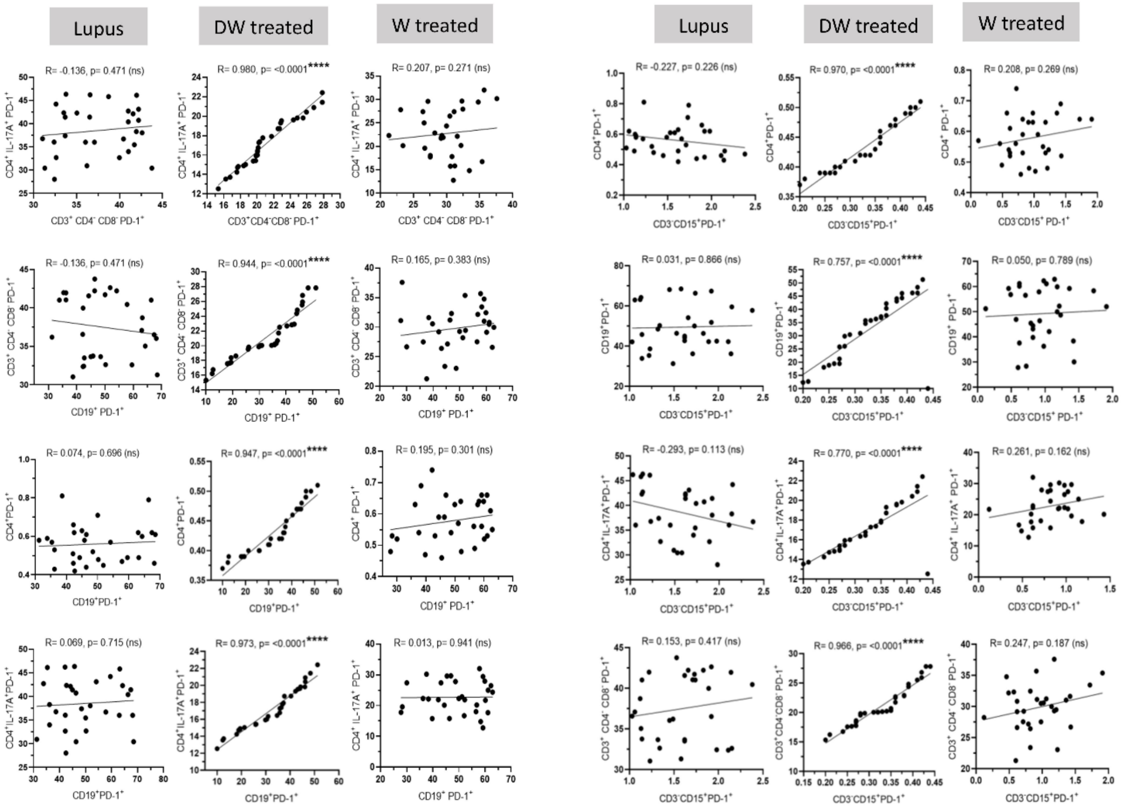


**Figure S2: Correlation between different subsets of Treg, Bregs, Th17, DN T and inflammatory neutrophil population in the SLE patient *in vitro*.** Pearson correlation coefficients 'R' and corresponding p values are indicated. non-significant (ns) $p > 0.05$, *$P < 0.05$, **$p < 0.01$, ***$P < 0.001$ and ****$P < 0.0001$.

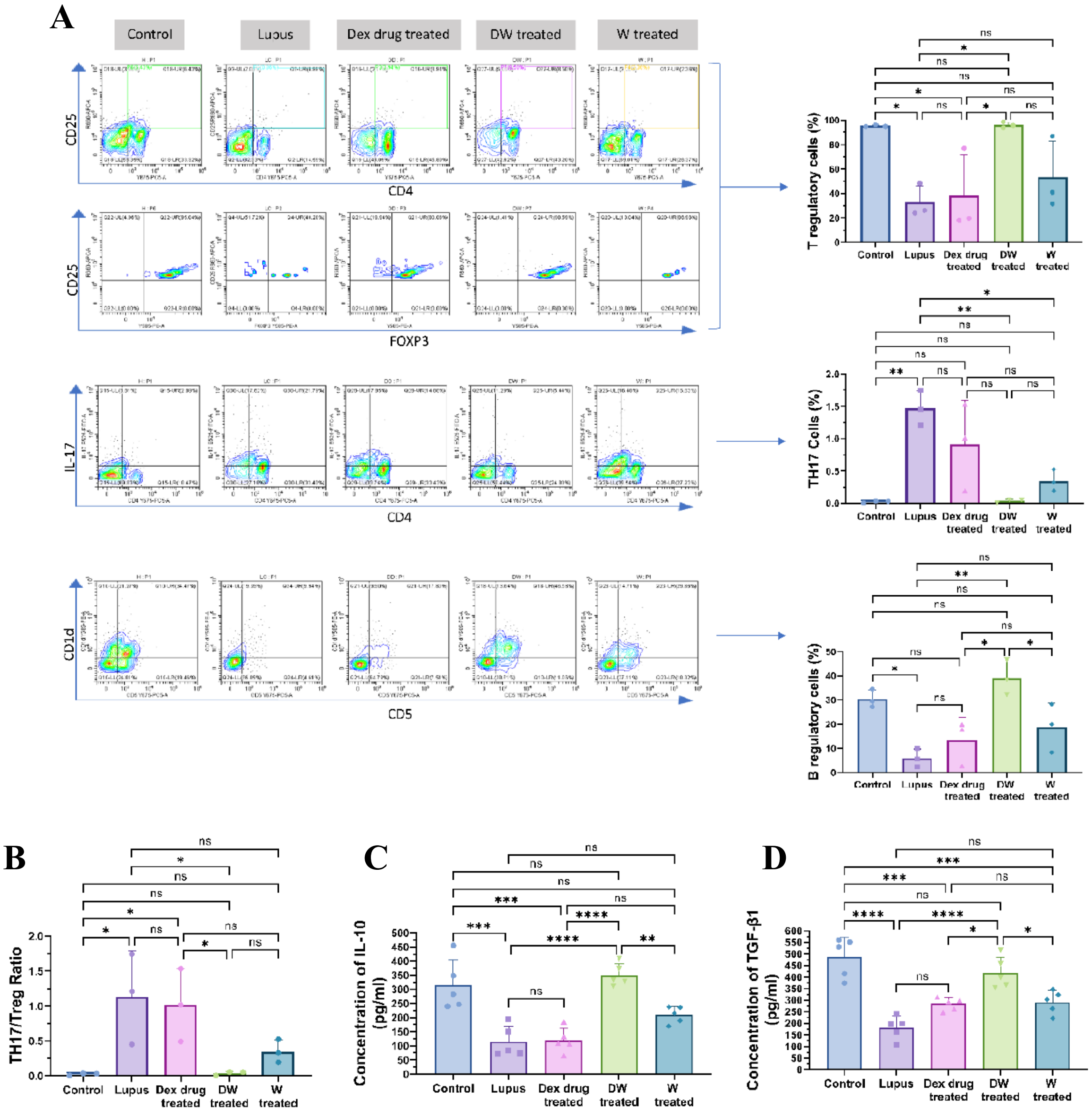


**Figure S3: Suppression of SLE-associated splenic Th17 cells, promotion of regulatory T and B cells, and elevation of anti-inflammatory cytokine IL-10 through TGF-β by DW treatment.** Frequencies of (A) $CD4^{+}$ $CD25^{+}$ $FOXP3^{+}$, $CD4^{+}$ IL-$17A^{+}$ Th17 cells, $CD5^{+}$ CD1d Bregs and (B) Th17/Treg ratio and production of (C) IL-10 and (D) TGF-β in serum of from control (healthy, normal saline treated), lupus (normal saline treated), Dex drug, DW and W treated mice (n = 3) for one month were compared with each other. The contour plots depict findings from an individual sample of mice, and corresponding bar graphs are presented with a sample size of 3. Error bars show mean ± SD. p values indicate significant changes as follows: non-significant (ns) $p > 0.05$, *$P < 0.05$, **$p < 0.01$, ***$P < 0.001$ and ****$P < 0.0001$; One-way ANOVA.

**Video 1 and 2: Development of Seizure in PIL mice**

**Video 3: Seizure resolved with DW treatment**
